# Development of Clinically Viable Non-Muscle Myosin II Small Molecule Inhibitors with Broad Therapeutic Potential

**DOI:** 10.1101/2024.10.07.617018

**Authors:** Laszlo Radnai, Erica J. Young, Carlos Kikuti, Madalyn Hafenbreidel, Rebecca F. Stremel, Li Lin, Katalin Toth, Paolo Pasetto, Xiaomin Jin, Aagam Patel, Michael Conlon, Sherri Briggs, Leïla Heidsieck, H. Lee Sweeney, James Sellers, Teresa Krieger-Burke, William H. Martin, Jay Sisco, Steven Young, Paul Pearson, Gavin Rumbaugh, Gian Luca Araldi, Steven K. Duddy, Michael D. Cameron, Matthew Surman, Anne Houdusse, Patrick R. Griffin, Theodore M. Kamenecka, Courtney A. Miller

## Abstract

Non-muscle myosin II (NMII), a molecular motor that regulates critical processes such as cytokinesis and neuronal synaptic plasticity, has substantial therapeutic potential. However, translating this potential to *in vivo* use has been hampered by the lack of selective tools. The most prototypical non-selective inhibitor, blebbistatin inactivates both NMII and cardiac myosin II (CMII), a key regulator of heart function. Using rational drug design, we developed a series of NMII inhibitors that improve tolerability by selectively targeting NMII over CMII, including MT-228, which has excellent properties such as high brain penetration and efficacy in preclinical models of stimulant use disorder, which has no current FDA-approved therapies. The structure of MT-228 bound to myosin II provides insight into its 17-fold selectivity for NMII over CMII. MT-228’s broad therapeutic window opens the door to new disease treatments and provides valuable tools for the scientific community, along with promising leads for future medication development.

**Highlights:**

- Research suggests numerous indications, from axon regeneration and cancer, would benefit from a small molecule inhibitor of non-muscle myosin II, a molecular motor that regulates the actin cytoskeleton.
- Current chemical probe options are very limited and lack sufficient safety for *in vivo*
- studies, which we show is primarily due to potent inhibition of cardiac myosin II.
- Rational design that focused on improving target selectivity over the pan-myosin II inhibitor, blebbistatin, led to the identification of MT-228, a small molecule inhibitor with a wide therapeutic window.
- High-resolution structure of MT-228 bound to myosin II reveals that selectivity results from a different positioning compared to blebbistatin and an important sequence difference between cardiac and non-muscle myosin II in the inhibitor binding pocket.
- A single administration of MT-228 shows long-lasting efficacy in animal models of stimulant use disorder, a current unmet and rapidly escalating need with no FDA-approved treatments.

## INTRODUCTION

Myosin proteins are a largely untapped superfamily of potential drug targets for new therapeutics. Upon interaction with filamentous actin (actomyosin), these actin-based molecular nanomotors produce mechanical force by utilizing the chemical energy stored in ATP (*1, 2*). They are kinetically fine-tuned and structurally adapted to achieve a wide range of cellular functions, including the transport of organelles along actin filaments, tethering of actin to various cellular compartments, and the sliding motion of actin and myosin filaments relative to each other during muscle contraction. The general mechanochemical cycle of myosins starts with ATP binding to their motor domain, which results in the shift of the otherwise strong binding between myosin and actin towards a weakly bound state. In this state, dissociation from actin occurs with a higher probability. Following ATP hydrolysis and re-binding to actin, the release of the hydrolysis products (phosphate and ADP) coupled to conformational changes in the motor domain eventually result in the displacement of myosin relative to the actin filament. Members of the myosin superfamily show kinetic differences in this general mechanochemical cycle, changing the way they interact with actin (e.g. the fraction of time spent strongly bound to actin). This kinetic adaptation, together with their individual structural features and temporospatial patterns of expression, determine the physiological roles of different myosin motors *in vivo* (*1, 2*). Across 340 eukaryotic species, 45 different myosin classes have been identified (*3*), with 12 classes in humans bearing unique distributions and functions.

One class of particular interest is the myosin II family, which generates the mechanical force to achieve processes that include organelle transport, endocytosis, chemotaxis, cytokinesis, muscle contraction and neuroplasticity (*4*). Myosin II achieves this through differing expression patterns of four subfamilies: skeletal muscle myosin II (SkMII), cardiac muscle myosin II (CMII), smooth muscle myosin II (SmMII) and non-muscle myosin II (NMII) (*1*). Myosin IIs have two heavy chains and several light chains. The heavy chains bear the force generating motor heads and are what distinguish the differential expression and functions of the myosin II subfamilies and their paralogs throughout the body (*1*).

Two NMII paralogs, IIA and IIB, have been implicated in a wide range of disease states, including, but not limited to, microbial infection, cancer, and neuronal regeneration (*5-22*). In addition, we have identified an unexpected role for NMIIB in the brain pointing to its potential therapeutic application to methamphetamine (METH) use disorder (MUD) (*23, 24*). MUD currently has no FDA-approved treatments and the pipeline for potential new therapeutics is very limited (*25, 26*). Relapse rates among individuals diagnosed with MUD are extremely high (*27*) and overdose deaths are increasing at an alarming rate (*28*). MUD is marked by the sustained motivation to seek and consume METH in spite of adverse consequences. Motivated behavior is supported and maintained by structural neuroplasticity within dendritic spines, postsynaptic structures on neurons that contribute to information storage (*29*). We have previously demonstrated that structural neuroplasticity is driven by NMIIB’s ability to drive actin dynamics within dendritic spines and these synaptic cytoskeletal dynamics are subsequently maintained by rapid stabilization (*30*). However, we discovered that METH prevents actomyosin dynamics from stabilizing in dendritic spines of the basolateral amygdala (BLA) (*31*), a subregion of the brain that regulates emotional memory and motivation (*32*). The uniquely sustained actomyosin dynamics that result from METH exposure render the structural neuroplasticity changes supporting the motivation for METH selectively vulnerable to NMIIB genetic or pharmacologic interference long after drug exposure, without altering the actin cytoskeleton and motivation supporting other behaviors (*33*). Indeed, in preclinical models of METH seeking, we discovered that blocking NMII reverses METH-associated synaptic changes in the BLA to levels seen in neurons never exposed to METH. Further, NMII inhibition produces an immediate, long-lasting, and selective disruption of the motivation for METH with a single administration (*23, 31*). This introduces NMII as a new therapeutic target for the treatment of MUD that bears a unique benefit of potentially only requiring a single administration for efficacy (*23, 24, 34-36*). This is desirable, as daily administration of medication can be challenging in this patient population.

The potential of myosin IIs as therapeutic targets has received limited attention until recently. However, there are now multiple programs focused on the development of SkMII (*37, 38*) and CMII inhibitors. The CMII inhibitor mavacamten (Camyzos®) was recently approved by the FDA for hypertrophic cardiomyopathy (*39, 40*). CMII is arguably one of the most challenging myosins to target because of its fundamental importance to cardiomyocyte contractility and, therefore, heart function. Mavacamten safety was achieved through patient-specific pharmacokinetics and careful dose escalation combined with echocardiogram (echo) measures (<u>NCT03442764</u>) (*41*). Unfortunately, current options for targeting NMII are largely limited to blebbistatin (Blebb) (*42*). While Blebb does not inhibit other myosin superfamilies (*43, 44*), the small molecule and its currently available derivatives, such as nitro- and aminoblebbistatin (*45, 46*), do inhibit both the ATPase and force generation of all three muscle myosin II subclasses (*43, 44, 47, 48*), making them pan-myosin II inhibitors. This introduces significant tolerability issues.

Given the myriad of indications with unmet needs for which NMII has been implicated, there is clear value for safe and selective NMII inhibitors (*49*). Here we report the results of a medicinal chemistry campaign to improve the selectivity and drug-like properties of the tool compound, Blebb, and initiate therapeutic development of a first-in-class NMII small molecule inhibitor. MT-228 bears improved CNS penetrance, solubility, photostability and *in vivo* tolerability through increased selectivity. High resolution structural data with MT-228, reveals the molecular basis for the selectivity of NMII versus CMII. We also demonstrate MT-228’s efficacy and behavioral selectivity in preclinical models of MUD.

## RESULTS

### Blebbistatin has Poor Tolerability Due to Cardiotoxicity

To explore Blebb’s therapeutic potential, we first determined its solubility and oral bioavailability to establish a clinically feasible route of administration. Blebb is chiral and we determined the aqueous solubility of its active enantiomer to be low (21 µM). When animals were administered racemic Blebb, the parent compound could not be detected in plasma following PO administration (C_max_ < 8.77 ng/mL). Therefore, we examined the pharmacokinetics, brain penetrance and tolerability of intravenously (IV) administered Blebb. IV infusion of the active enantiomer of Blebb (which was used for all future experiments) resulted in excellent brain penetrance (B:P = 2.2 at 30 minutes, **Table 1**, top line). It was detected at concentrations of 65.2 ± 7.6 ng/mL at 30 minutes after a 0.5 mg/kg IV bolus infusion (t_½_ = 3.3 hours; **Table 1**). While uncommon in drug development, Blebb’s short half-life was considered desirable, as we have previously shown that NMII inhibition achieves near immediate and long-lasting effects on METH seeking in preclinical models (*23, 24, 35*), indicating sustained or repeated exposures are likely not necessary in a therapeutic for MUD. Further, a short half-life would presumably mitigate potential on-target, but unwanted effects of systemic NMII inhibition.

**Table 1.** NMII inhibitor pharmacokinetic properties.

| Compound | Species | Dose<br>(mg/kg) | Route of<br>Admin | ½ Life<br>(hr) | C<sub>max</sub><br>(ng/mL) | AUC<sub>last</sub><br>(hr*ng/mL) | B:P<br>(30min) |
| --- | --- | --- | --- | --- | --- | --- | --- |
| Blebbistatin | Rat | 0.5 | IV | 3.3 | 65.2<br>± 7.8 | 53.7<br>± 5.2 | 2.2 |
| MT-106 | Rat | 0.5 | IV | 1.5 | 136.5<br>± 21.5 | 94.7<br>± 11.3 | 3.5 |
| MT-150 | Rat | 0.5 | IV | 1.3 | 457.0<br>± 18.0 | 203.0<br>± 0.0 | 1.5 |
| MT-151 | Rat | 0.5 | IV | 1.6 | 548.5<br>± 27.5 | 194.0<br>± 4.0 | 1.6 |
| MT-152 | Rat | 0.5 | IV | 0.25 | 69.1<br>± 6.7 | 45.1<br>± 5.3 | 8.3 |
| MT-154 | Rat | 0.5 | IV | 1.6 | 133.5<br>± 20.5 | 98.0<br>± 24.1 | 7.1 |
| MT-160 | Rat | 0.5 | IV | 3.4 | 74.3<br>± 19.1 | 60.5<br>± 14.7 | 5.1 |
| MT-228 | Rat | 0.5 | IV | 1.8 | 91.9<br>± 12.1 | 57.0<br>± 12.8 | 5.0 |
| MT-228 | Rat | 2 | IV | 3.8 | 533.5 ±<br>73.5 | 289.0<br>± 35.0 | 5.0 |
| MT-228 | Mouse | 2 | IP | 1.8 | 41.5<br>± 4.3 | 52.0<br>± 3.2 | 2.9 |

Values represented are means ± SEM

Unfortunately, tolerability issues prevented evaluation of higher doses. To understand the source of Blebb’s poor tolerance, we profiled its inhibitory activity at each of the four myosin II’s using a steady state ATPase assay (*50*). As we and others have previously reported (*38*) and shown in **Table 2** and **Table S1**, Blebb is a potent inhibitor of SkMII (K_I_ = 0.28 µM), with slightly less inhibition of NMIIA (K_I_ = 2.9 µM), CMII (K_I_ = 1.9 µM), and SmMII (K_I_ = 3.2 µM). We used a COS7 cell-based cytokinesis assay (*51*) to determine inhibition of our primary target of interest, NMIIB. COS7 cells were selected because their cytokinesis depends exclusively on NMIIB, as NMIIA and NMIIC are not expressed (*52*). Therefore, the half maximal effective concentration (EC_50_) determined in this assay reflects the compounds’ potency for inhibition of NMIIB activity. The NMIIB EC_50_ of Blebb was 1.9 µM. Together, this demonstrated that Blebb has similar potency at NMIIA and IIB, but also CMII.

**Table 2.**
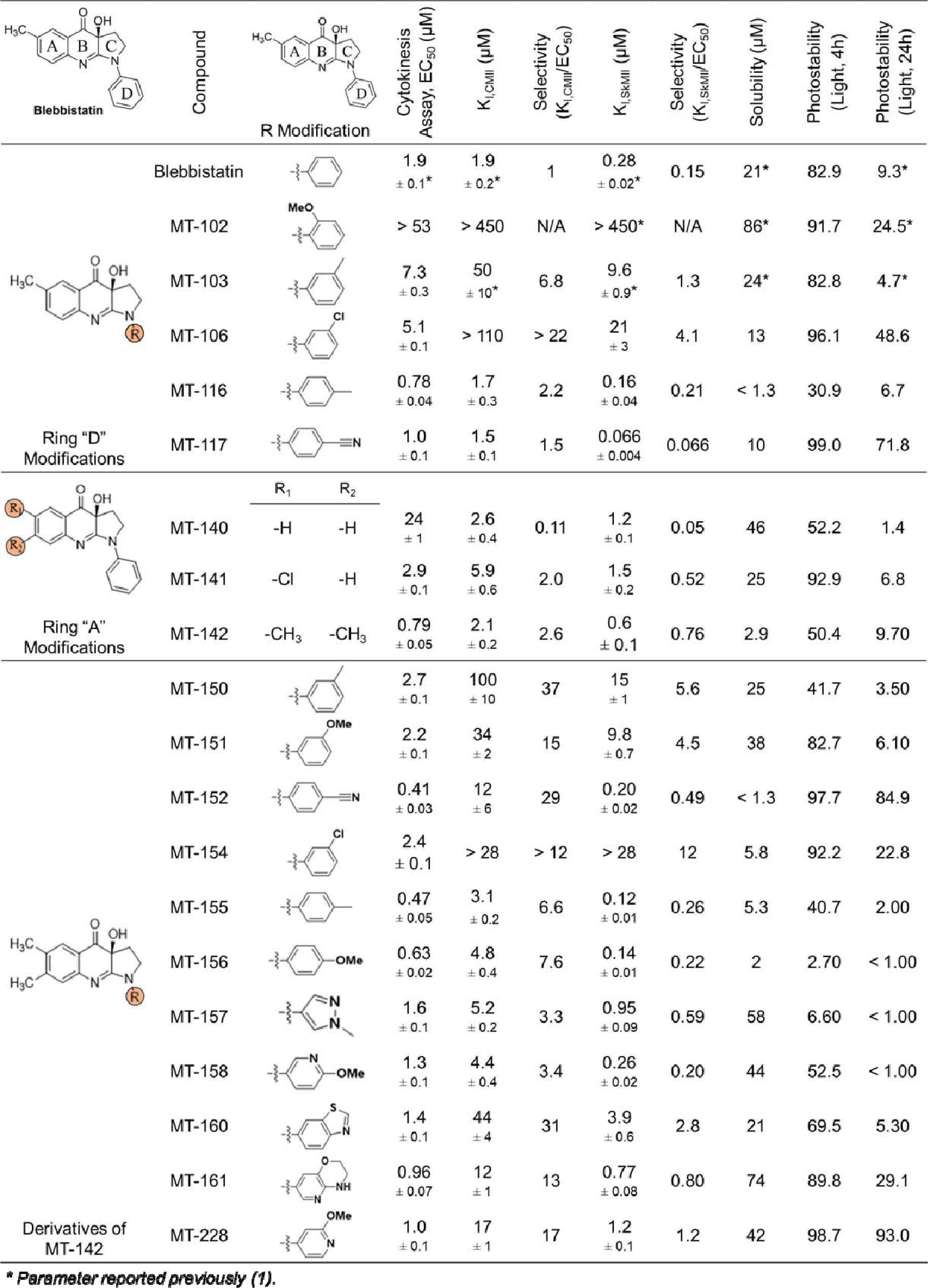
Structure-activity relationships (SAR) of modifications to Blebb at the A and D rings.

Given the critical role of CMII in driving cardiomyocyte contractility (*53*) and, therefore, heart function, we hypothesized that CMII inhibition was the greatest liability (*38, 54*). To test this, we performed an *in vitro* measure of Blebb’s effects on cardiac function, assessing the amplitude of spontaneous contractility in human induced pluripotent stem cell (hiPSC)-derived Cor.4U cardiomyocytes as measured by impedance analysis (**Fig. 1A**). Blebb significantly reduced cardiomyocyte contractility across all four concentrations tested, with a 94% reduction at 3 µM (**Fig. 1B-C**), essentially flatlining contractility (Cell Index Changes from DMSO control: *P* < 0.05 at all doses tested; **Fig. 1B-C**). To determine if this dramatic reduction in cardiomyocyte contractility translated to *in vivo* effects, we next tested Blebb in an echo assay in rats (**Fig. 1D**). Following IV infusion of vehicle, there was no change in cardiac output, a combined measure of heart rate and volume output, over the 10 minutes of observation (cardiac output, Baseline x Veh Start, *P* > 0.05; Baseline x Veh, + 5min *P* > 0.05; Baseline x Veh + 10min, *P* > 0.05, **Fig. 1D**) or when vehicle was infused three times across the full hour-long test period (**Fig. 1D**). However, Blebb treatment significantly decreased cardiac output, (rmANOVA F_(4, 9)_ = 27.79, *P* < 0.0001; **Fig. 1D)**. A single infusion of Blebb at 0.5 mg/kg (IV) led to a 25-50% reduction in contractility that was sustained for the 10 minute imaging period (*P* < 0.0001 at D1 start x D1 + 5 min and + 10 min), followed by a further reduction (-80% from baseline) with a second 0.5 mg/kg infusion of Blebb (D2 start x D2 + 5min *P* < 0.001; D2 start x D2 + 10min *P* < 0.01; **Fig. 1D**). A higher second infusion of Blebb (2 mg/kg, IV) to a single animal was not tolerated (**Fig. 1D**).

**Figure 1.**
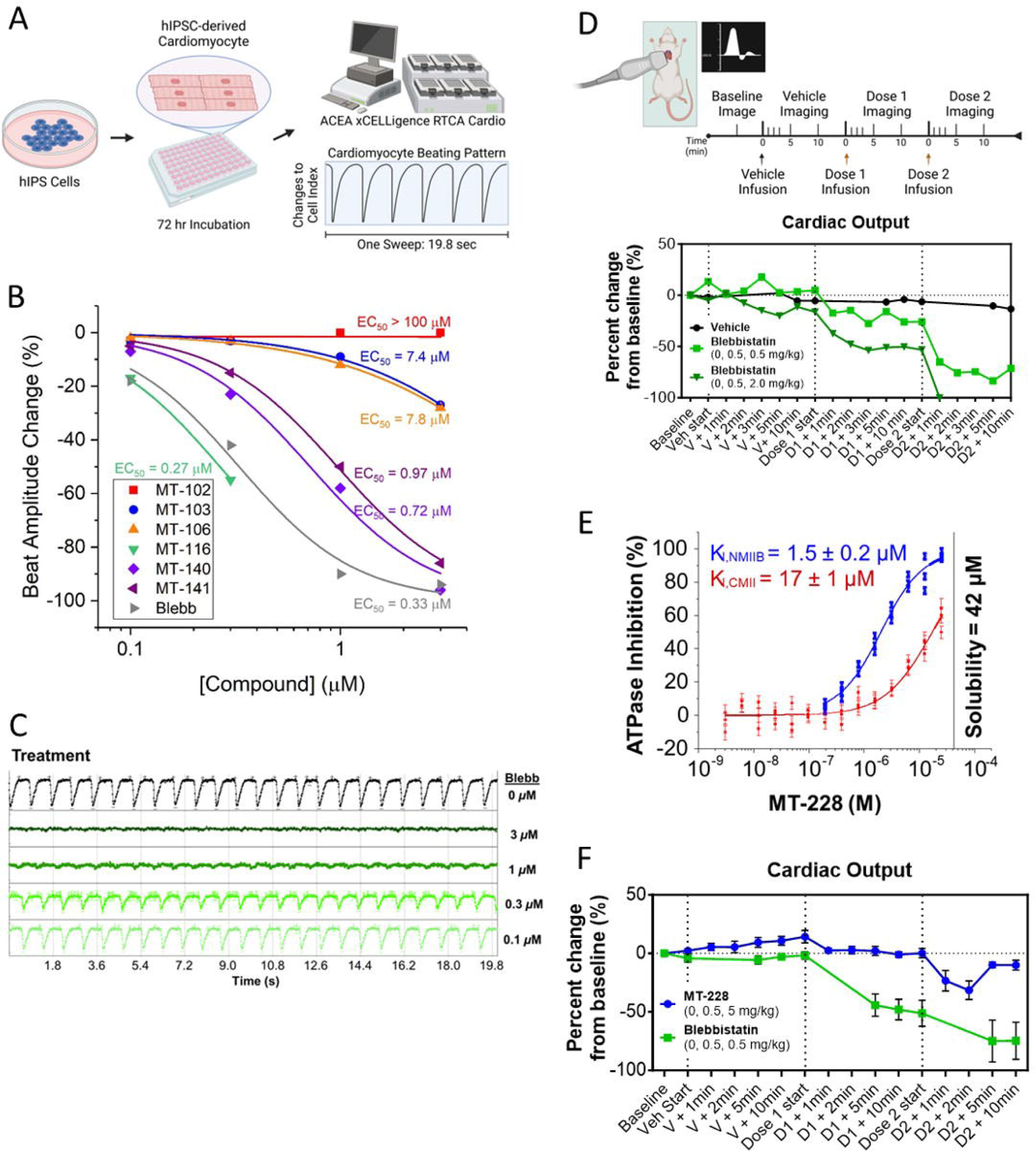
MT-228 has improved cardiac safety and selectivity compared to Blebb. A) Experimental timeline for measuring spontaneous beating cycle in hIPSC-derived cardiomyocytes before and after 30 min of exposure to Blebb. B) All but one of the novel compounds (MT-116) had less impact on beat amplitude than Blebb (grey line) across all concentrations tested. Potency (EC_50_) was estimated for each compound by fitting the Hill-equation to the dose-response data assuming that the minimum and maximum asymptotes were equal to 0% and -100% beat amplitude changes, respectively, in all cases. C) Treatment with 0.3 μM Blebb decreased beat amplitude by greater than 40% and at higher doses (1 and 3 μM) cardiomyocyte beating was almost completely disrupted. D) Cardiac output was measured via echo in rats following treatment with Veh and 2 infusions of Veh or Blebb. Doses as low as 0.5 mg/kg Blebb significantly decreased cardiac output and 2 mg/kg was lethal. E) MT-228 has improved selectivity and solubility. In an ATPase assay, the K_I,NMIIB_ for MT-228 was 1.5 µM, which indicates a 11.3-fold selectivity for NMIIB over CMII. F) Unlike Blebb, MT-228 had no significant effect on cardiac output measured via echo in rats, even when the dose was increased 10-fold to 5 mg/kg. Error bars represent SEM, * *P* < 0.05.

Given the significant negative cardiac effects of Blebb, the primary goals of our SAR strategy were to generate analogs with improved selectivity by reducing CMII inhibition, while maintaining or improving NMII inhibition and brain penetrance. Prior reports have indicated that Blebb is light sensitive *in vitro* and can be phototoxic in living cell studies (*55-58*). When maintained in the dark, we found that samples continued to contain > 99% Blebb after 24 hours in aqueous solution. However, Blebb displayed poor photostability with exposure to bright light. Levels were reduced to 82.9% after 4 hours of light exposure, followed by a dramatic reduction to just 9.3% by 24 hours (**Table 2**). Furthermore, Blebb is known to form precipitates under various experimental conditions both *in vitro* and *in vivo* due to its relatively poor solubility (*45, 59*). Therefore, photostability and kinetic aqueous solubility were determined for all compounds. The latter helped us to not only identify compounds with improved solubility, but also to avoid reporting data affected by artifacts related to compound precipitation.

### Synthesis, Purification and Characterization of Blebbistatin Derivatives

In total, close to 500 Blebb derivatives targeting the A and D rings were made and characterized in our medicinal chemistry campaign. Representative examples are detailed in the Supplemental Results section with detailed descriptions of synthetic routes, characterization of compounds, proof of purity, ADME and preliminary safety data. Ultimately a combination of A- and D-ring modifications were found to produce the largest improvement on solubility, potency, selectivity and cardiac safety compared to Blebb (**Table 2**). Ortho-substituted D-ring analogues were generally found to be suboptimal myosin II inhibitors, though the introduction of para-substituents slightly increased potency and other favorable properties (e.g., improved photostability). In general, meta-substituents significantly increased selectivity over CMII, but decreased NMIIB potency. To further explore modifications that enhanced selectivity, while increasing NMIIB potency, Blebb derivatives with ring “A” modifications were synthesized, as shown in **Table 2**. Although solubility was reduced, dimethyl-“A” ring modifications appeared to be the best option to simultaneously improve potency and selectivity, as evident with **MT-142**.

To test the effectiveness of combining D- and A-ring modifications, several ring “D” modified derivatives of **MT-142** were next synthesized (summarized in **Table 2**). Two of these compounds, **MT-158** and **MT-228**, were synthesized with a 2-methoxypyridine “D” ring. In these derivatives, the methoxy-group is in the para- or meta-position, respectively, and the only difference between **MT-158** and **MT-156** is the presence of the nitrogen atom in ring “D”. The same difference exists between **MT-228** and **MT-151**. Compared to **MT-156**, **MT-158** showed a ∼2-fold decrease in the selectivity for NMIIB over CMII as a result of the decreased potency on NMIIB. As expected, the solubility was improved. However, the photostability of **MT-158** remained poor, as did the selectivity for NMIIB over SkMII due to potent SkMII inhibition (K_I_ = 0.26 µM; **Table 2, Figs. S1P, S2P**). Compared to **MT-151**, the nitrogen in **MT-228** resulted in a 2-fold increase in the potency on NMIIB, providing excellent selectivity over CMII (∼17-fold improvement over Blebb). Interestingly, **MT-228**’s solubility was double that of Blebb at 42 µM and there was a drastic increase in the photostability, with 93% of the compound remaining after exposure to bright light for 24 hours. **MT-228**’s selectivity for NMIIB over SkMII also showed an 8-fold improvement over Blebb (**Table 2, Figs. S1S, S2S**).

To confirm that the EC_50_ determined in the cytokinesis assay reflects the inhibitory potential (K_I,NMIIB_) of the compounds on NMIIB, we also determined K_I,NMIIB_ for **MT-228** in an NMIIB ATPase assay. The K_I,NMIIB_ (1.5 µM, **Fig. 1E**) showed excellent agreement with the cytokinesis EC_50_ (1.0 µM, **Table 2** and **Fig. S1S**), providing >11-fold selectivity over CMII. The inhibitory effect on NMIIA and SmMII of a subset of compounds chosen for their selectivity of >10-fold for NMIIB over CMII was also determined in the ATPase assay (**Table S1**). The K_I,NMIIA_ values showed good correlation (Pearson correlation coefficient = 0.96) with the half maximal effective concentrations representing NMIIB inhibition (**Tables 2 and S1**), with the K_I,NMIIA_ typically slightly higher than the EC_50_ determined in the cytokinesis assay. Both parameters had similar values in the micromolar range. While the experimental conditions in an ATPase assay and in the cell-based cytokinesis assay are different, the similar results for NMIIA and NMIIB potency likely reflect the high degree of similarity between the two proteins. **MT-106, MT-150**, and **MT-152** are slightly more biased towards inhibition of NMIIB than NMIIA (NMIIB/IIA ratios = 0.26, 0.29, and 0.18, respectively), as compared to Blebb (NMIIB/IIA ratio = 0.66). An improvement in NMIIB selectivity over SmMII, similar to the improved NMIIB to CMII ratio, was observed in all cases. Also, the K_I,SmMII_ and K_I,NMIIA_ values were highly correlated, with the K_I,SmMII_ typically being higher.

Profiling of **MT-228** against a panel of cytochrome P450s resulted in inhibition greater than 50% at CYPs 2D6, 3A4, and 2C9 (**Table S2**). **MT-228** was also rapidly metabolized in the four species of liver microsomes tested (**Table S3**). It was determined to not be a P-gp substrate and, like Blebb, had high PPB (>90%; **Table S2**). Given **MT-228**’s promising profile, additional ADMET studies were performed. It was found to be negative in an Ames genetic toxicity assay (Salmonella typhimurium reverse mutation assay) as compared to positive controls and did not induce activation of the nuclear receptor PXR (*NR1I2*), as determined in a promoter-reporter gene assay, removing the concern of elevated PXR-induced drug metabolism. At a concentration of 10 µM, **MT-228** demonstrated no inhibition of hERG, less than 10% inhibition of Cav1.2 ion channels, and less than 18% inhibition of Nav1.5 and was negative in the hERG-CHO patch clamp assay, indicating that cardiac channel inhibition is not a concern. **MT-228** was also evaluated at 10 µM in an *in vitro* safety pharmacology binding assay panel of 55 receptors and ion channels. Inhibition of greater than 50% was only observed in assays for the adenosine receptor A3 (agonist radioligand) at 51.5% and the serotonin receptor 5-HT_2B_ (agonist radioligand) at 59.2%, demonstrating acceptable selectivity.

**MT-228**’s brain penetrance was also significantly improved over Blebb. The B:P was more than double (0.5 mg/kg, IV, B:P 30 min post-dose: MT-228 = 5.0, Blebb = 2.2; **Table 1**) and absolute brain concentrations achieved were increased (C_max_ MT-228 = 91.9 ± 12.1 ng/mL, Blebb 65.2 ± 7.6 ng/mL; AUC_last_ MT-228 = 57.0 ± 12.8 ng/mL.hr, Blebb = 53.7 ± 5.2 ng/mL.hr; **Table 1**). As noted above, a short half-life is desirable for a potential MUD therapeutic to balance efficacy and tolerability. To this end, **MT-228** was also considered improved in terms of half-life (t_1/2_ = 1.8 hours) over Blebb (t_1/2_ = 3.3 hours).

Consistent with its improvement in selectivity for NMIIB over CMII, **MT-228** had no effect on cardiac output in an echo study at 0.5 mg/kg (IV; **Fig. 1F**, overall rmANOVA F_(4, 15)_ = 7.2, *P* < 0.001; Baseline x D1 + 1min: *P* > 0.05; Baseline x D1 + 2min: *P* > 0.05; Baseline x D1 + 5min: *P >* 0.05; Baseline x D1 + 10min: *P* > 0.05; all comparisons between Veh Start or D1 Start and all D1 timepoints *P* > 0.05), and only a minor impact that fully recovered in less than 5 minutes at 10 times the dose (5.0 mg/kg + previous 0.5 mg/kg infusion; Baseline x D2 + 1min: *P* < 0.05; Baseline x D2 + 2min: *P* < 0.05; Baseline x D2 + 5min: *P* > 0.05; Baseline x D2 + 10min: *P* > 0.05). Further, heart rate was also normal at all doses tested (overall rmANOVA F_(4, 15)_ = 0.93, *P >* 0.05). Taken together, these data indicate that **MT-228** successfully addressed the cardiotoxicity of Blebb.

All properties of interest for our SAR strategy were greatly improved in **MT-228** over Blebb, including selectivity for NMIIB over CMII, high brain penetrance with a reduced half-life in plasma, and better solubility and photostability. An additional measure incorporated into our cytokinesis assay is an assessment of potential cytotoxicity (*51*). With the exception of **MT-157**, which showed signs of mild cytotoxicity, all analogs reported in this study, including **MT-228**, were not cytotoxic in the concentration range tested (**Fig. S1**). And, of course, due to the nature of the cell-based cytokinesis assay, we can conclude that **MT-228** is cell membrane permeable.

### MT-228 Metabolite Identification

For some disease indications, a more stable compound with a longer half-life may be desirable. To support future efforts to that end, putative metabolites of **MT-228** were determined in rat. Metabolites generated by incubating **MT-228** with rat hepatic microsomes elucidated three pathways that impact the stability of the molecule, comprising hydroxylation, reduction, and demethylation (**Fig. 2**). Hydroxylated metabolites were the most abundant detected in the *in vitro* hepatic microsomal incubations. Plasma collected from rats 30 minutes after administration of **MT-228** (10 mg/kg, IP) had the same metabolites observed in the *in vitro* incubation. Rat plasma samples contained a higher ratio of the reduced metabolite and the reduced plus hydroxylated secondary metabolites. This difference is likely due to extra-hepatic tissues contributing to the reduction of **MT-228**. Based upon the metabolite data, eliminating the sites of reduction and exploring replacements for the methyl substituents would be expected to improve the pharmacokinetics of **MT-228**.

**Figure 2.**
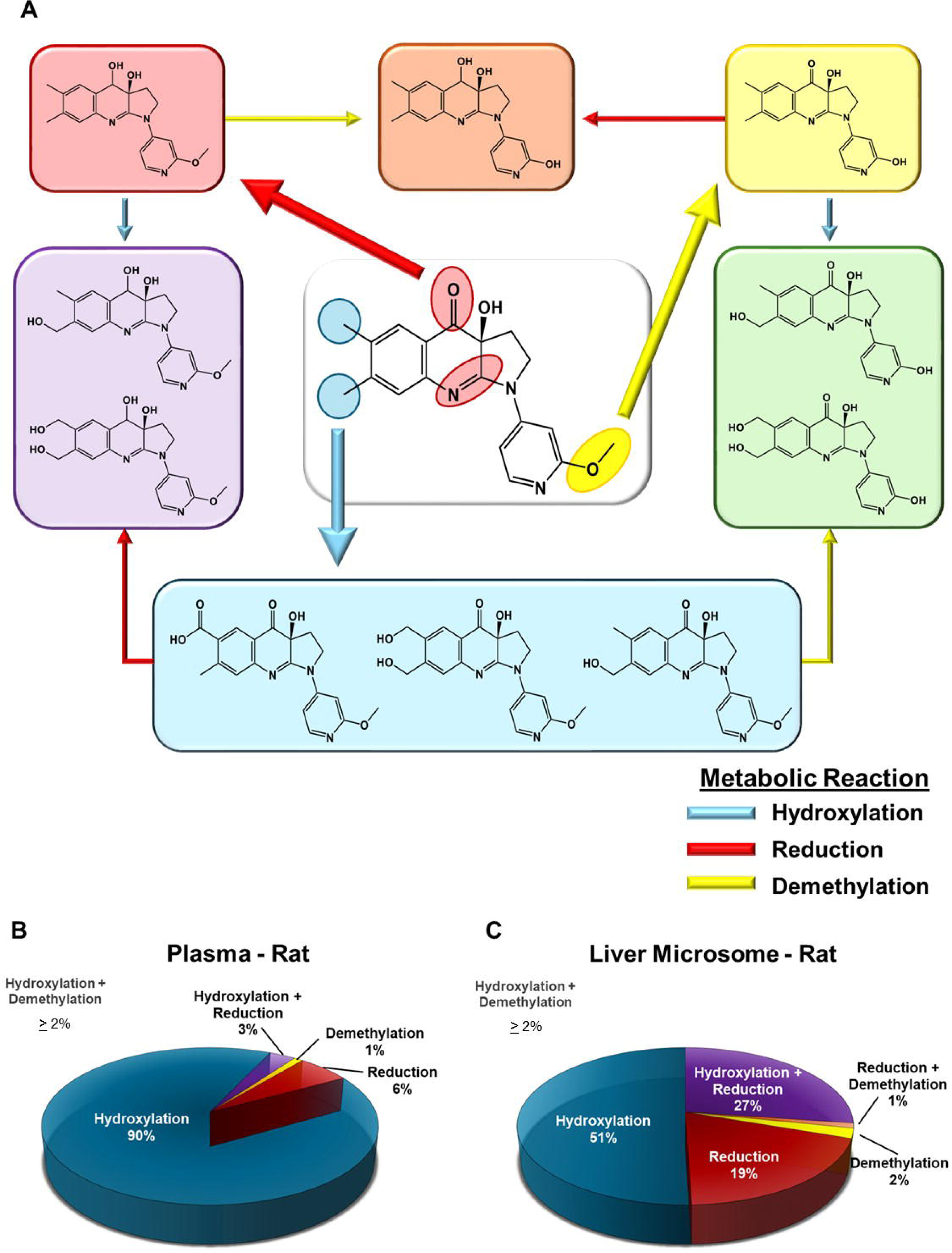
Identification of MT-288 metabolites. A) The sites of metabolism of MT-288 were identified and structures are depicted and grouped using primary colors according to the type of reaction. These correspond to demethylation (yellow), reduction (red), and hydroxylation (blue). Metabolites containing more than one type of modification use the corresponding secondary color. Multiple mono-hydroxylation and two reduction products were identified, and a single representative structure is provided to reduce the complexity of the presented metabolites. The relative content of the individual MT-228 metabolites were determined from rat plasma 30 minutes after a 10 mg/kg IP injection B) and compared to *in vitro* samples generated during a 30-minute incubation in rat liver microsomes. C) Values in the pie charts reflect the percentage of the total peak area the individual metabolites represent. Slices without a listed value contributed less than 2% of the total.

### MT-228 Selectivity Insights Through Crystal Structure

To reveal the molecular basis of **MT-228**’s improved selectivity, we obtained crystals of the human SmMII motor domain (MD) bound to **MT-228** and Mg.ADP-vanadate. **MT-228** is a four-fold more potent inhibitor of SmMII (K_I_ = 4.0 µM) than CMII (K_I_ = 17.0 µM) and the sequence of SmMII MD is 83.6% identical to that of NMIIB, enabling some investigation into potential sources of the compound’s selectivity. The map at 2.4 Å shows clear electron density for **MT-228** in the pocket, previously described as the binding site for Blebb (**Fig. 3A-F**) (*60*). Most of the interactions of **MT-228** with residues found in the binding site are similar to those of Blebb (**Table S4**), with the predominant contacts being of hydrophobic nature. Polar contacts with Gly243, Leu265 and Ala461 are maintained. The methoxy group on the D ring makes close contact with Cys475 (**Fig. 3B-D)**.

**Figure 3.**
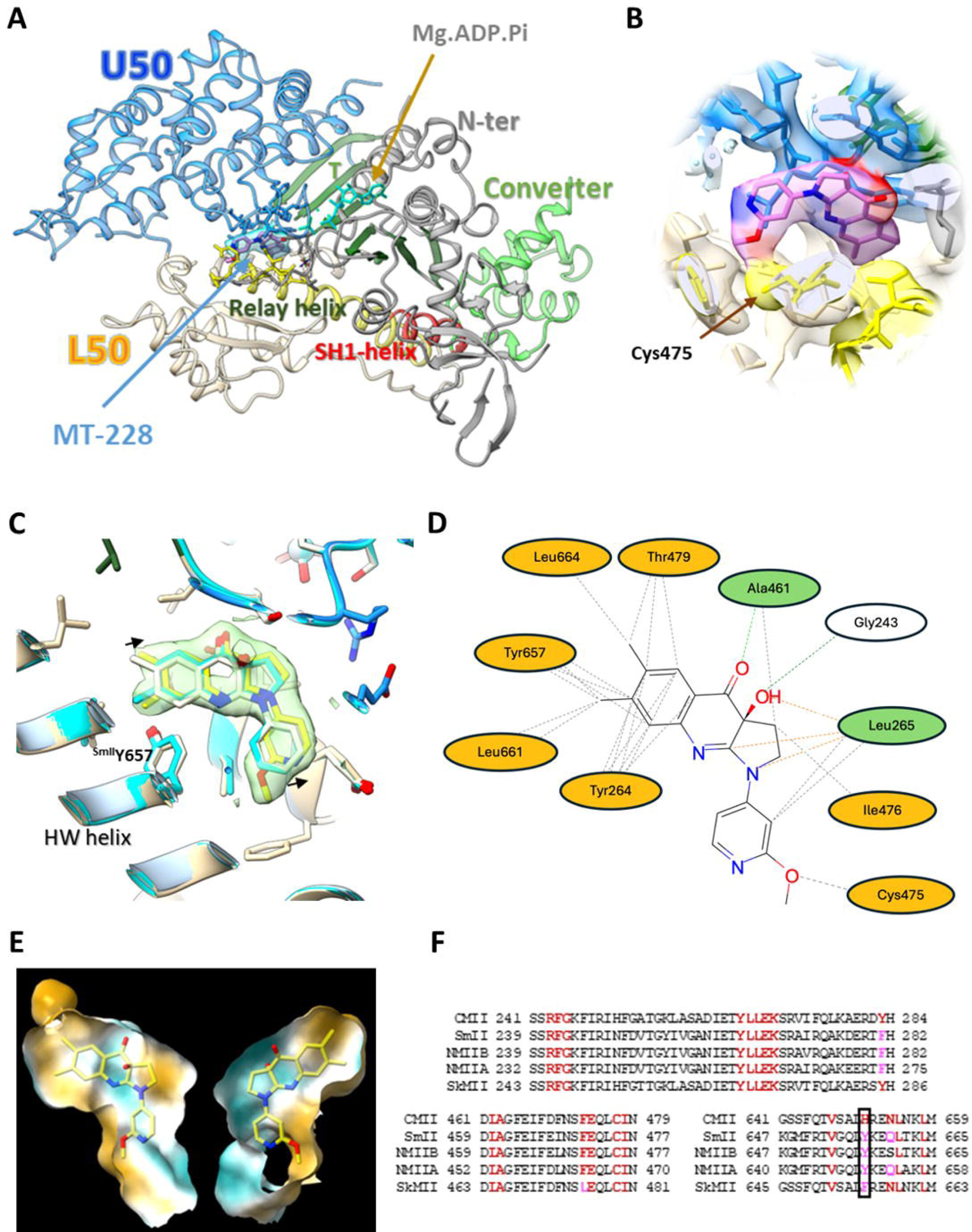
Structural description of the binding site of MT-228 in SmMII. A) Crystallographic structure of the SmMII motor domain (MD) in complex with MT-228. Positioning of the nucleotide and of the compound is indicated with arrows. The beta-sheet defined as the transducer is labeled with a *T*. Electron density corresponding to the compound is shown. B) Close-up of the binding pocket. Electron density corresponding to MT-228 and the surrounding residues is shown at contour level 1.0. C) Superposition of SmMII·MT-228 (yellow and beige carbons) with Dicty·BL6^2^ (blue carbons) and Dicty·Blebb (light gray carbons). Tyr657 is outlined as it plays a major role in defining specificity of MT-228 (see text); black arrows outline the shift in positions of equivalent atoms of MT-228 compared to the other two compounds. D) Residues forming the MT-228 binding site. Orange: hydrophobic contacts; white: polar contacts; green: residues that exhibit both polar and hydrophobic contacts with MT-228. E) Hydrophobicity of the binding site (blue: polar surface; brown: hydrophobic surface). F) Sequence alignment. Tyr657 and the homolog residues are outlined with a rectangle. This figure was prepared with ChimeraX 1.7.1 (A, B, C and E) and Cresset Flare 8.0.0 (D).

Comparison with the Blebb structure (PDB 1YV3) shows that the position of **MT-228** is displaced by up to 1.0 Å, further away from the HW helix (**Fig. 3C** and **Movie 1**), a finding only made possible by the experimental determination of the structure of this complex. Compared to Blebb, the **MT-228** additional methyl group on the A ring and methoxy extension of the D ring are positioned on the same side of the compound (**Fig. 3C**). Both modifications contribute to the observed displacement, since adding a methyl to the A ring of the previously known Blebb analog BL6 (PDB 3BZ8) (*61*) only slightly shifts its position (0.5 Å on the longest distance) when compared to Blebb, while it has no extension on the D ring. The position of the D ring of BL6 shows no shift at all (**Fig. 3C**). This is in agreement with the synergistic selectivity improvements associated with these modifications (see **Table 2**): introduction of the extra methyl group in **MT-142** leads to a 2.6-fold increase in selectivity for NMIIB over CMII relative to Blebb (Supplemental Results), while addition of the methoxy-group further increases selectivity in **MT-151** and **MT-228** (15- and 17-fold improvement relative to Blebb, respectively).

The sequence of SmMII MD (residues: 1-785) is 83.3% identical to that of NMIIA (residues: 1-778), and 83.6% to NMIIB (residues: 1-785), with all significant differences located away from the inhibitor binding site (**Fig. S5**). For comparison, sequence identities of 51.0% and 49.6% are found for CMII (residues: 1-780) and SkMII (residues: 1-784), respectively. Thus, this high sequence similarity allows for reliable homology modelling of the structural basis for **MT-228**’s selectivity for NMIIs (**Fig. S6**). Compared to SmMII, the compound is 4.3 times less active for CMII, 1.7 times more active for NMIIA, and 4 times more active for NMIIB (**Table 2**). Interestingly, almost all residues found near the compound are conserved between these myosin IIs. The main sequence difference in the inhibitor binding site is located on the HW helix (**Figs. 3C**, **3F**, and **S5**). The hydrophobicity and size of a residue in close contact with the apolar A and B rings of **MT-228** differ in these class II myosins. Both SmMII and NMII have a tyrosine at this position (^SmMII^Tyr657 in SmMII and ^NMIIB^Tyr657 in NMIIB, ^NMIIA^Tyr650 in NMIIA), while CMII has a histidine (^CMII^His651), which exhibits a potentially charged and smaller side chain. Interestingly, SkMII has a phenylalanine at this position (Phe655), which also correlates with a good potency of the compound for this myosin (**Figs. 3F, S6**). Thus, it appears that this residue difference on the HW helix together with the displacement of the compound in the pocket relative to the position occupied by Blebb contribute to the selectivity of diverse class II myosins for **MT-228**.

### In vivo Tolerability of MT-228

Given **MT-228**’s improved selectivity profile and cardiac safety, we next assessed the tolerability of the compound *in vivo*. Locomotion, an important measure to assess given **MT-228**’s activity at SkMII, was first determined in the open field. Mice were infused with either vehicle, 5, 7.5 or 10 mg/kg (IV) five minutes before being placed in an open field for 40 minutes. No significant effects on distance travelled over the first 10 min or entire 40 minute test period post-injection were observed (10 min: F_(3, 10)_ = 0.14, *P* > 0.05; 40 min: F_(3, 10)_ = 0.37, *P* > 0.05, **Fig. S9A-B**). Furthermore, there was no effect on velocity (Interaction of Time x Treatment over 10 min time bins: F_(9,30)_ = 0.21, *P* > 0.05; **Fig. S9C**) and no difference between time spent in the center versus the sides of the open field, a measure of anxiety-like behavior (Interaction of Location x Treatment: F_(3, 9)_ = 0.53, *P* > 0.05; **Fig. S9D**). Clinical observations revealed no general health effects at any of the doses tested. Taken together, these data indicate that **MT-228** is well-tolerated up to at least 10 mg/kg (IV).

During early tolerability studies, mice were administered **MT-228** intravenously via the tail vein. While the drug was well-tolerated and had no effect on open field behavior, the infusion process was sufficiently stressful to disrupt the expression of more complex behaviors, such as conditioned place preference (CPP; data not shown). Therefore, we shifted to intraperitoneal (IP) administration, a more commonly used route of drug delivery for rodent behavior studies. To ensure sufficient brain exposure to **MT-228** via IP route of administration, we determined drug concentrations in the brain following delivery of 2 mg/kg (IP) **MT-228**, finding that concentrations were approximately 0.34 μM in the brain, representing a B:P of 2.9 (**Table 1**). All further studies utilized IP route of administration, unless otherwise stated.

### MT-228 Disrupts the Motivation for Methamphetamine

Having demonstrated excellent tolerability of **MT-228** with no observable adverse effects up to 10 mg/kg, *in vivo* behavioral efficacy was next determined. To assess the impacts of **MT-228** on the motivation for METH, mice underwent training for METH-associated CPP (**Fig. 4A)**. Two days after the last training session, mice were tested for their place preference in the absence of METH reinforcement to determine context-associated motivation for the drug. Thirty minutes before testing, mice were given one of four doses of **MT-228** (0.5, 1, 2.5 or 5mg/kg; IP) or vehicle (Veh). As expected, Veh-treated mice displayed a significant preference for the METH-paired chamber (CS+) (Veh: CS+ vs CS-W = 56, *P* < 0.05). This was disrupted by all doses of **MT-228** (0.5 mg/kg: W = 5, *P* > 0.05; 1mg/kg: W = 13, *P* > 0.05; 2.5 mg/kg: W = -31, *P >* .05; 5 mg/kg: W = -4, *P* > 0.05; **Fig. 4A**), establishing that **MT-228** disrupted the motivation for METH at doses as low as 0.5 mg/kg.

**Figure 4.**
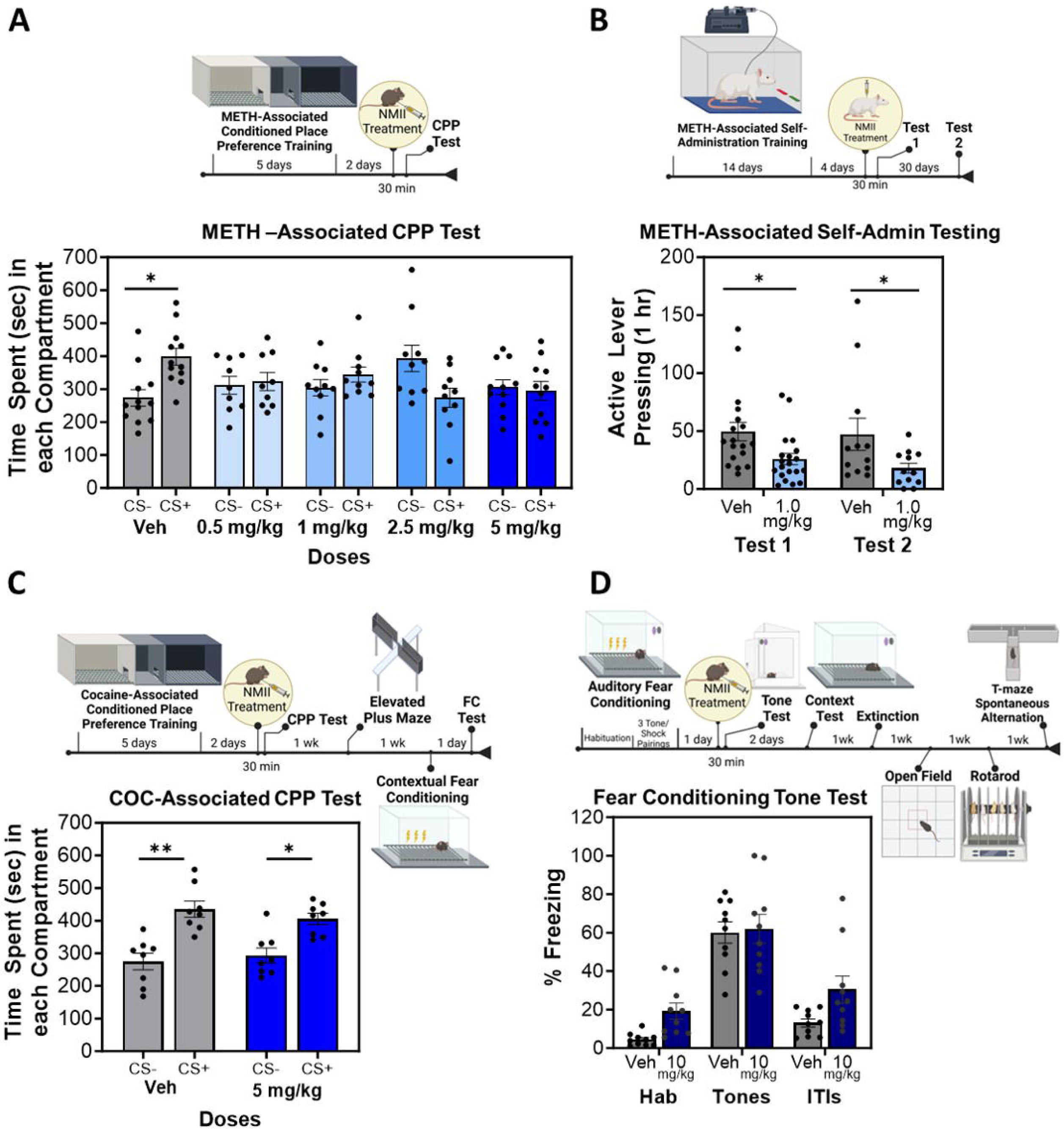
MT-228 produces a selective disruption of the motivation for METH. A) Doses as low as 0.5 mg/kg MT-228 (IP) disrupted METH-associated CPP in mice and B) 1 mg/kg MT-228 (IV) disrupted METH seeking following self-administration in rats. C) 5 mg/kg MT-228 (IP) had no effect on COC-associated CPP and D) 10 mg/kg MT-228 (IP) had no effect on auditory fear conditioning. ITI = Times between tone presentations. Error bars represent SEM, * *P* < 0.05.

The impact of **MT-228** on the motivation for METH was further assessed using IV drug self-administration, which is considered the gold standard for modeling drug-seeking behavior (**Fig. 4B**) (*62*). Rats were given daily access to METH for two weeks via pressing a lever in the self-administration chamber, followed by four days of forced abstinence in the home cage. Rats then received 1 mg/kg **MT-228** or Veh (IV) 30 minutes before testing. Because the animals had indwelling catheters through which they received multiple daily IV infusions of METH during training, the single IV administration of Veh or **MT-228** through the catheters was not a stressor. For testing, rats were returned to the self-administration chamber and lever pressing, which was not reinforced with METH infusion, was assessed as a measure of the motivation to seek METH. The robust lever pressing displayed by rats that received Veh infusions was markedly reduced in animals treated with 1mg/kg **MT-228** (Test 1: U = 87.50, *P* < 0.001; **Fig. 4B**). At the end of the one hour test, rats were returned to their home cages for 30 days. On Day 30, they were given a second METH seeking test, but no additional **MT-228** treatment. The disruption of METH seeking present on Day 1 was still present at the second test, a month later (Test 2: U = 37.50, *P* < 0.05; **Fig. 4B**), establishing that a single low dose infusion of **MT-228** removed the motivation to seek METH for at least one month.

### Selectivity of MT-228 for Methamphetamine Behaviors

Given that systemically administered **MT-228** produces a rapid and lasting effect on METH seeking, a broad battery of behavioral tasks was next used to confirm that the drug does not produce any immediate or lasting generalized effects on neurological processes such as memory, locomotion or anxiety-like behaviors. Tasks that require a wide range of brain subregions were intentionally selected to test for any impacts of NMII inhibition beyond the BLA, through which NMII inhibition disrupts the motivation for METH by reversing structural neuroplasticity (*23, 31, 35*). The first battery of behaviors consisted of cocaine (COC) CPP to determine the impact on motivation for another closely related stimulant, elevated plus maze, a measure of anxiety-like behavior, and contextual fear conditioning to assess hippocampus-dependent long-term associative memory (**Figs. 4C, S10A**). Thirty minutes before testing for a COC-associated place preference, mice received vehicle or 5 mg/kg (IP) **MT-228**, a dose 10-fold higher than the minimally effective dose (MED) in METH CPP (**Fig. 4A**). As expected, **MT-228** had no effect, with both groups expressing a strong preference for the COC-paired chamber (Veh: W = 36, *P* < 0.01; MT-228 W = 34, *P* < 0.05; **Fig. 4C**). One week later, the same mice were tested in elevated plus maze (**Fig. S10B**). Prior exposure of the brain to **MT-228** (administered at the time of COC CPP testing) had no effect on any measure in elevated plus maze, including the total distance travelled in the maze (*P* > 0.05), latency to enter the open (unprotected) arm (*P* > 0.05), or time spent in the open arm (*P* > 0.05; **Fig. S10B**). One week later, the mice underwent contextual fear conditioning and were tested for retrieval of the long-term memory the next day. Again, the prior **MT-228** treatment had no effect, as expected ( T_(14)_ = 1.5, *P* > 0.05; **Fig. S10C**), indicating the ability to learn, consolidate and retrieve a long-term memory with the hippocampus, which relies on actomyosin dynamics to drive the underlying structural neuroplasticity (*30*).

The second behavior battery tested potential treatment effects on the expression of an amygdala-dependent long-term auditory fear memory, extinction of that fear memory requires new learning (*63-65*), open field, rotarod and spontaneous alteration (**Figs. 4D, S11A**). Given the total lack of effect of 5 mg/kg (IP) **MT-228** on the first battery, the dose was increased further to 20-fold the MED in METH CPP (10 mg/kg). Twenty-four hours after training for auditory fear conditioning, mice received vehicle or **MT-228**, followed by a memory retrieval test cued by an auditory tone. Again, **MT-228** had no effect, as expected (Tone Test: F_(1, 18)_ = 3.15, *P* > 0.05; **Fig. 4D**). Similarly there was no effect on a hippocampus-dependent context retrieval test (F_(1, 18)_ = 0.025, *P* > 0.05; **Fig. S11B**) or extinction learning (F_(1, 18)_ = 1.16x10^-4^, *P* > 0.05; **Fig. S11B**). One week later, these same mice were run through two assays to assess different aspects of mobility and coordination, open field and rotarod (**Fig. S11A**). Prior **MT-228** treatment had no effect in open field on distance travelled (*P* > 0.05), average velocity (*P* > 0.05) or time spent in the center and sides/corners (Group x location: F_(1, 18)_ = 1.316, *P* > 0.05; **Fig. S11C**), and no effect on coordinated movement in rotarod (*P* > 0.05; **Fig. S11D**). Short-term spatial memory was also assessed in these mice using spontaneous alternation with both no delay and a 1-minute delay. Again, there was no effect of **MT-228** (Delay x Group: F_(1, 18)_ = 0.1304, *P* > 0.05; Fig. S11E).

Associations for both METH and fear rely on the BLA and we have previously shown that the disruption of a consolidated METH association is achieved by specific inhibition of NMII in the BLA (*23*). Therefore, as a final test of the selectivity for METH via NMIIB targeting, a combined genetic and pharmacologic approach was taken to completely suppress NMIIB function in the BLA during the expression of an amygdala-dependent auditory fear memory to ensure that, unlike METH, it remains intact (**Fig. S11F**). One day after auditory fear conditioning, an siRNA targeting NMIIB’s heavy chain, *Myh10*, or a control siRNA was injected directly into the BLA via indwelling cannulae. We have previously reported that this procedure and timeline selectively reduces *Myh10* mRNA levels by close to 60% in the BLA (*23*). The following day, *Myh10* siRNA mice also received an injection of **MT-228** (10 mg/kg, IP) to further inhibit NMIIB function, while control siRNA mice received a Veh injection, followed by an auditory tone test. Even this combined genetic and pharmacologic targeting of NMII had no effect on the expression of a consolidated fear memory (F_(2, 28)_ = 0.26, *P* > 0.05; **Fig. S11F**). Taken together, these data provide strong support for **MT-228**’s ability to selectively disrupt the motivation for METH without impacting other brain processes and highlights its potential for development as a medication for the treatment of MUD.

## DISCUSSION

There are a wide range of potential disease applications for a safe and selective small molecule inhibitor of NMII. Examples include but not limited to glaucoma (64), erectile dysfunction (65), liver fibrosis and portal hypertension (66), overactive bladder syndrome and the associated depression (67, 68), eryptosis (69), neointimal hyperplasia after vascular injury (70), airway hyper-responsiveness (71), arterial thrombosis (75), and ototoxic drug-induced hair cell loss (76). To begin to address these needs, here we report the synthesis and physicochemical, biochemical and *in vivo* characterization of novel derivatives of the pan-myosin II small molecule inhibitor, Blebb. With the goal of developing NMII inhibitors that are safe for *in vivo* administration, we reasoned that Blebb’s potent inhibition of CMII (*38, 54*) was likely the dose limiting liability. By examining structure-activity relationships (SAR) with a focus on reducing CMII activity, we showed that combined modifications to the A and D ring of Blebb improved selectivity for NMII and reduced potency for CMII, resulting in a strikingly improved *in vivo* safety profile. Indeed, these modifications to Blebb resulted in marked decreases in CMII activity. For example, **MT-106** was greater than 50-fold less potent at CMII than Blebb. Importantly, decreased CMII potency corresponded to profound improvements of derivatives over Blebb on cardiac function, as measured by contractility of human iPSC cardiomyocytes and *in vivo* cardiac output in an echocardiogram. Our results indicate that a CMII K_I_ of 10 μM or greater supports well-tolerated *in vivo* dosing of the NMII-targeting derivatives (dose determined by NMII potency (**Fig. 1**)). A number of derivatives showed improved potency at the primary target, NMIIB, to submicromolar levels (e.g. **MT-152**, NMIIB K_I_ = 410 nM). However, the hydrophobic nature of the binding pocket (**Fig. 2E**) imposes an inverse correlation between NMII potency and solubility, precluding the usefulness of those derivatives as drug candidates. Therefore, we selected **MT-228** for additional studies, as it presented a balanced profile of desirable qualities, including excellent NMIIB potency, photostability, solubility and brain penetrance, as well as weak CMII inhibition and *in vivo* cardiac tolerability. **MT-228** was well-tolerated and behavioral efficacy studies revealed long-lasting disruption to METH seeking with a single administration at doses as low as 0.5 mg/kg. The highest dose tested for tolerability and behavioral selectivity was 10 mg/kg and that showed no observable adverse effects on tolerability or in assays that require intact functioning of a wide array of brain regions, including the hippocampus, cortex, striatum and cerebellum. Taken together, this translates into a therapeutic index greater than 20-fold. ADMET profiling of **MT-228** revealed it has a number of additional properties that make it an excellent preclinical candidate for the treatment of METH use disorder. It is not cytotoxic, genotoxic or cardiotoxic. **MT-228** demonstrated minimal CYP inhibition (*66, 67*) and did not induce PXR activation (*68*), indicating that it is unlikely to interfere with the metabolism of other drugs. In further support of its anticipated tolerability, *in vitro* safety pharmacology profiling demonstrated that **MT-228** is highly selective. Finally, the long-lasting efficacy seen with a single administration of **MT-228** is considered a particular strength from the perspective of tolerability, but also for the treatment of METH use disorder specifically, as patient compliance can make daily, chronic treatment options a challenge for this clinical population.

Of the four myosin II’s, Blebb is most potent against SkMII (SkMII K_I_ = 280 nM). And, while we successfully reduced CMII potency, SAR demonstrated that SkMII is the most tolerant of modifications to Blebb, making it the simplest target to maintain. Indeed, of the 471 derivatives made in this medicinal chemistry campaign, only 102 showed a marked (>100-fold) loss of SkMII potency and in most cases, that was accompanied by a complete inactivation of the compound across all myosin II’s assayed (58.8% of total). Further, we and others have previously published SkMII analogs of Blebb, such as MT-134, which has a 173-fold preference for SkMII over CMII, was well-tolerated *in vivo* with systemic administration, achieved high levels in skeletal muscle, and produced the expected *in* vivo effect of interfering with coordinated motor function (*37, 38, 69-71*). Fortunately, at the doses of **MT-228** needed to inhibit the NMIIB target, skeletal muscle effects are not apparent. This is likely due to the high levels of SkMII in the body and degree of saturation that must be achieved to interfere with skeletal muscle function. Approximately 40 % of the total body weight is skeletal muscle tissue (*72*) and the protein content of skeletal muscle is ∼177 µg protein/mg tissue (*73*), from which ∼25% is SkMII (*74*). With a molecular weight of ∼220 kDa, the estimated SkMII concentration in skeletal muscle tissue must be ∼200 µM (∼44 mg/ml). The relatively large doses (>30 mg/kg) of a Blebb analogue specific for SkMII, and the resulting tissue concentrations (> 30 µM in skeletal muscles) necessary to interfere with skeletal muscle function *in vivo* in rats (*37*) is consistent with the above estimates.

It is very difficult to speculate how the interaction between these inhibitors and the amino acid residues of the binding pocket are altered in each myosin isoform based on sequence comparisons and structural models predicted by the current algorithms. A 2.4A crystal structure of the enzyme-inhibitor complex demonstrated that **MT-228** achieves selectivity for NMII over CMII through a mechanism that depends on a shift in the position of the molecule within the binding pocket relative to that of Blebb. The atomic resolution structure reveals that His651 in CMII is a difference in close proximity of the bound inhibitor that could explain, in large part, this selectivity. Instead of a histidine at residue 651, a phenylalanine is found in SkMII, whereas tyrosine is at this location in NMIIB, NMIIA, and SmMII. Most of the other residues in the binding pockets are identical.

Interestingly, the compound **MT-140** is selective for CMII over NMII (K_I,CMII_ = 2.6 µM, EC_50,NMIIB_ = 24 µM, see **Table 2**). The selectivity of **MT-140** and **MT-228** are shifted in opposite directions for two reasons: **MT-140** has reduced NMIIB potency relative to the non-selective Blebb (K_I,CMII_ = 1.9 µM, EC_50,NMIIB_ = 1.9 µM), while **MT-228** is less potent on CMII (EC_50,NMIIB_ = 1.0 µM, K_I,CMII_ = 17 µM). The crystal structure of **MT-140** in complex with *Dictyostelium* myosin (PDB 3BZ9) reveals that the position of the compound is also shifted relative to the position occupied by Blebb, due to the lack of the methyl group attached to ring “A” in Blebb (*61*). However, this shift occurs in the opposite direction compared to **MT-228**, apparently leading to energetically unfavorable interactions within the inhibitor binding pocket of NMIIB (**Fig. S7**).

**MT-228** is well-suited for the treatment of METH use disorder. However, there is room to further improve the compound for other uses, particularly those in which prolonged inhibition may better impact disease progression, such as axonal regeneration after neural injury (*75*). The metabolism pathway identification suggests that further optimization of **MT-228** analogs would benefit from a focus on minimizing oxidation of the A Ring-methyl substituents on the quinoline and the reduction of the ketone and/or amidine. In terms of selectivity, further decreases in SkMII inhibition would be desirable, even if skeletal muscle effects are not apparent with **MT-228**. In summary, the results of the medicinal chemistry campaign presented here provide new tool compounds for the scientific community and most importantly, **MT-228** serves as a promising lead molecule for future medication development.

## ACKNOWLEDGEMENTS

We thank the NINDS Blueprint Neurotherapeutics program for their support, particularly Dr. Mohamed Hachicha. We also thank Susan Khan (UF Scripps) for technical assistance. In addition, we thank the beamline scientists of Proxima 2A from the Soleil synchrotron for their excellent support during data collection. We also thank Marie Juillé for help in the analysis of the binding site. E.J.Y. was supported by grant R44DA055420. A.H. was supported by grants from CNRS and ANR-21-CE11-0022-01. C.A.M., P.R.G. and T.M.K. were supported by grant UH3NS096833. C.A.M. was also supported by grant R01DA049544.

## AUTHOR CONTRIBUTIONS

Conception and design: LR, EJY, CK, KT, TKB, GR, MDC, MS, AHJ, PRG, TMK, CAM

Development of methodology: LR, EJY, CK, KT TKB, MDC, MS, AHJ, PRG, TMK, CAM

Reagent supply: HLS, JS

Acquisition of data: LR, EJY, CK, MH, RFS, LL, KT, PP, XJ, MG, LH, JRP, AP, MC, SB, TKB

Analysis and interpretation of data: LR, EJY, CK, KT, TKB, GR, MDC, MS, AH, PRG, TMK, CAM

## AUTHOR DECLARATIONS

Authors C.A.M., P.R.G., T.M.K. and E.J.Y. hold equity positions in Myosin Therapeutics, which has licensed these NMII inhibitors from Scripps Research and the University of Florida. Authors C.A.M., P.R.G., and T.M.K are members of the Myosin Therapeutics Board of Directors. The following patents and patent applications are related to this work: WO2019241469A1 and US63/431,234.

## MATERIALS AND METHODS

For complete methods, please see Supplemental Information.

### Data analysis

All behavioral experiments were analyzed using a one-way or two-way repeated measures Analysis of Variance (ANOVA) to examine differences between treatments or treatment by day interactions, respectively, where applicable. Some data was analyzed by paired or independent *t*-test or Mann Whitney test, where applicable. All *post hoc* tests were conducted, when appropriate, using Tukey’s Honest Significance Difference (HSD) test.

### Materials Availability

Small molecule inhibitors generated in this study will be made available on request with a completed materials transfer agreement.

## SUPPLEMENTAL INFORMATION

- Document 1:

○ Supplemental Results
○ Supplemental Methods
○ Table S1-S5 and Figures S1–S11

