## Supplemental Information for "Development of Clinically Viable Non-Muscle Myosin II Small Molecule Inhibitors with Broad Therapeutic Potential"

Supplemental Results

**Synthesis, Purification and Characterization of Blebbistatin Derivatives**

*Modifications to the D Ring.* First, a series of ring “D” modified Blebb derivatives was synthesized (**Table 2**). Because we have previously determined that NMIIB inhibition is sufficient to disrupt METH seeking, NMIIB inhibition, cell membrane permeability, and potential cytotoxicity of these compounds were first evaluated in our *in vitro* COS7 cell-based cytokinesis assay, which includes a measure of cytotoxicity (*1*). Using the steady state ATPase assay (*2*), we also determined inhibitory constants (K_I_) for CMII and SkMII. These data were then used to determine selectivity of the compounds for NMIIB relative to CMII, with a goal of at least 10-fold selectivity for NMII over CMII. Because of the importance to the SAR strategy, *in vivo* plasma exposure and brain penetrance were also routinely determined (*3*).

We have previously reported that, compared to Blebb, the ortho-methoxy- substituent in **MT-102** improved solubility and abolished potency on SkMII (*3*). We found a similarly drastic decrease in CMII and, unfortunately, NMIIB potency with this compound (**Table 2, Figs. S1A, S2A**). In **MT-103**, the methyl- substituent in the meta-position reduced potency on CMII and SkMII (*3*). Although this modification produced a ~4-fold decrease in NMIIB potency, the selectivity over CMII and SkMII was significantly improved (e.g. 6.8-fold increase in the selectivity of **MT-103** for NMII over CMII compared to Blebb). No improvements in solubility or photostability were observed (**Table 2, Figs. S1B, S2B**).

Another meta-substituted analog, **MT-106** showed a similar pattern of effects, with slightly greater potency on NMIIB (EC_50_ = 5.1 µM), resulting in a better overall selectivity profile for NMIIB over CMII (>22-fold) and SkMII (4.1-fold) (**Table 2, Figs. S1C, S2C**). Moreover, the meta-chloro substitution also led to significantly improved photostability (**Table 2**). Methyl- and cyano- substituents in the para- position in **MT-116** and **MT-117**, respectively, led to a ~2-fold increase in NMIIB potency, with no significant change in CMII inhibition compared to Blebb (**Table 2, Figs. S1D-E, S2D-E**). This resulted in slightly improved selectivity for NMIIB over CMII in both cases. Interestingly, while the methyl- substituent had minimal impact on SkMII potency, the cyano-derivative proved to be a strong SkMII inhibitor with a K_I_ of only 66 nM (**Fig. S2E**). The same para-cyano-substituent led to a substantial improvement in photostability compared to Blebb, while maintaining a similar solubility. The para-methyl substituent in **MT-116** did not improve photostability, but drastically reduced solubility (**Table 2**).

To determine the extent to which the CMII K_I_ in the ATPase assay predicted impacts on cardiomyocyte contractility, we tested several of these derivatives in the hIPS cardiomyocyte assay used to test Blebb (**Fig. S3A**). Consistent with their respective CMII K_I_’s, **MT-102** (CMII K_I_ = >450 µM) had no measurable impact on cardiomyocyte contractility (Cell Index Changes from DMSO control: *P* > 0.05 at all doses; **Figs. 1B, S3B-C**). **MT-103** (CMII K_I_ = 50 µM) and **MT-106** (CMII K_I_ = 110 µM) had minimal effects at 0.1-1 µM (Cell Index Changes from DMSO control: *P* > 0.05 at 0.1-1 µM). However, both compounds produced a reduction in contractility of >25% at the highest concentration (3 µM; *P* < 0.05) (**Figs. 1B, Fig. S3B-C**). This was unexpected because of their very low potency at CMII in our *in vitro* ATPase assay. Moreover, **MT-116** (CMII K_I_ = 1.7 µM) had strong effects, even at submicromolar concentrations (Cell Index Changes from DMSO control: *P* < 0.05; **Figs. 1B, Fig. S3B-C**). The highly ordered actomyosin system of sarcomeres in living cardiomyocytes and cardiac muscle (*4*) obviously differs from the randomly interacting actin and myosin filaments in a biochemical assay. Also, while the readout in the biochemical assay depends on a reduction in ATPase activity, the cardiomyocyte impedance assay depends on a reduction in force generation. These differences may explain the increased potency of the compounds in the hIPS cardiomyocyte assay, highlighting the value of definitively determining cardiac effects of promising derivatives *in vivo* with echo studies of the adult heart.

For the D ring group of modified derivatives, **MT-106** was selected for additional characterization to support further medicinal chemistry optimization efforts, as it had the widest CMII to NMII selectivity ratio (>22-fold). Several ADMET properties were next assessed, using Blebb as a benchmark (**Table S2-3**). All assays utilized positive controls. The cytochrome P450 (CYP450) inhibition profile was first assessed for analogs in order to predict the potential of compounds to inhibit these enzymes, as this could alter the actions of concomitant medications (*5, 6*). Next, hERG (human ether-a-go-go related gene) inhibition activity (*7*) and hERG-CHO patch clamp functional activity were determined (*8*). The hERG channel regulates cardiac repolarization and drug-induced hERG dysfunction can cause long QT syndrome and sudden death (*9*). Metabolic stability was determined in human, rat, mouse, and dog liver microsomes, as well as P-glycoprotein (P-gp) substrate analysis in MDR1-MDCK cell monolayers to determine the likelihood that a compound will be effluxed from the brain by the P-gp transporter, located within the vessel walls of the brain’s capillaries (*10-13*). P-gp substrates do not make for effective CNS therapeutics. Finally, plasma protein binding (PPB) was also assessed, as the free fraction of a compound can impact its pharmacokinetics and *in vivo* activity (*14*). CYP inhibition by Blebb was greater than 50% for CYPs 2B6, 2C9, 2D6, 3A4 and 2C8 (**Table S2**). The profile was overlapping, but distinct for **MT-106**, showing greater than 50% inhibition on CYPs 1A2, 2B6, 2C19 and 3A4 (**Table S2**). Blebb and **MT-106**, as well as all other analogs tested in this study lacked potent inhibition of hERG, such that IC_50_’s either could not be generated or were greater than 10mM (**Table S2**). Similarly, IC_50_’s could not be determined or were greater than 10mM in the hERG-CHO patch clamp assays and potency for all analogs, including **MT-106**, ranked as low. Blebb and **MT-106** were rapidly metabolized in the four species of liver microsomes tested (human, rat, mouse and dog; **Table S3**). Further, Blebb and all analogs tested, including **MT-106**, were determined to not be substrates for P-gp and had high PPB (> 90%; **Table S2**).

*In vivo* pharmacokinetic parameters were also determined for **MT-106** following a single infusion of 0.5 mg/kg (IV). Compared to Blebb, **MT-106** had a shorter half-life (1.5 h vs 3.3 h), but approximately 2-fold higher C_max_ and AUC at 136.5 ± 21.5 ng/mL and 94.7 ± 11.3 ng/mL, respectively (**Table 1**). Consistent with **MT-106** not being a P-gp substrate, it had high brain penetrance, with a 3.5 brain to plasma ratio (B:P) 30 minutes after IV infusion (**Table 1**).

In summary, ortho-substituted D-ring analogues are most likely suboptimal myosin II inhibitors in general. The introduction of para-substituents slightly increased potency and may result in other favorable properties (e.g., improved photostability). In general, meta-substituents significantly increased selectivity over CMII, but decreased NMIIB potency.

*Modifications to the A Ring.* To further explore ways of enhancing selectivity, while increasing NMIIB potency, Blebb derivatives with ring “A” modifications were synthesized, as shown in **Table 2**. Replacement of the “-R_1_” methyl-group of Blebb by “-H” in **MT-140** resulted in a greater than 10-fold reduction in NMIIB potency (K_I_ = 24 µM), with only a slight decrease in CMII potency (K_I_ = 2.6 µM; **Figs. S1F, S2F**) and a corresponding suppression of hIPS cardiomyocyte contractility at 0.3 μM and above (Cell Index Changes from DMSO control: *P* < 0.05 at 0.3-3 μM; **Figs. 1B, S3B-C**). To further validate the hIPS-CM results and to confirm the poor tolerability of Blebb in the echo assay was due to CMII effects, **MT-140** was tested by *in vivo* echo at the same doses as Blebb and compared alongside Blebb (**Fig. S4B**). Replicating the results in **Fig. 1D**, Blebb decreased cardiac output at the first and second 0.5 mg/kg infusion. The first 0.5 mg/kg infusion of **MT-140** produced a similar decrease in cardiac output as Blebb (Blebb: F_(2,9)_ = 46.5, *P* < 0.0001; MT-140: F_(1,9)_ = 52.9, *P* < 0.0001; Baseline vs D1+5min *P* < 0.0001, Baseline vs D1+10min *P* < 0.0001, Baseline vs D2 start *P* < 0.0001). But, despite a similar CMII K_I_ to Blebb (2.6 vs 1.9 µM), a second infusion of **MT-140** was not at all tolerated and was stopped (**Fig. S4B**). The difference in tolerability may be related to **MT-140**’s ~5-fold reduction in SkMII potency, and possibly the ~10-fold reduction in NMII potency, relative to Blebb. **MT-140**’s reduced affinity for these myosins may reduce the compound sequestration in skeletal muscles and other tissues, leaving more compound bioavailable for CMII inhibition in the heart (*15*). Reducing the dose by 50% to 0.25 mg/kg readily overcame this dramatic effect, resulting in less than 10% reduction in cardiac output (F_(2,9)_ = 1.90, *P* > 0.05; **Fig. S4B**). Unfortunately, **MT-140** was not only selective for CMII over NMIIB, but also had reduced photostability. The synthesis and purification of Blebb, **MT-102**, **MT-103,** and **MT-140** have been reported previously (*3, 16*).

**MT-141**, an “-R_1_” chloro-substituted analog of Blebb, showed a 2-fold increase in selectivity for NMIIB over CMII. However, overall potency was slightly decreased with no change in photostability or solubility (**Table 2, Figs. S1G, S2G**). Further, **MT-141’s** CMII K_I_ (5.9 µM) remained sufficiently potent to interfere with hIPS-CM contractility (Cell Index Changes from DMSO control: *P* < 0.05 at all doses; **Figs. 1B, S3B-C**). Again, this effect (e.g. 50% beat amplitude decrease at 1 µM **MT-141**) appeared to be somewhat stronger than would be predicted from the compound`s performance in the ATPase assay. Therefore, we have analyzed all cardiomyocyte data further by extracting EC_50_ values (**Fig. S3C**). Although the dose-response data were somewhat limited for this analysis, very reasonable fits to the Hill-equation were obtained by assuming that the maximal decrease in the signal is always -100%. No compounds showed any significant deviation from this simple model, indicating that the contractions of cardiomyocytes depend on a single “functional unit” (sarcomeres) inhibited by our compounds. Comparison of K_I,CMII_ and EC_50,Cardiomyocyte_ values (**Fig. 1B, S3D**) revealed that the *in vitro* ATPase data do, indeed, predict compound performance in the cardiomyocyte impedance assay. The compounds are simply more potent in cells (~6-fold difference).

Finally, introduction of a second methyl-group into the “-R_2_” position in **MT-142** produced a ~2-fold increase in NMIIB potency, with a 2.6-fold improvement in selectivity over CMII and a 5-fold improvement in selectivity over SkMII (**Table 2, Figs. S1H, S2H**). This is quite a significant finding as it is one of the few modifications to the Blebb scaffold that led to an improvement in potency. Bis-methyl substitution of the A-ring also improved selectivity for NMII vs CMM which was amplified with additional substitution to the D-ring (**Table 2**). The photostability did not change compared to Blebb (**Table 2**).

In addition to not being a P-gp substrate, having no hERG effects, and high PPB as noted above, ADMET profiling of **MT-140** and **MT-142** revealed greater than 50% inhibition by both compounds on CYPs 2B6 and 2C19 (**Table S2**). **MT-140** also inhibited CYPs 1A2 and 2D6, and was stable in dog liver microsomes (**Table S3**).

*Combined Modification of the A and D Rings.* Although solubility was reduced, the dimethyl- “A” ring clearly appeared the best option to simultaneously improve potency and selectivity. To test this hypothesis, several ring “D” modified derivatives of **MT-142** were synthesized in the next step (summarized in **Table 2**). First, two analogs with methyl- and chloro- substituents in the meta position of ring “D” were synthesized (**MT-150** and **MT-154**, respectively). Combination of these modifications with the dimethyl “A” ring resulted in improved NMIIB potency in both cases, compared to the mono-methyl derivatives **MT-103** and **MT-106**, respectively (**Fig. S1I, L**). **MT-150** also showed ~5-fold improvement in selectivity over CMII. In the case of **MT-154**, this could not be determined. Only a lower limit of K_I,CMII_ could be established in the ATPase assay due to the limiting solubility of the compound (5.8 µM). Similarly, both **MT-150** and **MT-154** showed improved selectivity over SkMII, and decreased potency at SmMII (**Table S1, Fig. S2I, L**). Unfortunately, both the photostability and solubility were negatively affected by the introduction of the second methyl group. A third derivative with a methoxy-group in the meta-position of ring “D” was also synthesized (**MT-151**). Although this compound showed good solubility and selectivity over CMII, SkMII, and SmMII, the photostability was poor (**Tables 2** and **S1, Figs. S1J** and **S2J**).

Next, two analogs with cyano- and methyl- substituents in the para position of ring “D” were synthesized (**MT-152** and **MT-155**, respectively; **Tables 2 and S1, Figs. S1K, M** and **S2K, M**). Introduction of these substituents in combination with the bis-methyl “A” ring resulted in a slightly increased NMIIB potency and corresponding improvement in selectivity over CMII vs **MT-11**7 and **MT-11**6 respectively. Photostability was also greatly improved in the cyano- derivative (**MT-152**). **MT-152** also showed excellent selectivity over CMII (29x). In the case of **MT-155**, poor solubility and photostability were observed. In another compound, **MT-156**, a methoxy-group in the para position of the “D” ring was combined with the dimethyl “A” ring. This compound was potent, and somewhat selective for NMIIB over CMII. However, not only the solubility and photostability, but also the selectivity over SkMII were poor (**Table 2, Fig. S1N, S2N**).

**Tables S2-3** summarize the ADMET profiling for these combined A and D ring derivatives, which were very similar to the A and D ring alone derivatives. Interestingly, **MT-154** displayed greater than 50% inhibition of all CYP450s tested. In terms of microsomal stability, **MT-152** was stable in all species.

*In vivo* pharmacokinetic profiling of **MT-150**, **MT-151**, **MT-152** and **MT-154** was next performed. Like **MT-106**, the half-life of **MT-150** and **MT-151** was reduced by approximately 2-fold relative to Blebb, but the C_max_ and AUC were markedly increased (**Table 1**). **MT-152**, however, was rapidly cleared, with a half-life of only 0.25 hours. This was accompanied by a very low C_max_ and AUC. Interestingly, **MT-152’s** B:P was high at 30 minutes, at 8.2, reflecting a more rapid removal from plasma than brain. **MT-154’s** pharmacokinetic profile was nearly identical to **MT-106**, with the exception of brain penetrance, which was double that of **MT-106** at B:P = 7.1 (**Table 1**).

The pharmacokinetic data were used to support selection of **MT-150** and **MT-152** doses for the *in vivo* echo assay that matched plasma exposure levels of Blebb at 0.5 mg/kg (IV). Consistent with their CMII K_I_’s, neither **MT-150** (K_I_ = 100 µM) nor **MT-152** (K_I_ = 12 µM) impacted cardiac output (MT-150: F_(4,9)_ = 1.2, *P* > 0.05; MT-152: F_(4,9)_ = 0.62, *P* > 0.05; **Fig. S4C**). Given the NMII potency of **MT-152**, we tested it in echo at higher doses. The impact of moving from 0.85 mg/kg to 1.5 mg/kg caused a small, but significant decrease in cardiac output (MT-152 Overall: F_(4,9)_ = 36.0, *P* < 0.0001, Baseline vs D1+5min *P* < 0.0001, Baseline vs D1+10min *P* < 0.0001; **Fig. S4C**). Consistent with the CMII K_I_ (12 µM), further raising the dose to 3.0 mg/kg resulted in a 40% reduction in cardiac output present at 5 minutes post-infusion, that began to recover by 10 minutes (D2 start vs D2+5min *P* < 0.0001, D2 start vs D2+10min *P* < 0.01, D1+10min vs D2+10min *P* < 0.01, **Fig. S4C**). The rapid recovery likely reflects the particularly short half-life of **MT-152**. Although the general selectivity profile of **MT-152** was somewhat promising, the cardiac output effects and poor solubility (< 1.3 µM) limited its usefulness for subsequent studies.

Several of the above mentioned meta- and para- substituents in combination with the dimethyl “A” ring resulted in potent and selective compounds. However, either the photostability or the solubility of these derivatives was considered limiting. Therefore, we next hypothesized that replacement of ring “A” with various nitrogen-containing heterocycles may have a beneficial effect. An N-methyl-pyrazole ring in **MT-157** indeed resulted in enhanced solubility. However, it was accompanied by poor selectivity and photostability (**Table 2, Figs. S1O, S2O**). In **MT-160**, the phenyl “D” ring was replaced by a benzothiazole. This compound showed excellent selectivity for NMIIB over CMII, in combination with acceptable solubility, but poor photostability (**Table 2, Figs. S1Q, S2Q**). Introduction of a similar double-ring (3,4-dihydro-2H-pyrido[3,2-b][1,4]oxazine) in **MT-161** resulted in acceptable selectivity over CMII with excellent solubility. However, photostability was only slightly improved (**Table 2, Figs. S1R, S2R**).

The ADMET properties of **MT-160** are detailed in **Tables S2-3**, with the only notable difference being reasonable stability in all liver microsomes. Pharmacokinetic testing of **MT-160** revealed a nearly identical profile to Blebb, with the exception of higher brain penetrance (B:P = 5.1 vs 2.2 30 minutes post-infusion; **Table 1**).

Supplemental Methods

Proof of purity

1. Purity and Chiral Purity

| **Compound Name** | **Purity** | **Chiral Purity** |
| --- | --- | --- |
| MT-106 | >99% | >99% |
| MT-116 | >99% | >99% |
| MT-117 | >99% | Mixture of Enantiomers |
| MT-140 | >99% | >99% |
| MT-141 | >99% | >99% |
| MT-142 | >99% | >99% |
| MT-150 | >99% | >99% |
| MT-151 | >99% | >99% |
| MT-152 | >99% | 97.2% |
| MT-154 | >99% | >99% |
| MT-155 | >99% | >99% |
| MT-156 | >99% | >99% |
| MT-157 | >99% | 92.3% |
| MT-158 | >99% | 82.7% |
| MT-160 | 97.5% | 82.2% |
| MT-161 | 99.0% | 84.2% |
| MT-228 | 99.0% | 98.6% |

Proof of purity for Blebbistatin, MT-102, and MT-103 has been published elsewhere(*3*).

2. HPLC Conditions

**Method A**

Column: Waters Symmetry 5 µm C18 (250 x 4.6 mm)

Mobile Phase A: Water containing 0.1 % v/v Trifluoroacetic Acid

Mobile Phase B: Acetonitrile containing 0.1% v/v Trifluoroacetic Acid

Detection: 254 nm

Method A Gradient:

| Time (min) | Flow (mL/min) | %A | %B |
| --- | --- | --- | --- |
| 0.0 | 1.0 | 98.0 | 2.0 |
| 20.0 | 1.0 | 0.0 | 100.0 |
| 25.0 | 1.0 | 0.0 | 100.0 |

**Method B**

Column: YMC ODS-AQ C18 120 Å (150 x 4.6 mm)

Mobile Phase A: Water containing 0.1 % v/v Trifluoroacetic Acid

Mobile Phase B: Acetonitrile containing 0.1 % v/v Trifluoroacetic Acid

Detection: 254 nm

Method B Gradient:

| Time (min) | Flow (mL/min) | %A | %B |
| --- | --- | --- | --- |
| 0.0 | 1.0 | 98.0 | 2.0 |
| 15.0 | 1.0 | 0.0 | 100.0 |
| 19.0 | 1.0 | 0.0 | 100.0 |

**Method C**

Column: xBridge 3.5 µm C18 (150 x 4.6 mm)

Mobile Phase A: Water containing 0.1 % v/v Trifluoroacetic Acid

Mobile Phase B: Acetonitrile containing 0.1 % v/v Trifluoroacetic Acid

Detection: 254 nm

Method C Gradient:

| Time (min) | Flow (mL/min) | %A | %B |
| --- | --- | --- | --- |
| 0.0 | 1.0 | 95.0 | 5.0 |
| 20.0 | 1.0 | 0.0 | 100.0 |
| 25.0 | 1.0 | 0.0 | 100.0 |

3. UPLC Conditions

**Method A**

Column: Acquity UPLC BEH 1.7 µm C18 (75 x 2.1 mm)

Mobile Phase A: Water containing 0.1 % v/v Trifluoroacetic Acid

Mobile Phase B: Acetonitrile containing 0.1 % v/v Trifluoroacetic Acid

Detection: 254 nm

Method A Gradient:

| Time (min) | Flow (mL/min) | %A | %B |
| --- | --- | --- | --- |
| 0.0 | 0.5 | 95.0 | 5.0 |
| 6.0 | 0.5 | 0.0 | 100.0 |
| 8.0 | 0.5 | 0.0 | 100.0 |

4. Chiral HPLC Conditions

**Method A**

Column: Chiralpak AD 5 µm (250 x 4.6 mm)

Mobile Phase A: Heptane

Mobile Phase B: *I*-Propyl Alcohol

Detection: 254 nm

Method A Gradient:

| Time (min) | Flow (mL/min) | %A | %B |
| --- | --- | --- | --- |
| 0.0 | 1.0 | 90.0 | 10.0 |
| 5.0 | 1.0 | 90.0 | 10.0 |
| 20.0 | 1.0 | 50.0 | 50.0 |
| 35.0 | 1.0 | 50.0 | 50.0 |

5. HPLC Traces

**MT-106**

**HPLC**

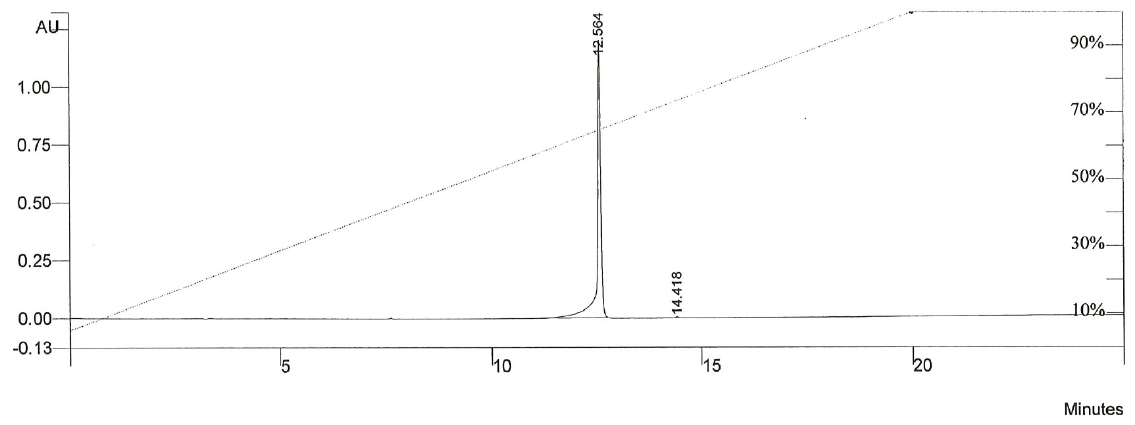

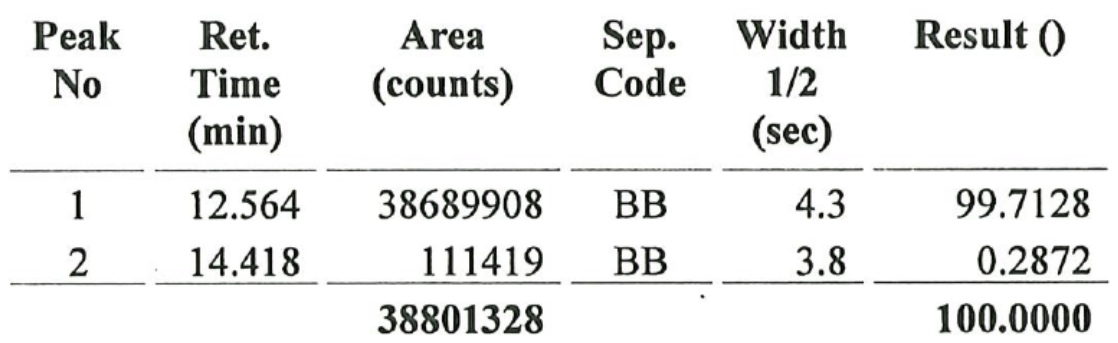

**Chiral HPLC**

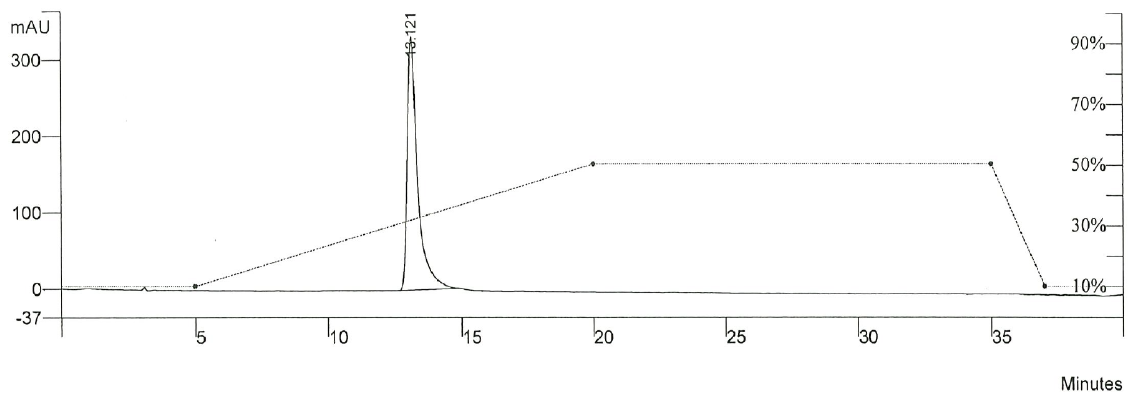

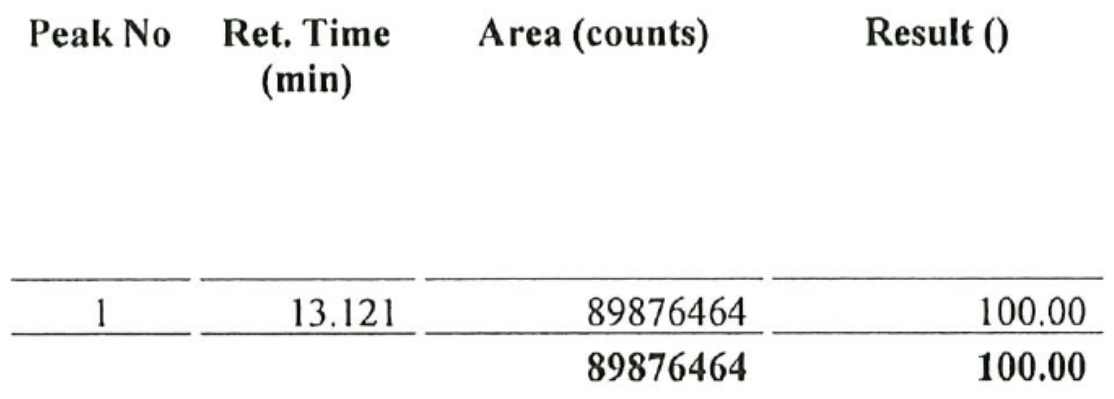

**MT-116**

**HPLC**

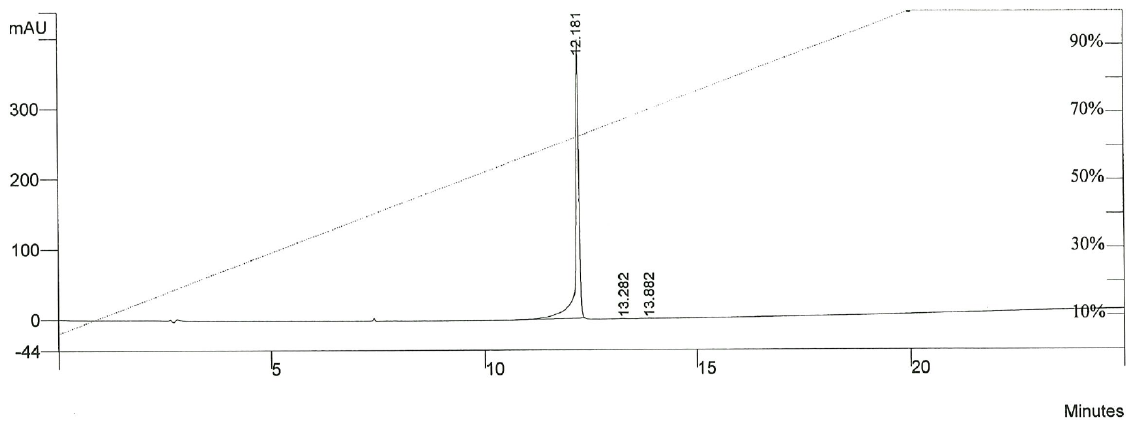

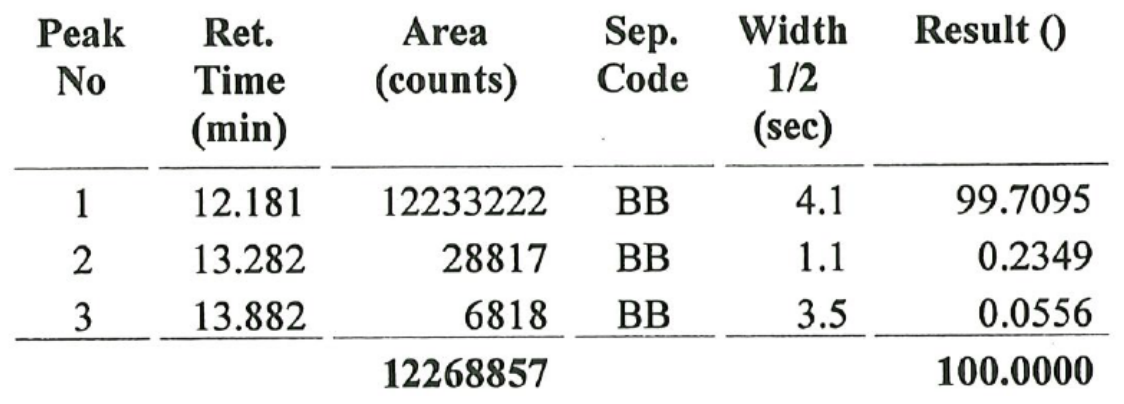

**Chiral HPLC**

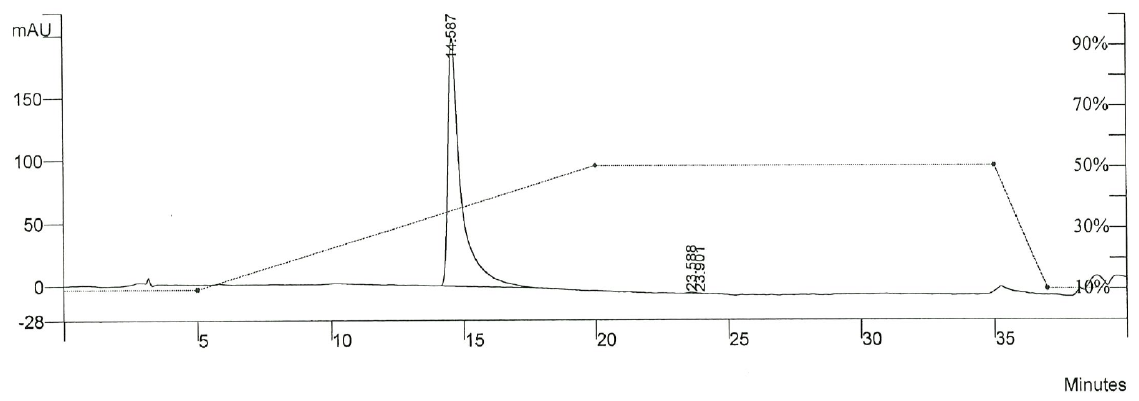

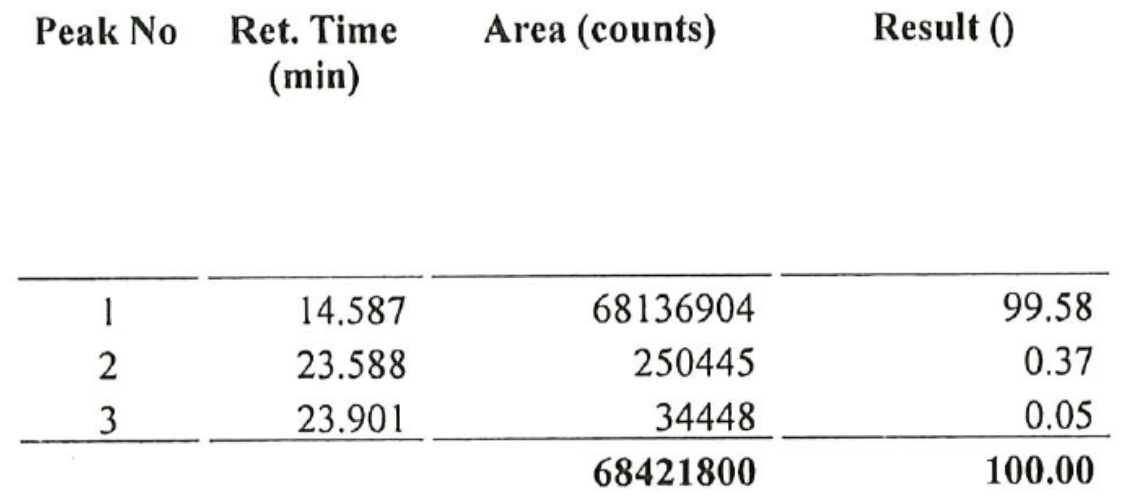

**MT-117**

**HPLC**

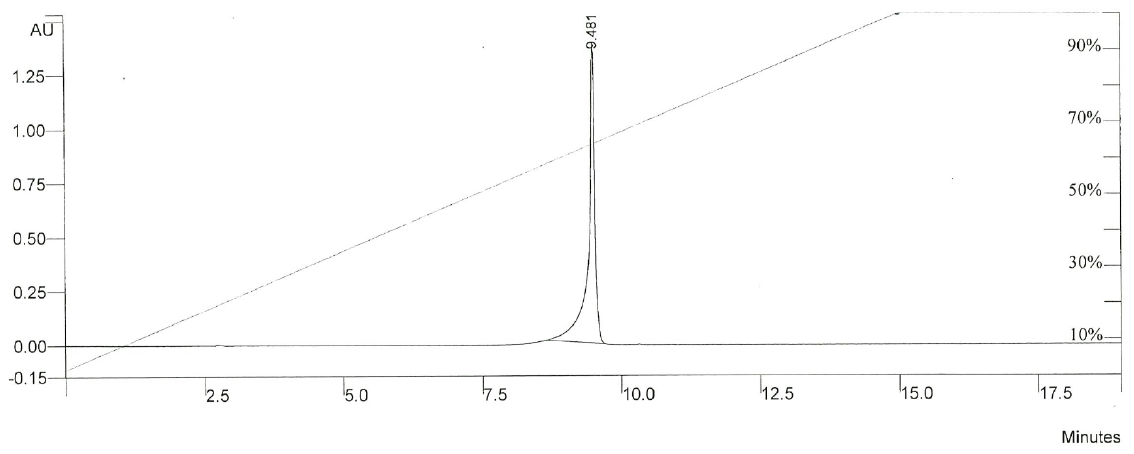

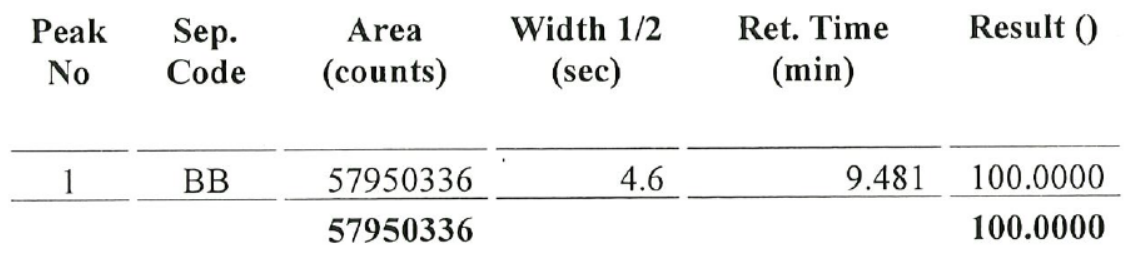

**MT-140**

**HPLC**

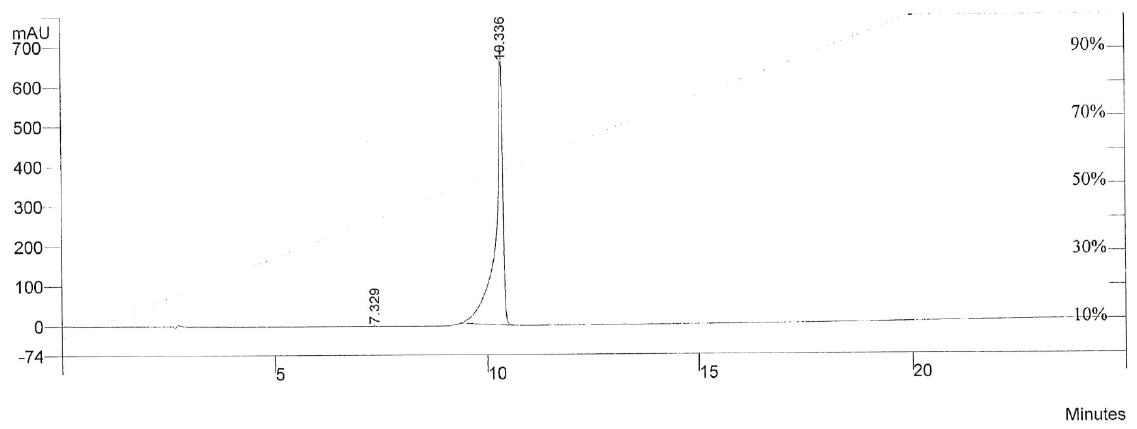

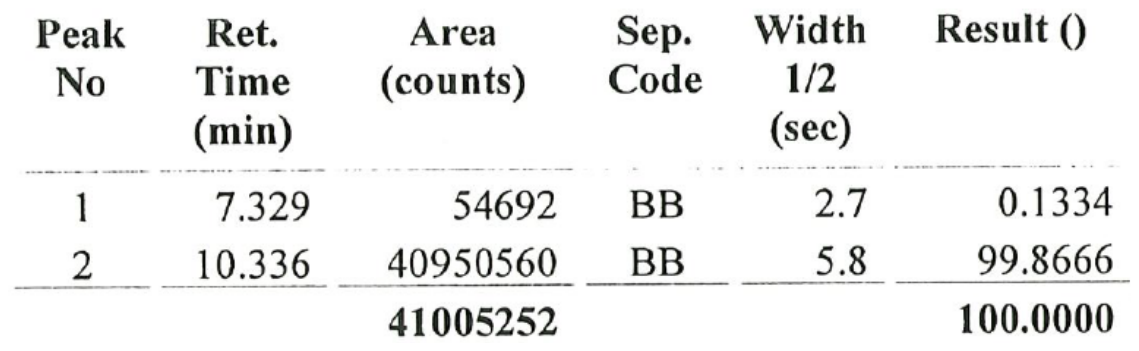

**Chiral HPLC**

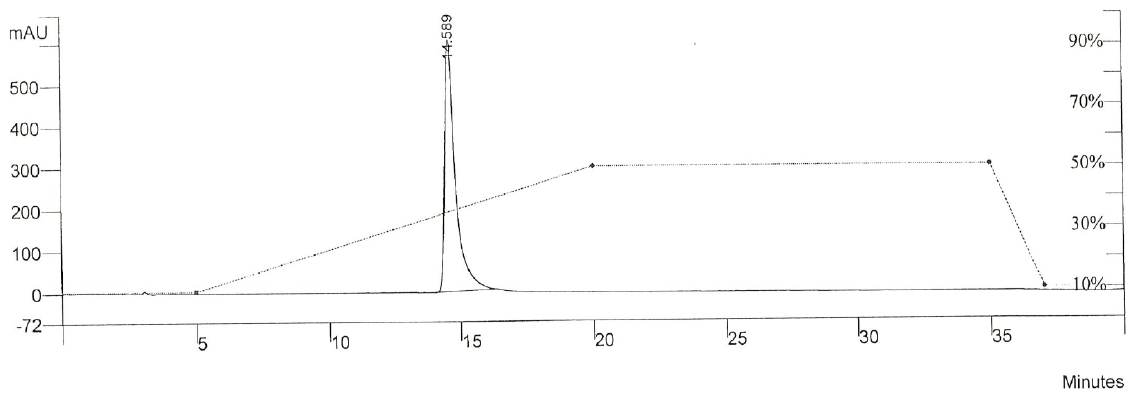

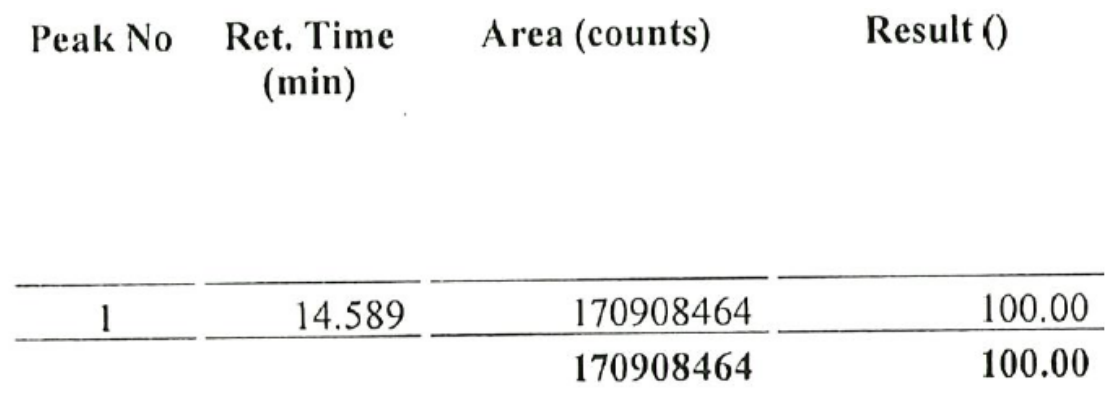

**MT-141**

**HPLC**

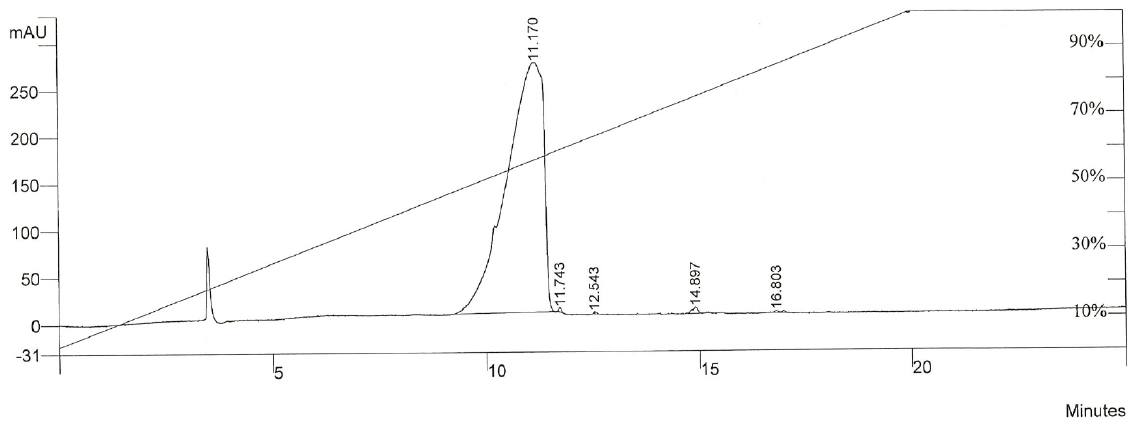

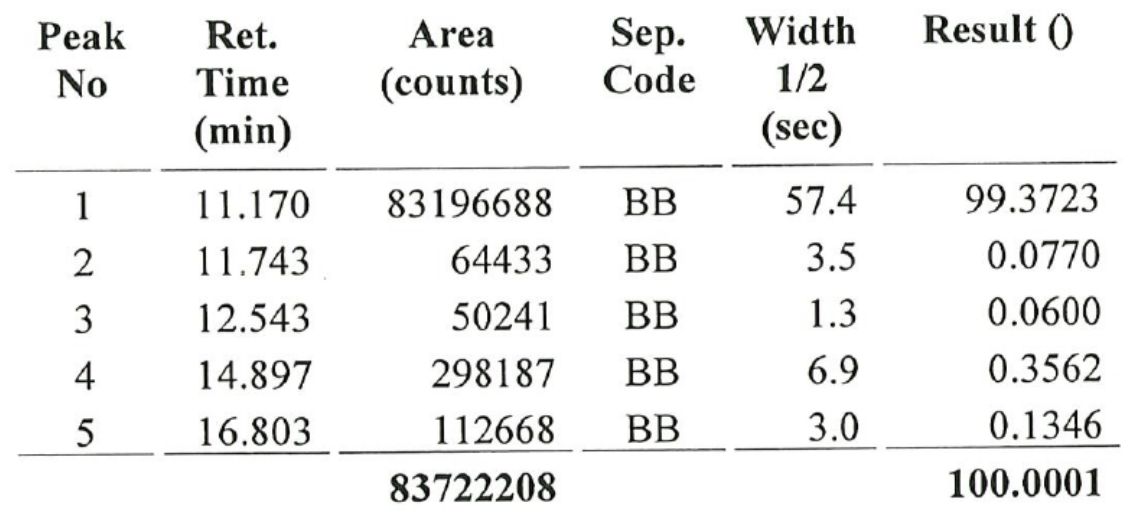

**Chiral HPLC**

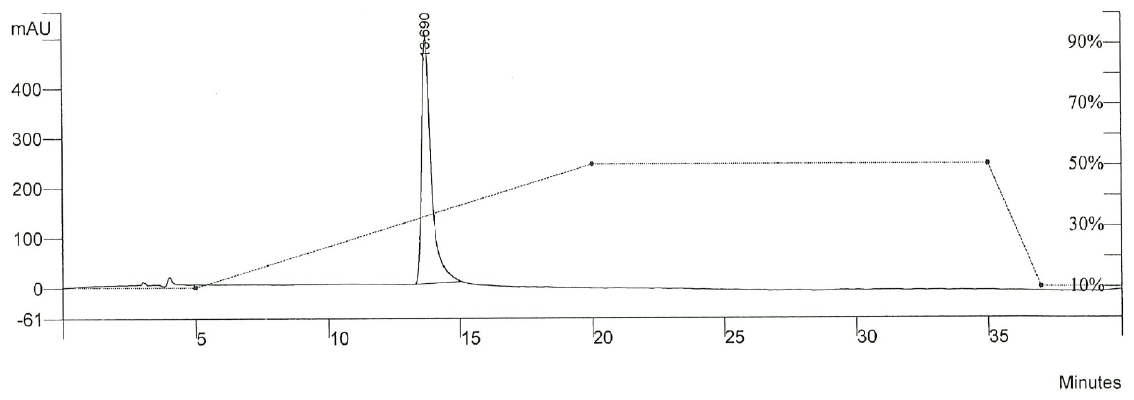

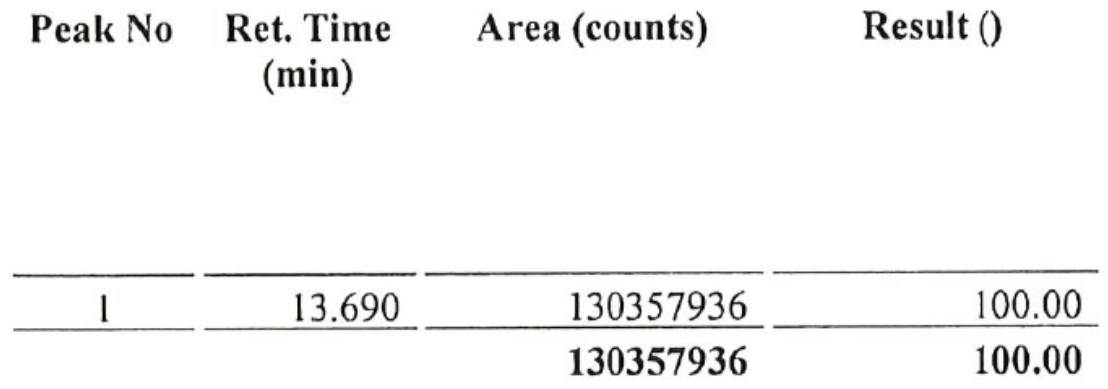

**MT-142**

**HPLC**

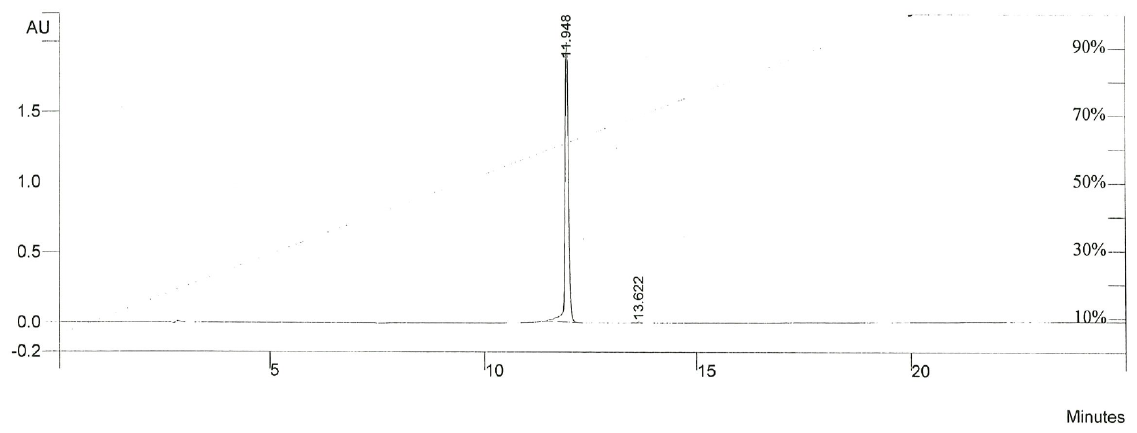

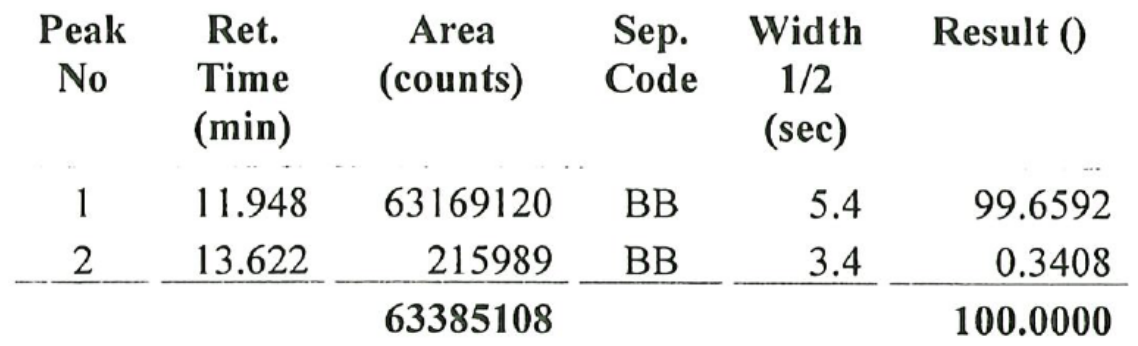

**Chiral HPLC**

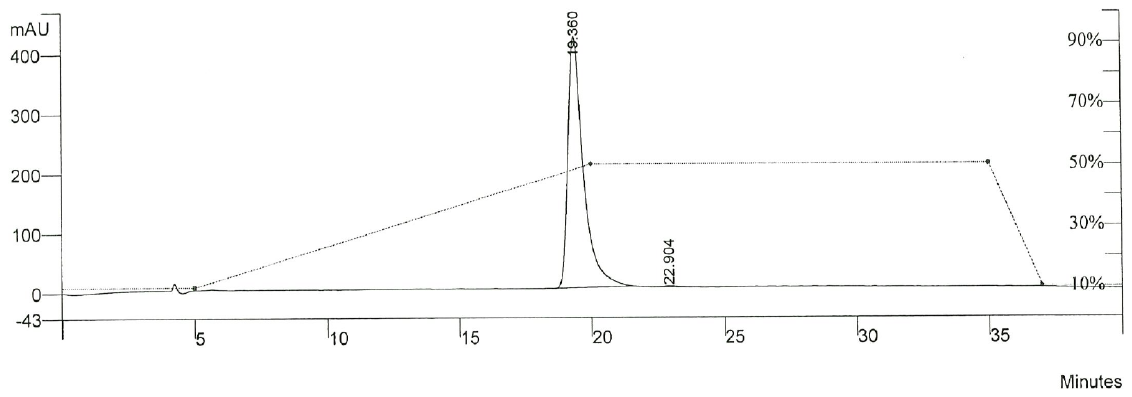

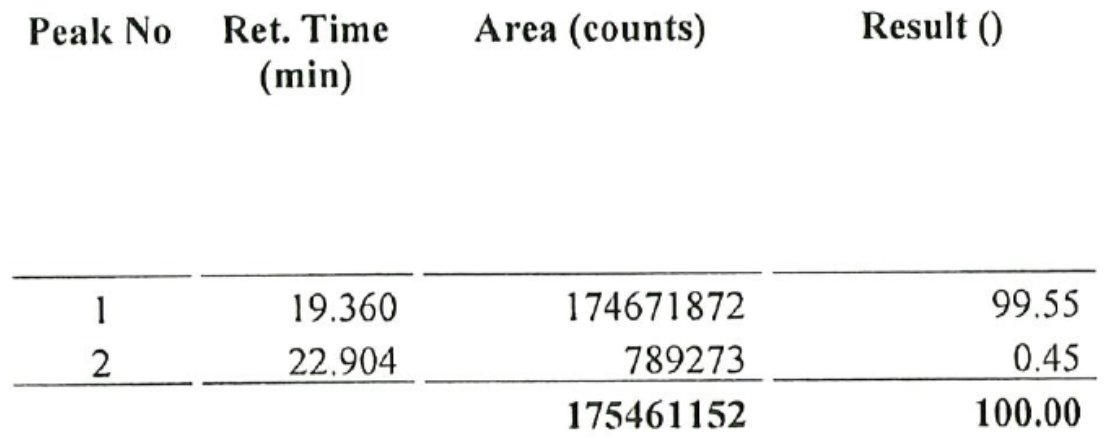

**MT-150**

**HPLC**

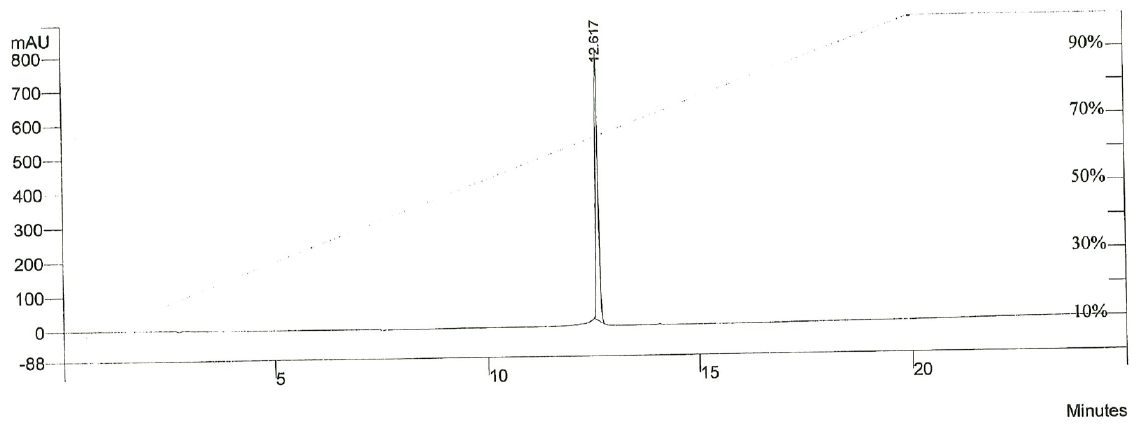

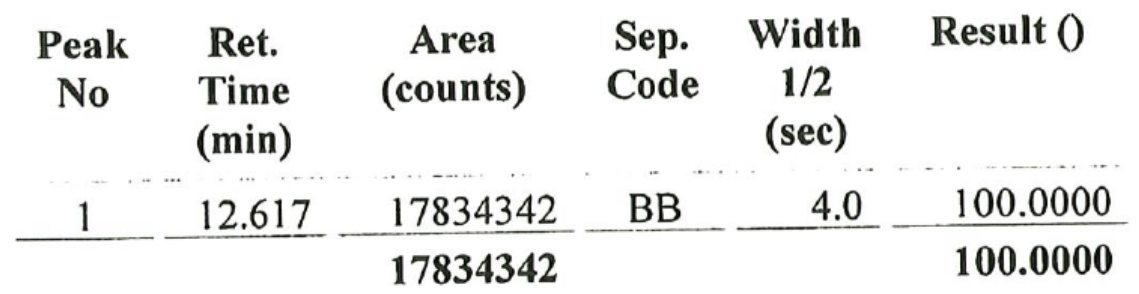

**Chiral HPLC**

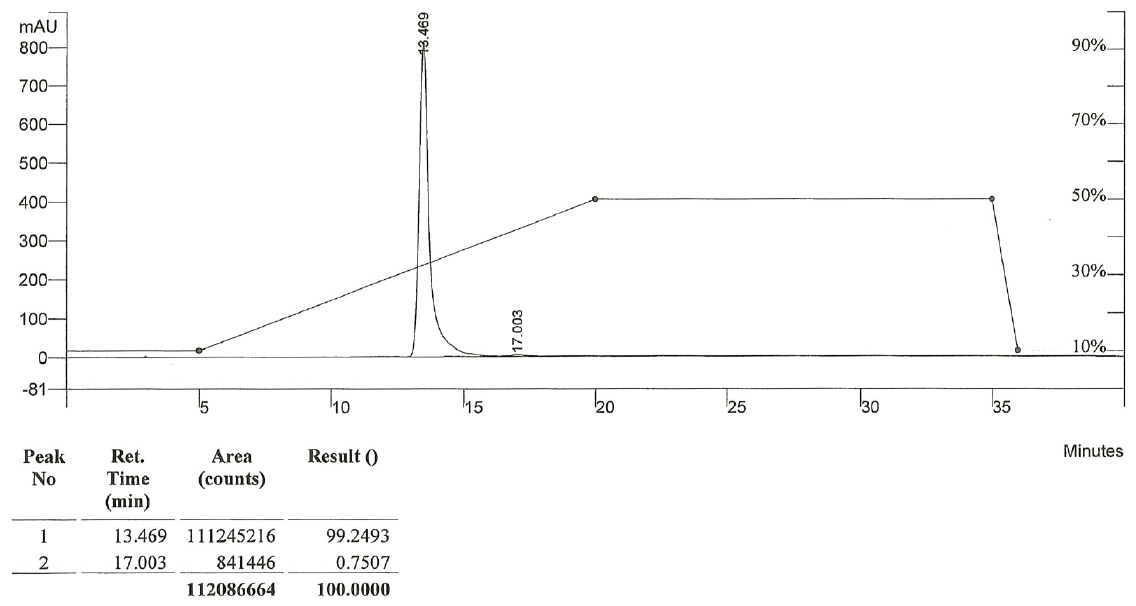

**MT-151**

**HPLC**

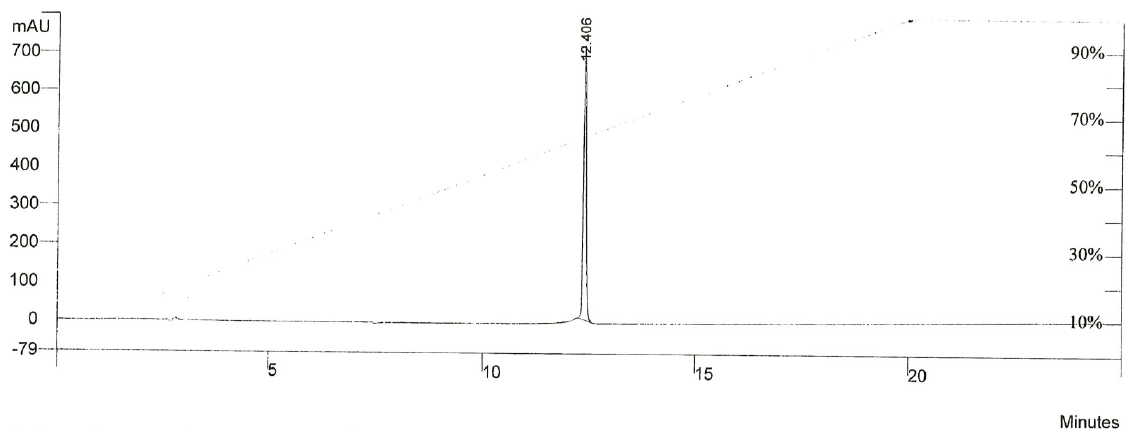

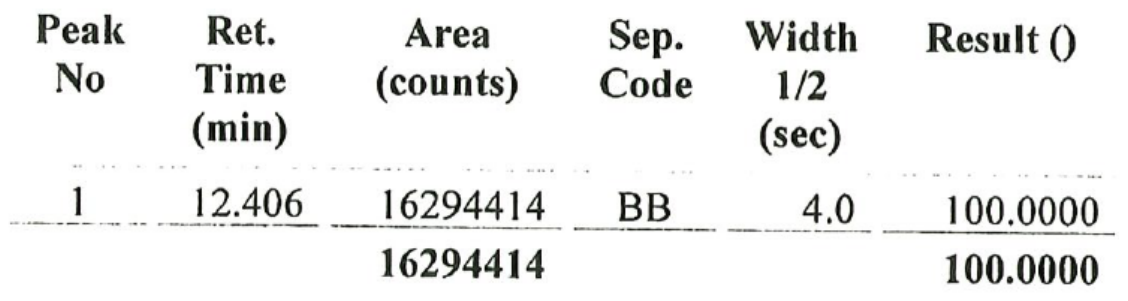

**Chiral HPLC**

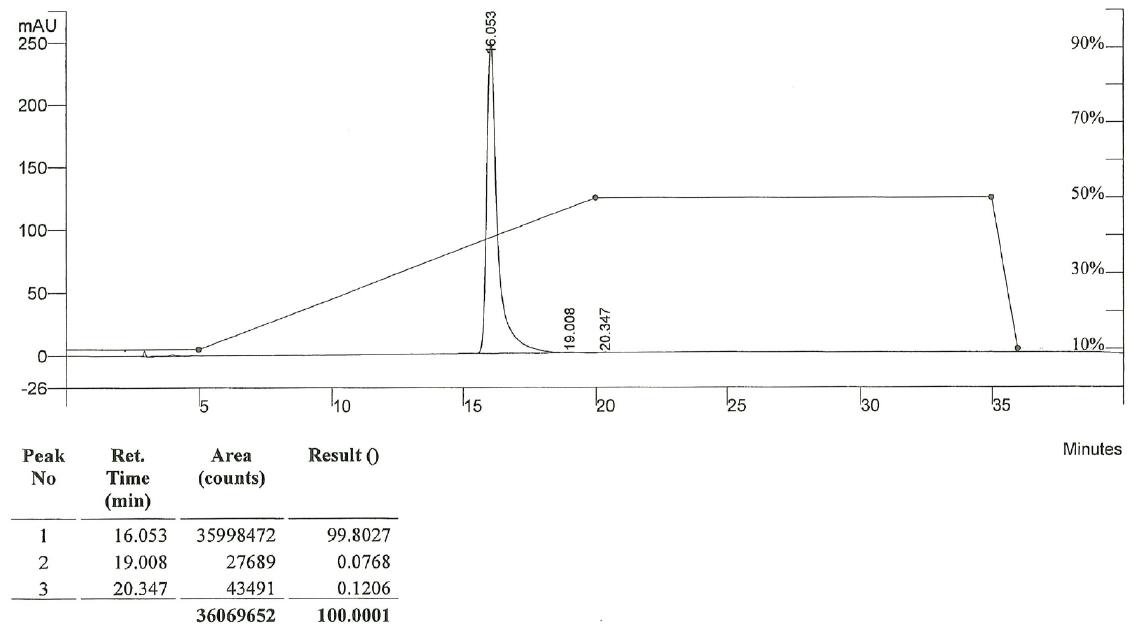

**MT-152**

**HPLC**

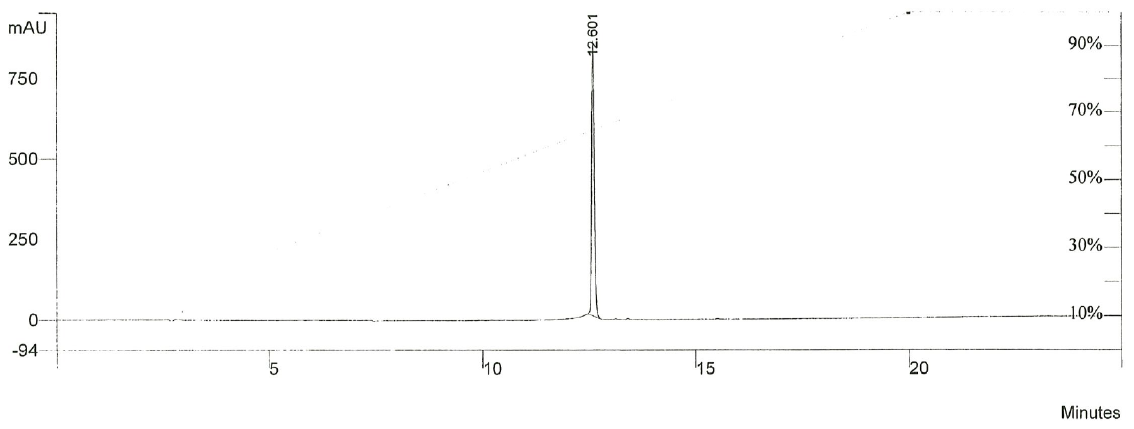

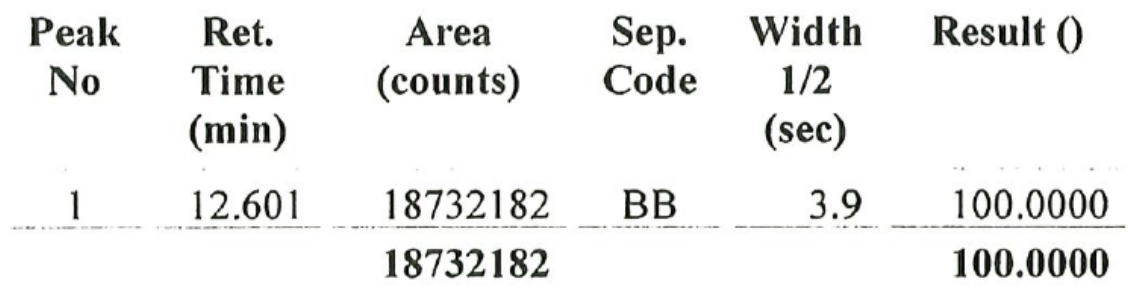

**Chiral HPLC**

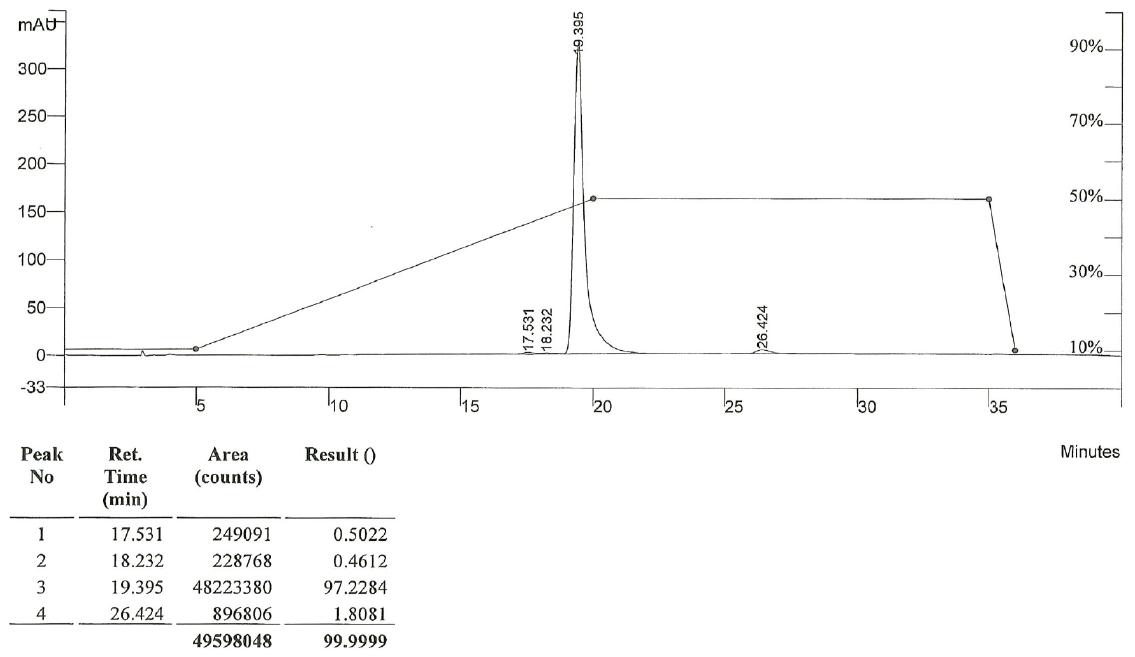

**MT-154**

**HPLC**

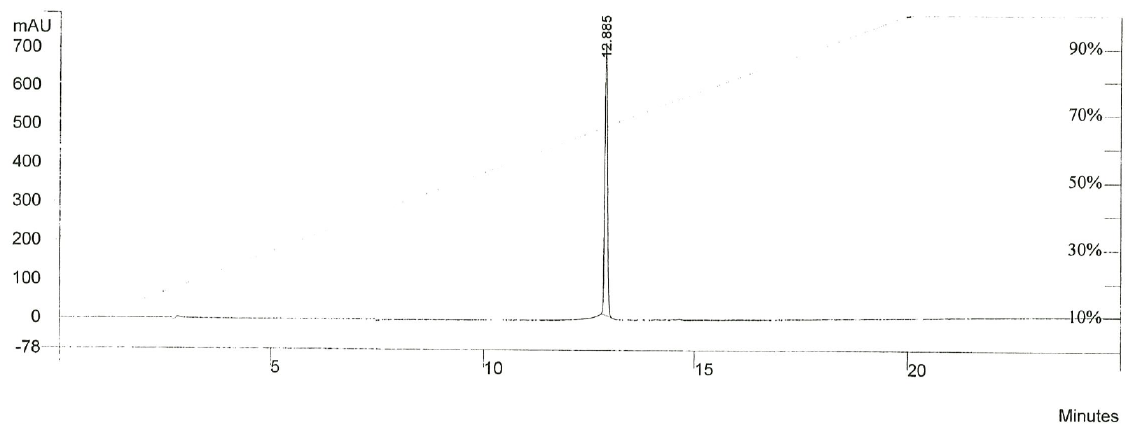

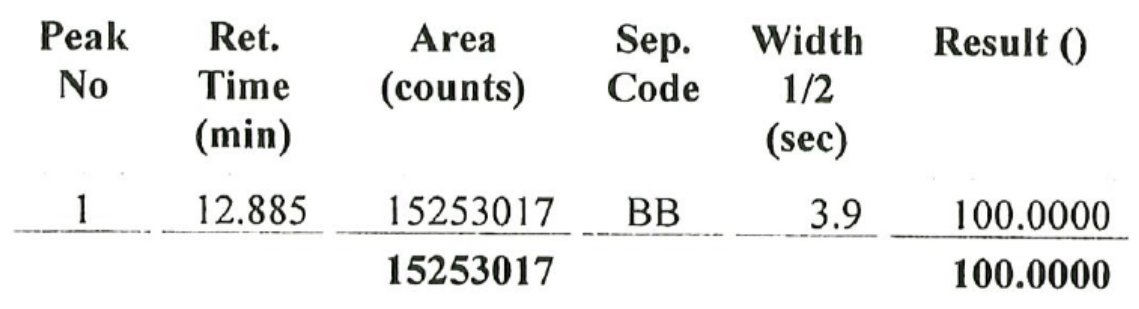

**Chiral HPLC**

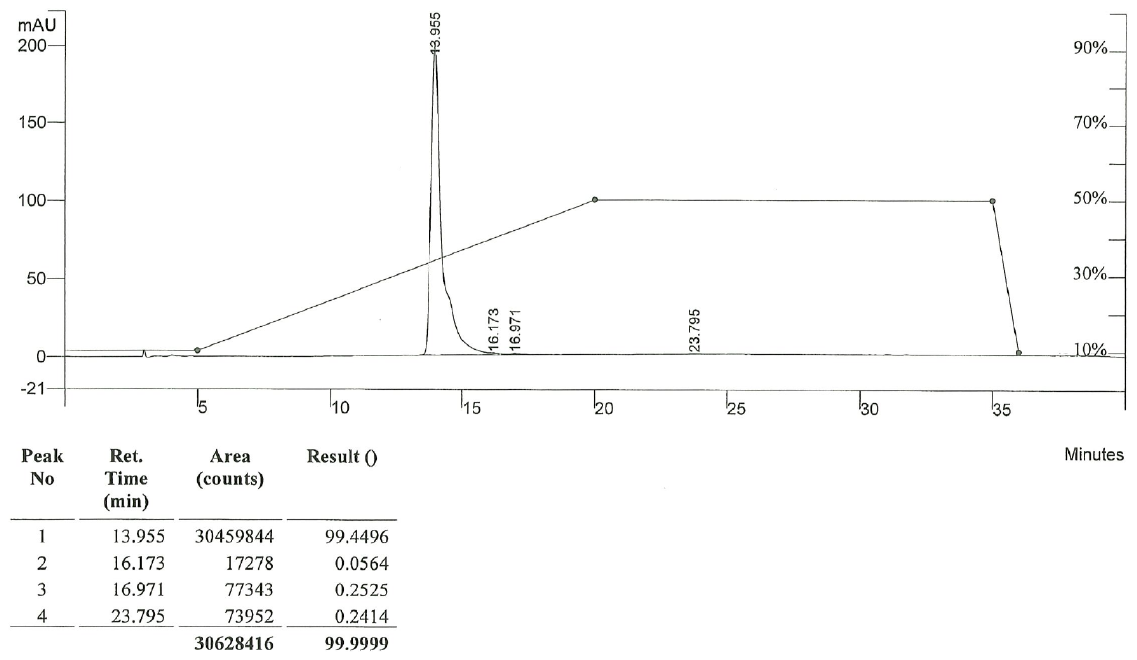

**MT-155**

**HPLC**

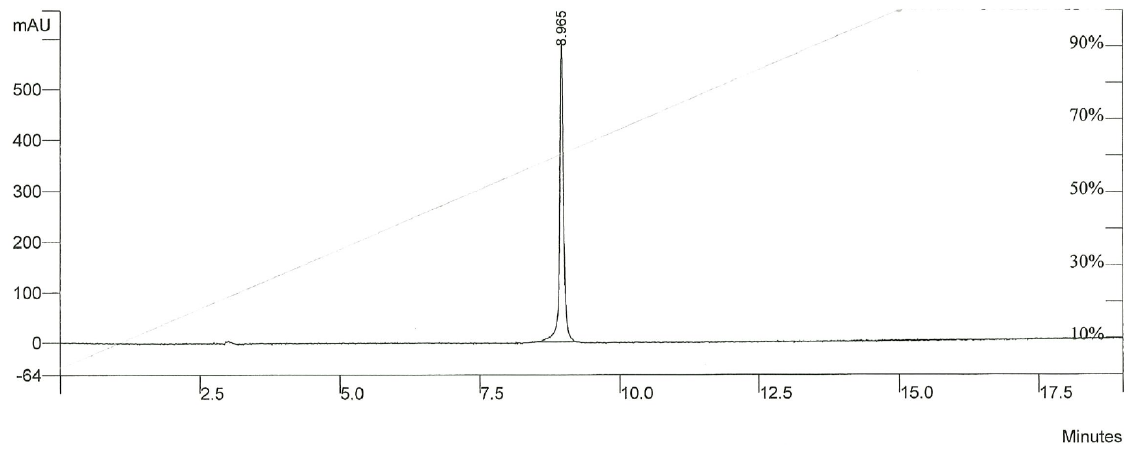

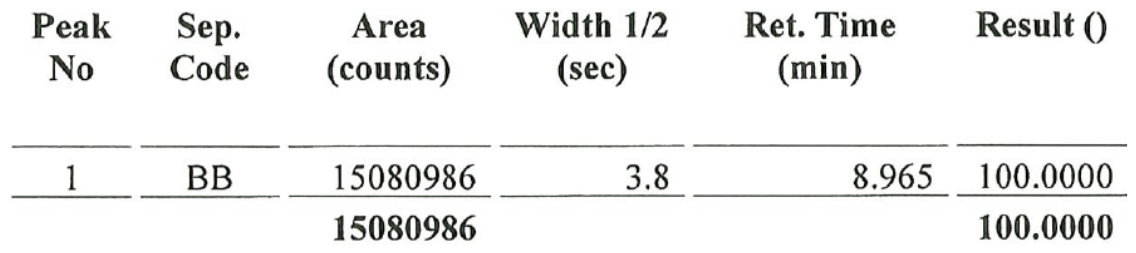

**Chiral HPLC**

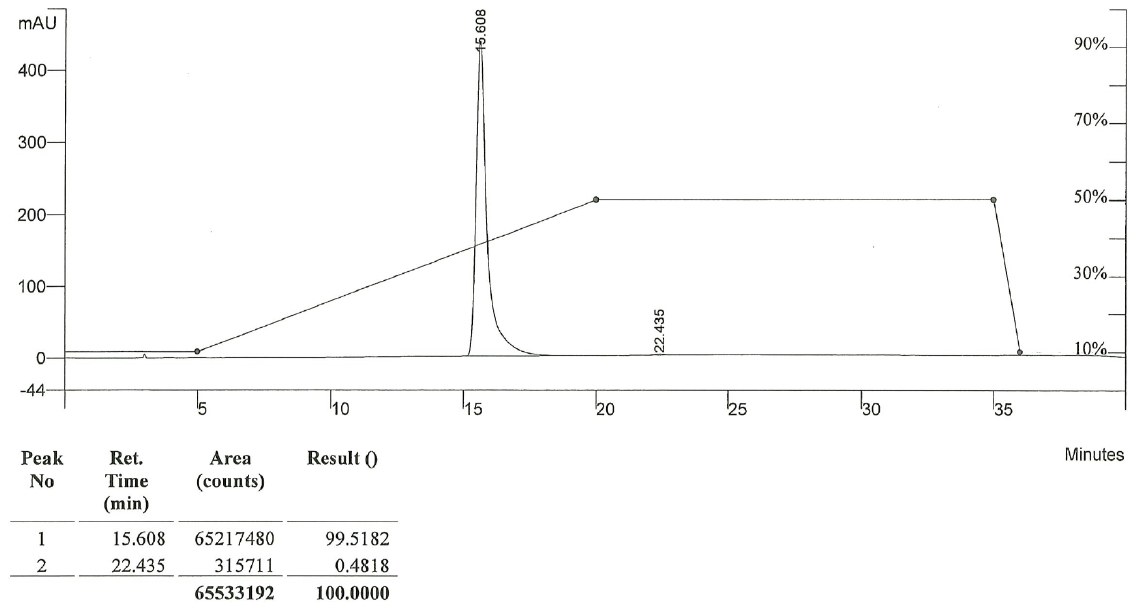

**MT-156**

**HPLC**

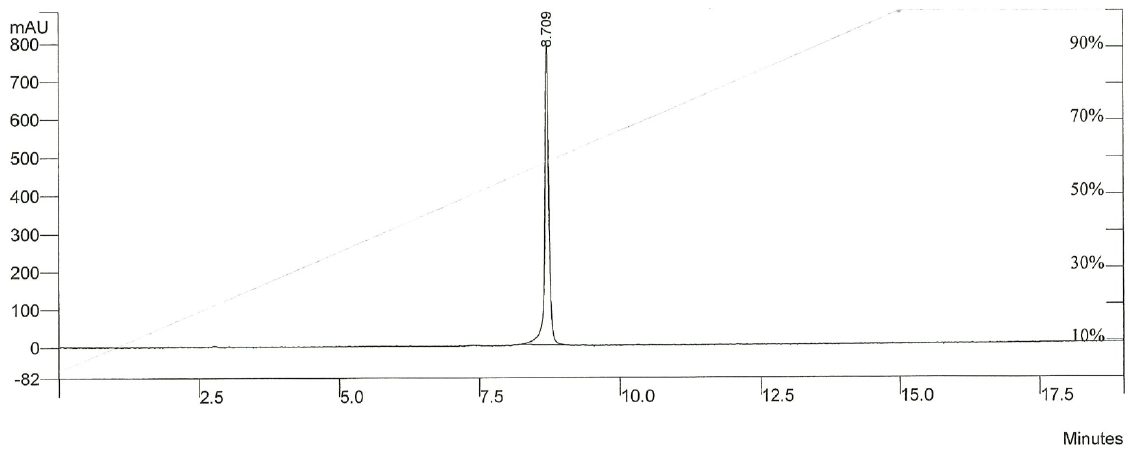

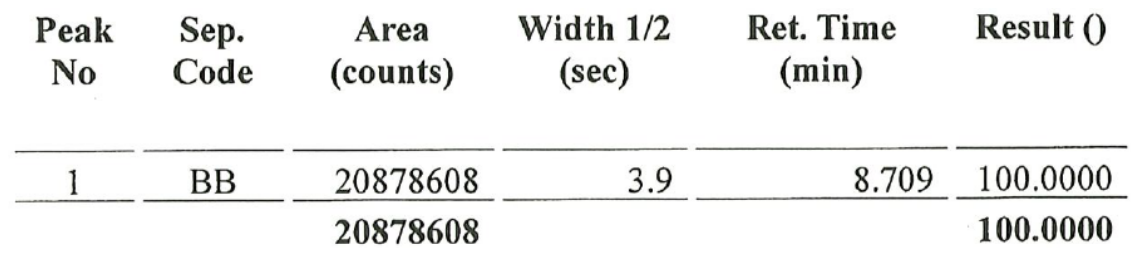

**Chiral HPLC**

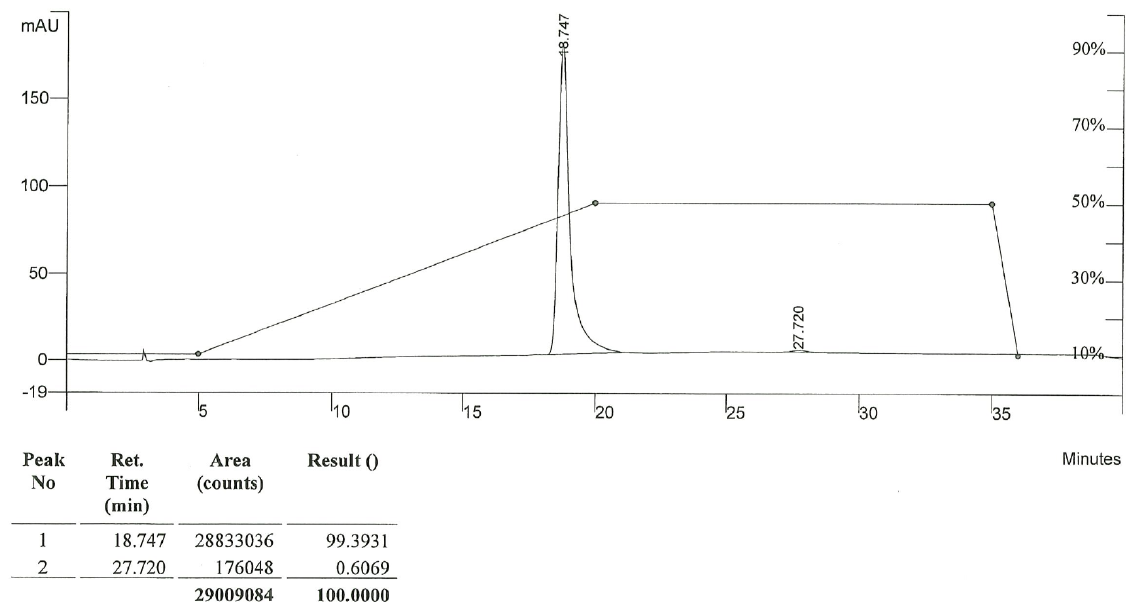

**MT-157**

**HPLC**

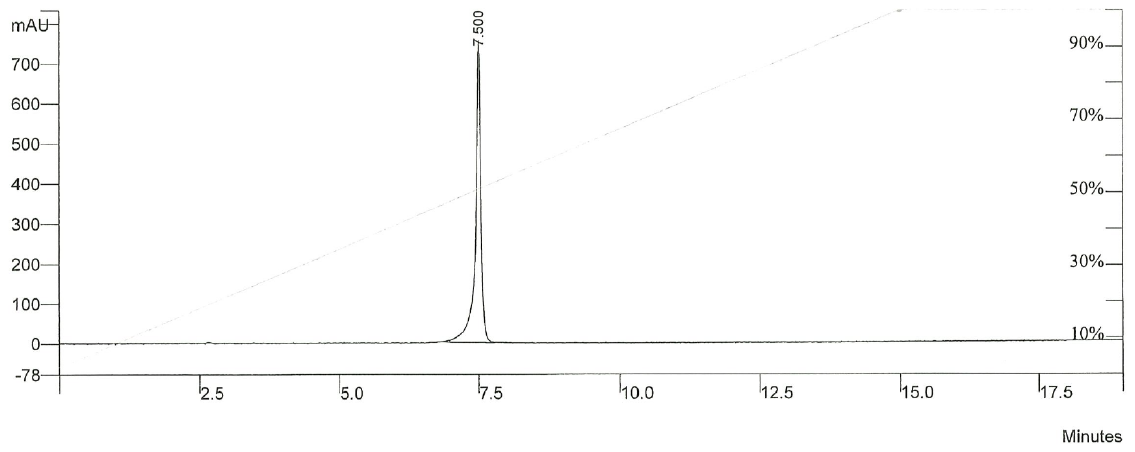

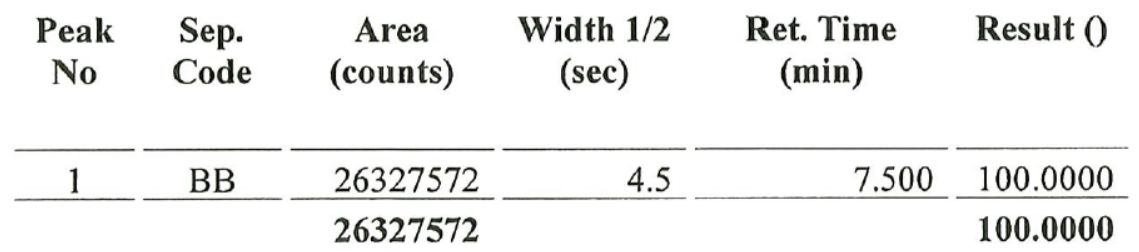

**Chiral HPLC**

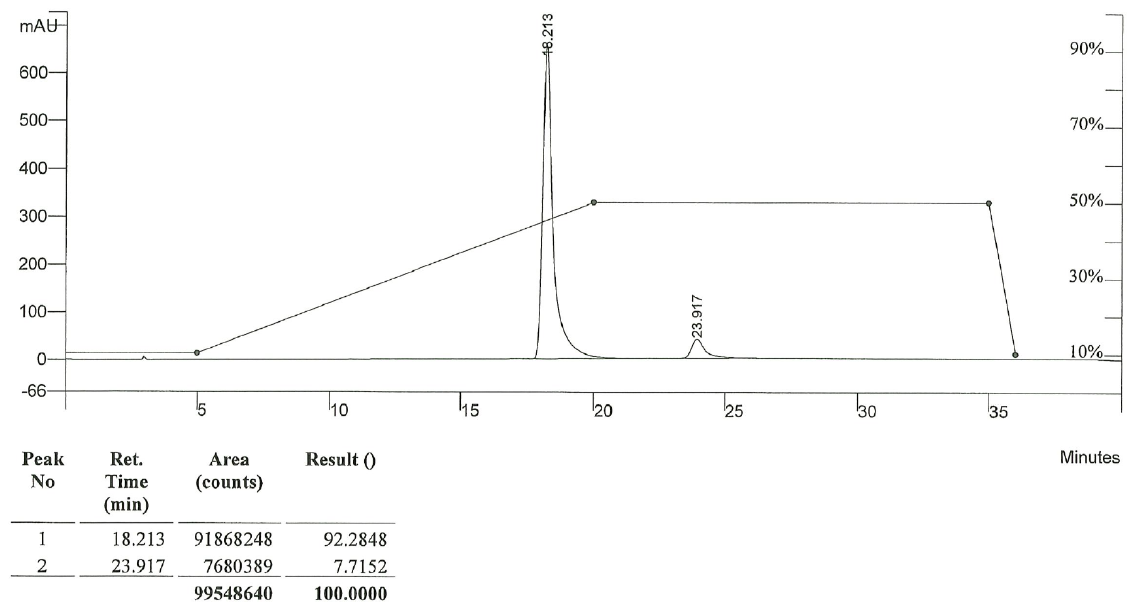

**MT-158**

**HPLC**

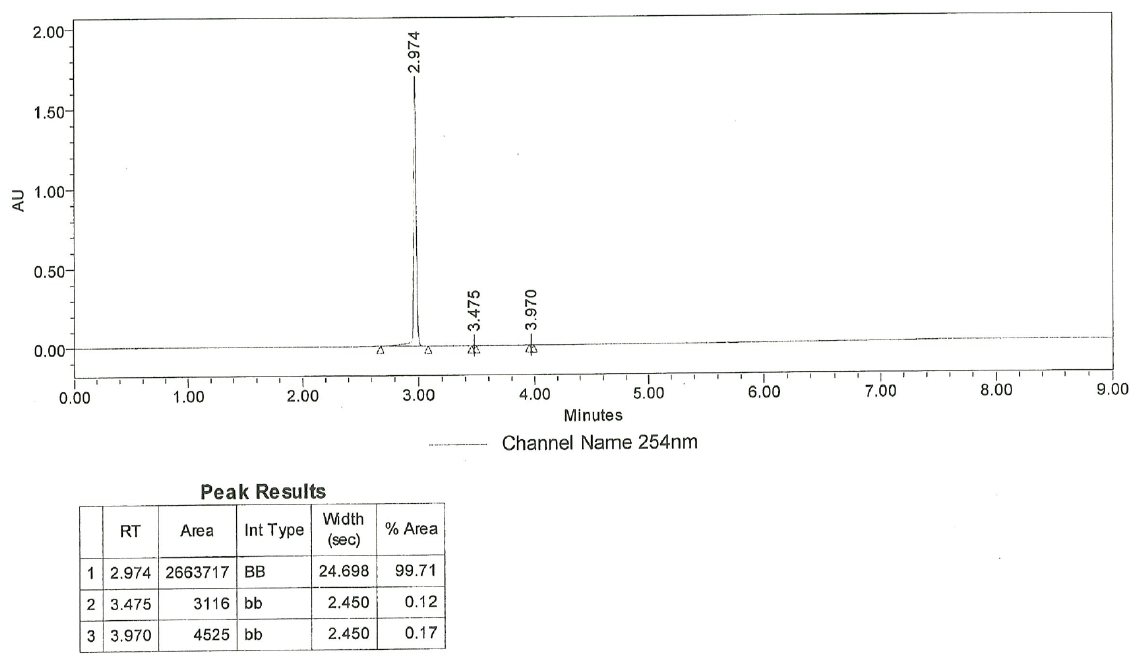

**Chiral HPLC**

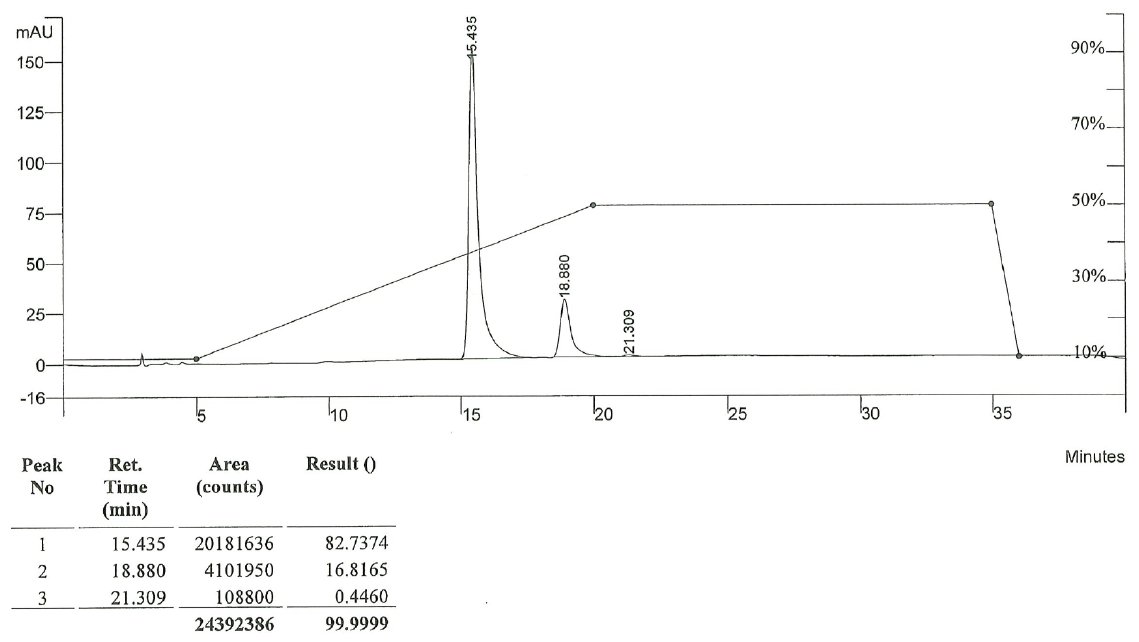

**MT-160**

**HPLC**

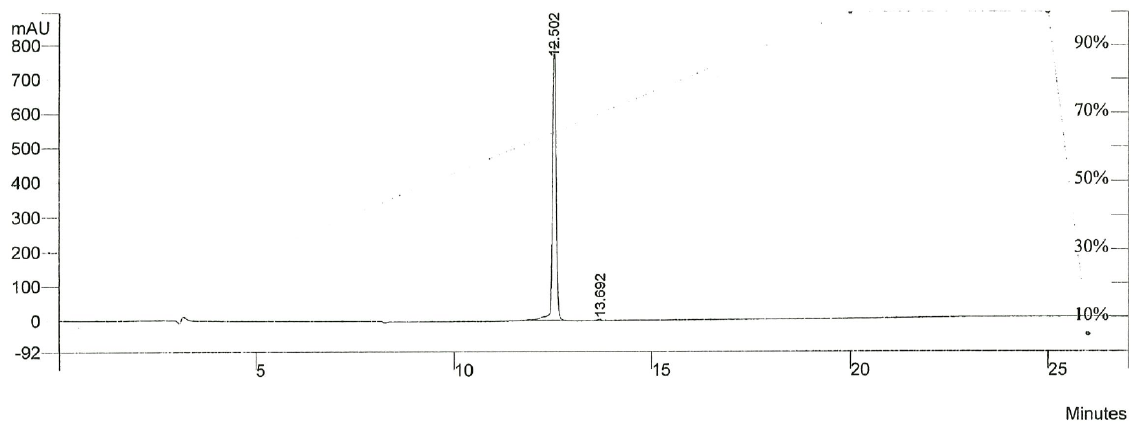

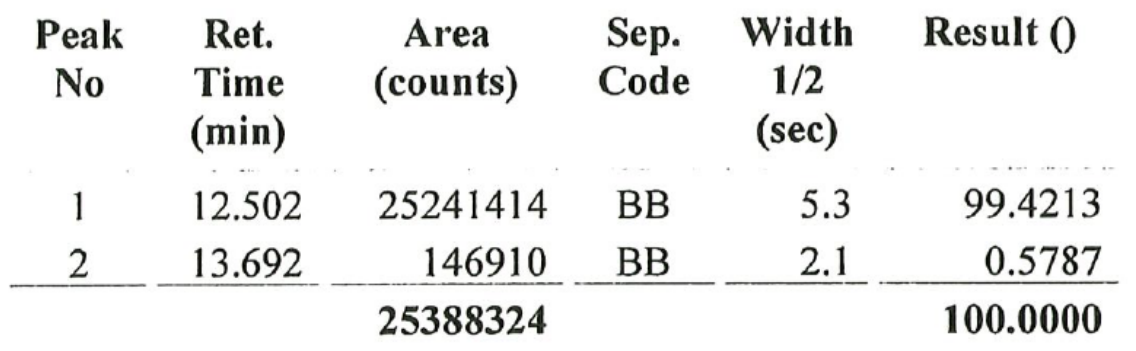

**Chiral HPLC**

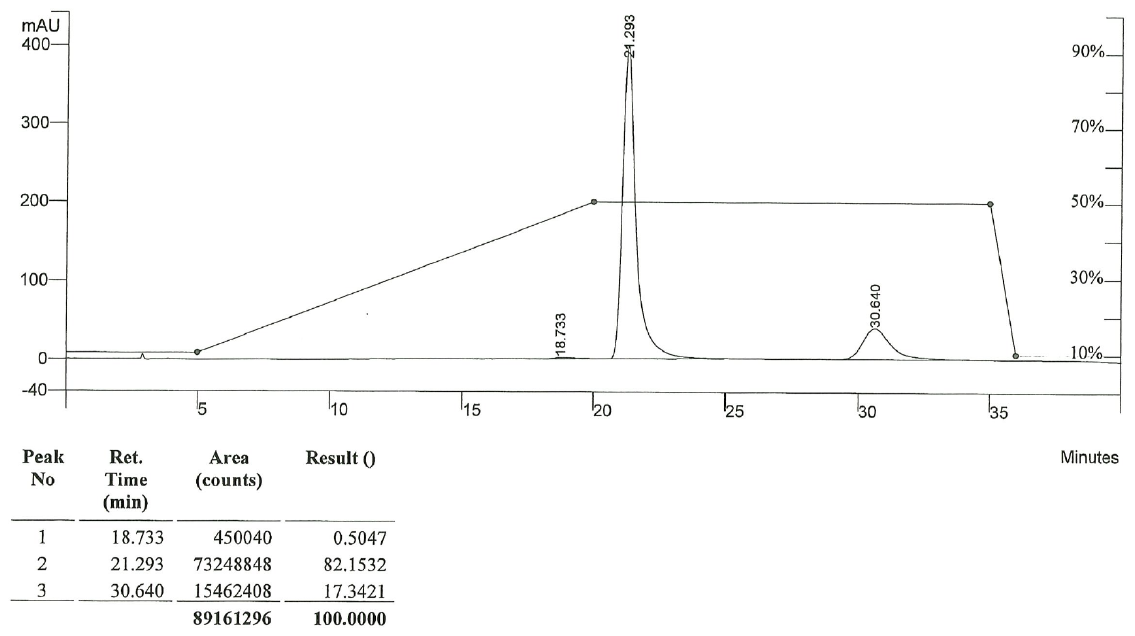

**MT-161**

**HPLC**

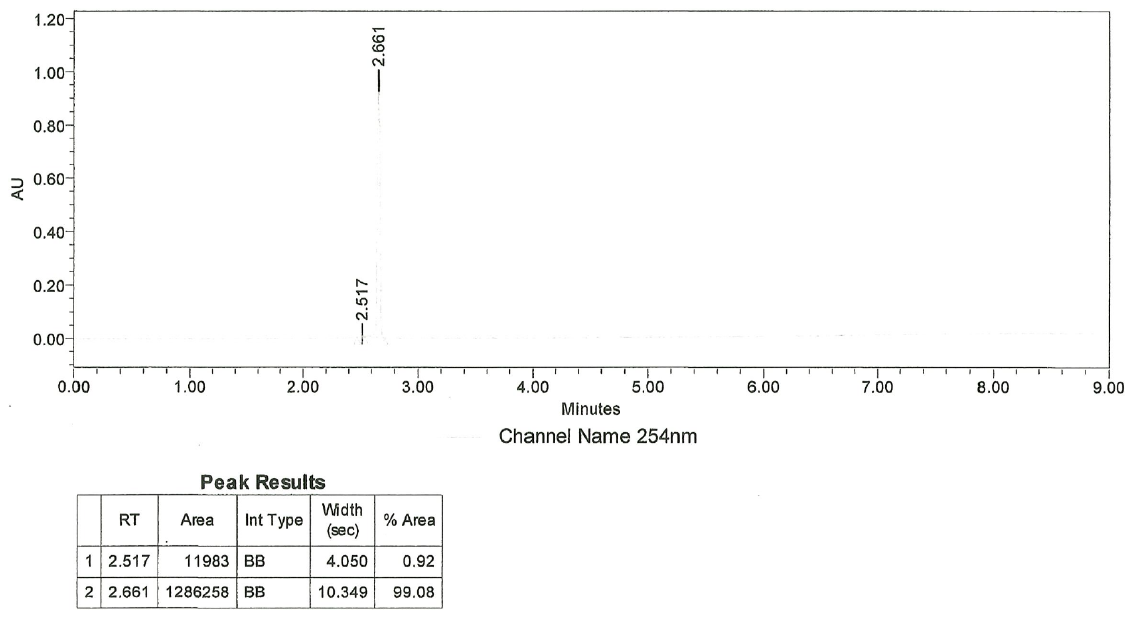

**Chiral HPLC**

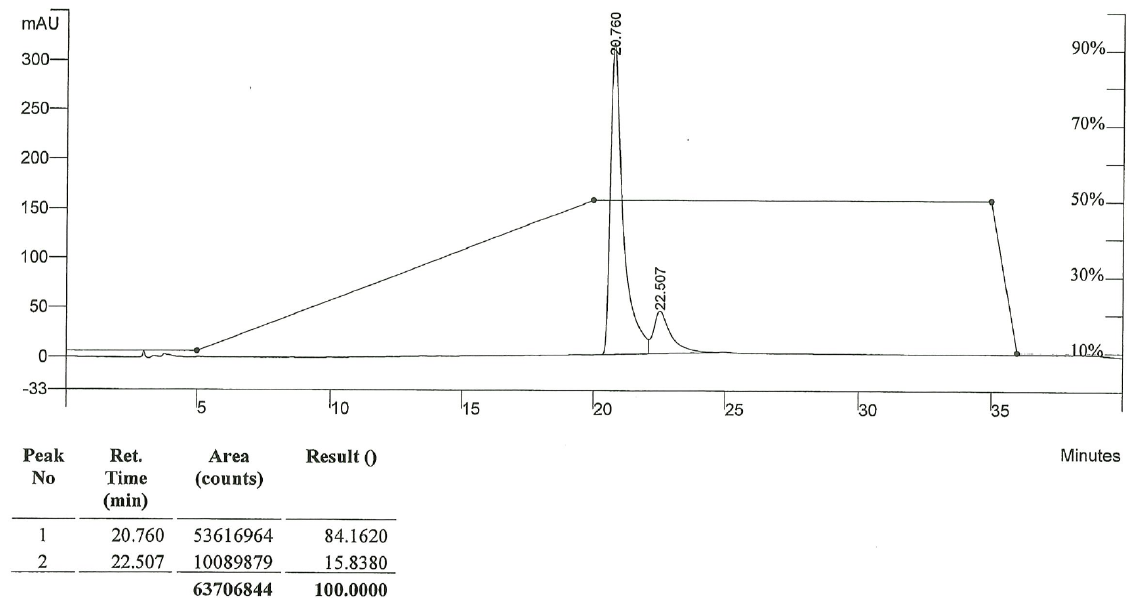

**MT-228**

**HPLC**

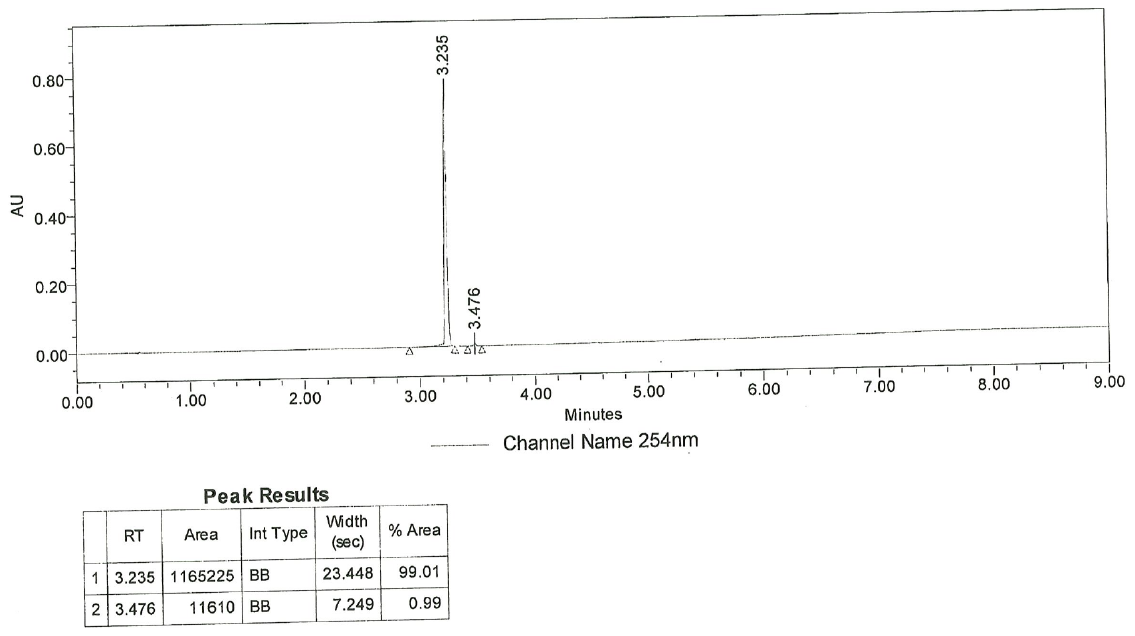

**Chiral HPLC**

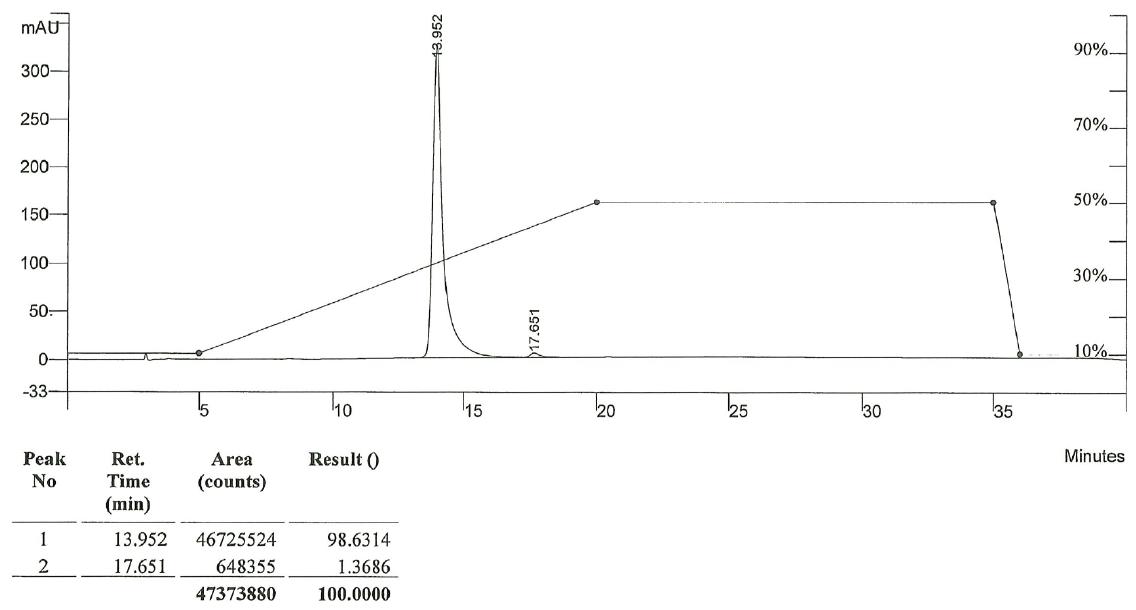

General Methods

**Kinetic aqueous solubility.**

Kinetic aqueous solubility of blebbistatin derivatives was determined as described previously (*3*).

**Photostability**

The photostability of blebbistatin derivatives was determined as described previously (*3*).

**Protein Isolation and Purification**

NMIIA was isolated from human platelets and thiophosphorylated as described previously (*3*). Bovine CMII (cat. MY03), rabbit SkMII (cat. MY02), and the S1 fragment of chicken gizzard SmMII (cat. CS-MYS05) were obtained from Cytoskeleton, Inc. F-actin was prepared from rabbit muscle acetone powder (cat. 41995-2, Pel Freez Biologicals) according to the protocol of Pardee and Spudich (*17*). The full length recombinant human non-muscle myosin IIB was co-expressed with myosin regulatory and essential light chains in the baculovirus/Sf9 system and thiophosphorylated as described previously (*3, 18*).

For structural studies, Sf9 cells expressing human SmMII with a C-terminal Flag tag were mechanically lysed by 7 shots in a Dounce homogenizer in buffer: 200 mM NaCl; 20 mM HEPES pH 7.5; 4 mM MgCl_2_; 0.5 mM EDTA; 1 mM EGTA; 0.5% Igepal; 7% Sucrose; 1 mM NaN_3_; 1 mM PMSF; 10 μg/mL Aprotinin; 10 μg/mL Leupeptin; 5 mM DTT; 2 mM ATP. The lysate was centrifuged at 20,000 rpm for 45 min in a 25.50 rotor. The supernatant was incubated with anti-Flag epitope antibody affinity resin under stirring for 2 h at 4 °C. The anti-flag resin was then transferred to a column. The resin was washed with 200 mL of buffer: 150 mM KCl; 20 mM imidazole pH 7.5; 5 mM MgCl_2_; 1 mM PMSF; 3 mM DTT; 1 mM EDTA; 1 mM EGTA; 10 μg/mL aprotinin; 10 μg/mL leupeptin; 3 mM ATP. Purified myosin was eluted via Flag peptide competition. The sample was then ultracentrifuged at 78,000 rpm at 4 °C for 15 min in a TLA110 rotor to spin out any actin that could still be bound to the myosin. The protein was then injected in a Superdex 200 HR 16/60 column (Cytiva) previously equilibrated in 10 mM HEPES pH 7.5; 50 mM NaCl; 2.5 mM MgCl_2_; 0.5 mM ATP; 1 mM DTT; 1 mM NaN_3_ and concentrated by ultrafiltration. The purity of all myosin preparations was confirmed by SDS-PAGE.

**NADH-coupled ATPase assays**

The potency of blebbistatin derivatives against NMIIA, CMII, SKMII, and SmMII was assessed in a nicotinamide adenine dinucleotide (NADH)-coupled ATPase assay as described previously (*2, 3*)

**Cytokinesis Assay**

The potency of Blebb derivatives on NMIIB was determined in a COS7 cell-based cytokinesis assay as described previously(*1, 3*).

**Animals**

Adult, 8-10-week-old male C57BL/6 mice (25-30 g, Jackson Laboratory) were housed under a 12:12 light/dark cycle, with food and water ad libitum. Adult male 250-300 g rats (Charles River) at the start of the experiment were handled and housed under a 12:12 reverse light/dark cycle, as previously described (*19, 20*). Rats used for self-administration had unlimited water access, but were food restricted (20-24 g rat chow/day) throughout experiments except during surgery and recovery. Unless otherwise noted, rats and mice were handled for three days before behavioral testing. All procedures were performed in accordance with the Scripps Research Institutional Animal Care and Use Committee.

**Drugs**

For behavioral injections, methamphetamine hydrochloride (METH; 2 mg/kg, IP for mice, and 0.02mg/0.05ml/infusion, IV for rats; Sigma-Aldrich) or cocaine hydrochloride (COC; 15 mg/kg, IP for mice; NIDA) was dissolved in sterile 0.9% saline. The hydroxypropyl-b-cyclodextrin (HPbCD) vehicle was made by dissolving 9 g of HPbCD powder to 21 ml of 100mM Acetate Buffer, pH 5.4. Then, prior to testing, the 840 μl of HPbCD vehicle was mixed with 60 μl of fresh DMSO. Blebb or its derivatives (MT-106, MT-152, MT-228) were dissolved in DMSO to achieve the desired concentration then 60 μl of the DMSO-based compound stock solution and 840 μl HPbCD vehicle were mixed immediately prior to injection, as previously described (*20-22*). Treatment doses of racemic (+/-) Blebbistatin (Blebb; Tocris), Blebb derivative or Vehicle were administered via either IP or IV tail vein for mice and intrajugular vein catheter for rats. siRNA against MyH10 (ON-TARGETplus SMARTpool, mouse Myh10 or nontargeting pool; Dharmacon) using JetSI transfection reagent as previously described (*20*). JetSi was infused into the BLA 24 hrs prior to testing at 1 µL at 200nL/min, based on previous research (*19, 20*). Infusers were left in place for 1 min for diffusion. After behavioral procedures, verification of injector tip location was performed on cresyl violet-stained coronal sections.

**Surgery**

For self-administration experiments, rats were implanted with chronic intravenous catheters assembled in house (*19*) or purchased (P1 Technologies, assembled by Access Technologies; *23*), surgery was conducted as previously described (*19, 23*). Rats were given one week for recovery before being handled for three days. Catheter patency was maintained with daily IV infusions of heparinized saline (~0.2 cc of 30-60 i.u./ml), and 0.1 mg/ml cefazolin to prevent infection.

For one fear conditioning experiment, mice were deeply anaesthetized with isoflurane and implanted bilaterally with 26G guide cannula (P1 Technologies) targeting the BLA (AP: 1.5 mm, ML: +/-3.2 mm from bregma, and DV: -4.7 mm from the skull) as previously described (*19*). Animals were allowed to recover for a week prior to undergoing fear conditioning.

**Behaviors**

*Conditioned Place Preference (CPP)*

Mouse: CPP was conducted similarly as previously described (*19, 20, 24*). Briefly, mice underwent one or two 30 min pretests, during which mice were allowed to freely explore all three CPP chambers (Med Associates; black, 3 lux lighting, McCormmick coffee extract; middle grey, 15 lux lighting, no scent; white, 2 lux lighting, McCormmick orange extract; all three chambers had uniquely textured floors). The final 15 min of the pretest were used to measure bias and to counterbalance groups for which chamber mice received drug (CS+) or saline (CS-). Mice did not have a significant preference for the white or black chamber (two-way ANOVA chamber x group: METH CPP F_(4,53)_ = 0.44, *P* > 0.05; COC CPP F_(1,14)_ = 0.07, *P* > 0.05). However, mice that spent more than 70% of the total time in one chamber were excluded from the study (n = 5). Mice underwent conditioning over four consecutive days, with twice daily 30 min conditioning sessions separated by at least 4 hr, alternating injections of saline or drug (METH or COC) to receive 4 pairings each. In some experiments, mice were tested for memory retention during 15 min free access sessions, similar to pretest, 48 hr after the final conditioning session, as previously done (*19, 21*). Vehicle or treatment injections (IP or IV tail vein) were administered 30 min before testing. Mice were injected with saline (IP) before pre- and post-tests, to mimic conditioning sessions procedures. All testing and conditioning were conducted in red or no light, and with a white noise generator (San Diego Instruments) set at ~65-70 dB below the CPP apparatus.

*Self-Administration*

METH self-administration procedures were conducted using 16 sound attenuating operant HABITEST conditioning chambers (Coulbourn) as previously described (*19, 20*). Briefly, one week after arrival, rats were food restricted for three days prior to a ~20 hr food (45 mg grain pellet, Bioserv) training session on a fixed ratio-1 (FR1) schedule of reinforcement. One week after surgeries, rats were trained to self-administer METH during daily 2 hr sessions on a FR1 schedule for 14 sessions, as previously described (*19, 23*). Briefly, acquisition was conducted in a unique context composed of dark blue Plexiglas floors with the house light on and McCormick’s vanilla extract. Responding on the active (right) lever resulted in the initiation of an infusion of METH and was followed by a 20 s timeout during which no lever presses were reinforced. All lever presses, reinforced or not, are reported as active lever presses during acquisition and testing. Responding on the inactive lever was recorded but had no programmed consequences. Rats that did not maintain an average of 10 infusions during acquisition were excluded. In some experiments, drug-associated memory retention was assessed four days following the last day of acquisition during a 1 hr drug free session (Test 1). Memory retention was tested again (Test 2) 14 or 30 days following the final acquisition day. Groups were matched on total number of infusions received during self-administration, the average number of infusions and total active and inactive lever presses made over the last three days of self-administration.

*Fear Conditioning and Extinction*

Auditory Fear Conditioning: As previously described (*19*), mice were habituated to modified Noldus PhenoTypers chambers inside of sound attenuating boxes during three 4 min exposures to the fear conditioning context in red light. Next, 24 hr later, mice were placed into the same context for 3 min of habituation followed by three tone-shock pairings, consisting of a 30 s auditory tone (6kHz, 85dB) co-terminating with a foot shock (1 s, 0.5 mA) with an intertrial intervals (ITIs) of 90, 60 and 30 s. Memory retention testing was conducted 48 hrs later with no foot shocks co-terminating with tone presentation, and in a novel context consisting of the addition of a white floor and a curved white wall to change the shape of the chamber, an orange scent, a white noise generator, and white room lights. Additionally, 72 hrs following fear conditioning, some mice were placed into the fear conditioning context with no tones or foot shocks for 6 min. Some mice, 1 wk after fear conditioning, were placed into the fear conditioning context and underwent 3 min for habituation, followed by 30 presentations of the 30 s auditory tone with 30 s ITIs, but no foot shocks were administered to induce extinction. Freezing behavior was assessed during habituation, tone presentation, ITIs for cued and contextual tests, and extinction.

Contextual Fear Conditioning: Behavioral assays that were part of a battery were separated by one week. During fear conditioning, mice underwent 2 min of habituation prior to three 2 s 0.75 mA foot shocks. Foot shocks were separated by 60 s and 90 s inter-trial intervals (ITIs) and followed by a 30 s observation period. Contextual memory retention testing occurred 24 or 48 hrs later during a 5 min test in the fear conditioning context.

*Elevated Plus Maze (EPM)*

A standard EPM apparatus was used (Med Associates), which included two open white arms and two closed black arms with 30 cm high walls. The whole apparatus was elevated 60 cm from the floor, and behavior was recorded. Mice underwent a 5 min test conducted in low white light and with a white noise generator (San Diego Instruments) set at ~65-70 dB near the EPM apparatus. At the start of the session, mice were placed in the center of the apparatus facing an open arm and activity was monitored with CCTV cameras (Pansonic WV-BP334) feeding into a computer equipped with Ethovision XT (Noluds Information Technology) for data acquisition and analysis.

*Open Field (OF)*

Thigmotaxis, locomotion, and time spent in zones (center and Sides/Corners) were measured in an open field. Custom built open field boxes that were 17 in x 17 in with 12 in high plastic walls surrounded by opaque walls were used. Mice were placed in the center of the box and activity was monitored with CCTV cameras (Pansonic WV-BP334) feeding into a computer equipped with Ethovision XT (Noluds Information Technology) for data acquisition and analysis. Testing was conducted in low white or red light and with a white noise generator (San Diego Instruments) set at ~65-70 dB was near the OF box.

*Rotarod*

A five-lane accelerating rotarod (Med Associates) was used to assay balance and motor coordination over three trials separated by 30 min. For each trial, mice were placed on an 15 cm elevated rod, which accelerates from 4 to 40 rpm over 5 min or when a mouse fell off, whichever occurred first. The latency to fall off the rod was measured by photobeam breaks and averaged across all three trials. Testing was conducted in white light and with a white noise generator (San Diego Instruments) set at ~65-70 dB was below the apparatus.

*Spontaneous Alternation*

Spatial working memory was tested using a T maze with photobeams, and was comprised of a start box, entry arm and two choice arms (Med Associates). Matte white walls were placed outside of the maze to prevent observation of outside contextual cues. Testing was conducted in low white light and with a white noise generator (San Diego Instruments) set at ~65-70 dB was near the T maze. Mice were habituated to the room for 5 min before being placed in the start box with the door closed for no delay or 1 min. The door then opened, which allowed the mouse to freely explore for 5 min or when a mouse picked a choice arm, after which the door automatically closed. The mouse was removed, and the apparatus was cleaned while the mouse was placed back into the start box for no delay or 1 min. The mouse was then allowed free access again to pick a choice arm, using the same methods as before. Three days later, mice were tested again with no delay or 5 min in a counterbalanced manner, whichever was not used during the first test. Behavior was monitored with CCTV cameras (Pansonic WV-BP334) feeding into a computer equipped with Ethovision XT (Noluds Information Technology) for data acquisition and analysis.

**Pharmacokinetics**

Blood was collected from rats or mice at critical time points post infusion of Blebb or a derivative into lithium heparin coated tubes (cat. 07 6101, Ram Scientific), and stored on wet ice. Blood samples were later centrifuged for 3 min at 2655 g to separate plasma from red blood cells. Plasma was collected into a fresh tube and stored at -80 °C. In addition to blood samples, temporal lobe brain tissue was collected from each mouse. Each sample was flash frozen with 2-methylbutane and stored at -80 °C. Compound levels were quantified in brain and plasma by mass spectrometry using an ABSciex 5500 mass spectrometer using multiple reaction monitoring. Brain samples were homogenized in water and then immediately treated with 5‑times (v:v) acetonitrile to extract the compound and precipitate cellular protein. Plasma samples were directly treated with acetonitrile. Samples were filtered through a 0.45 µm filter plate prior to injection onto the LC‑MS/MS.

**Echocardiogram**

Animals were lightly anesthetized using 1.5-2.5% isoflurane in oxygen, delivered using a nose cone. An intravenous catheter was placed into the tail vein for dosing purposes. The chest area was clipped, and a depilatory cream applied to remove fur. Animals were positioned supine on the heated handling table of a Vevo 2100 high-frequency ultrasound unit (Visualsonics, Toronto, Canada). Electrode coupling gel was applied to the animal’s paws and the paws taped to the ECG sensors of the platform to allow simultaneous real-time collection of electrocardiograms. A rectal probe was placed to monitor body temperature and an overhead lamp used as supplemental heat to maintain body temperature during the imaging procedure. Warm ultrasound gel was applied to the skin overlying the heart. Baseline mid-papillary SAX (short axis) views of the left ventricle were collected in standard B-mode and M-mode using a 21 MHz transducer clipped to the mechanical arm of the imaging table. Animals were dosed with vehicle or test article according to the Treatment Schedule. Doses were delivered over approximately 30 sec at a dose volume of 2 mL/kg with approximately 10 min between doses. SAX echocardiogram images were collected at 0, 5 and 10 min after the initiation of each treatment. Analyses were performed post-acquisition using Visualsonics VevoLab software.

**Metabolite Identification**

MT-228 metabolites were evaluated in rat liver microsomes and in plasma after dosing rats IP with 10 mg/kg MT-228. Microsomal incubations were done using 1 mg/ml microsomal protein in 0.1M Tris-HCl buffer pH 7.4 with 1 µM MT-228 with and without 1 mM NADPH at 37 °C. Aliquots were transferred to acetonitrile/methanol (ACN/MeOH, 1:1, v:v) and internal standard (IS, diclofenac) containing tubes to terminate the reaction at time points 0, 2, 5, 10, 20, 30, and 60 minutes. Plasma samples were spiked with IS, followed by the addition of ACN/MeOH, 1:1, v:v. The samples were vortexed for 0.5 min before centrifugation at 15,000 x *g* for 10 minutes at 4 °C. The supernatant was directly analyzed using a Thermo Scientific Q Exactive hybrid quadrupole-Orbitrap mass spectrometer in positive ion Full Scan ddMS^2^ mode. The obtained MS raw data files were submitted to Compound Discoverer 3.3 software for metabolite identification.

**Crystallization of the SmMII·MT-228 complex and X-ray data processing**

Samples containing 10 mg/ml SmII MD were incubated on ice with 2 mM MgADP and 0.5 mM MT-228 for 30 min. Finally, 2 mM Na_3_VO_4_ were added to the mixture and incubated for 30 min. Crystallization by hanging drop vapor diffusion was immediately performed at 277 K by mixing 1 μl of protein solution with 1 μl of reservoir solution (10% PEG 3350; 50 mM Bicine, pH 8.2). Crystals grew spontaneously as branched plates approximately 3 weeks after. They were cryo-cooled in liquid nitrogen in a solution containing 11% PEG 3350; 50 mM Bicine pH 8.2; 10% DMSO; 0.5 mM MT-228; 30% glycerol. Despite our intensive efforts, NM2b fragments containing the Motor Domain (MD) were, in contrast, reluctant to co-crystallize with MT-228.

Several exploitable X-ray datasets of SmII·MT-228 were collected at the Proxima 2 beamline (Synchrotron Soleil, Gif-Sur-Yvette) and processed with Autoproc^3^. Initial structure factors were obtained by molecular replacement with Molrep^4^, showing clear positive density in Fo-Fc corresponding to MT-228, followed by several cycles of iterative edition with Coot^5^ and model refinement with Buster^6^. Resolution was automatically cut by Buster to 2.58 Å based on model-map cross-correlation. B-factors attributed to the compound by Buster are in the same range as of those of the surrounding residues (**Fig. S8**), suggesting full occupancy of the binding site. Crystallographic statistics are reported in **Table S5**. Sequences of the other myosins II were submitted to Swiss Model (*25*) for model building using the SmMII·MT-228 structure as template in User Template Mode.

Supporting Tables

**Table S1. NMIIA and SmMII selectivity**

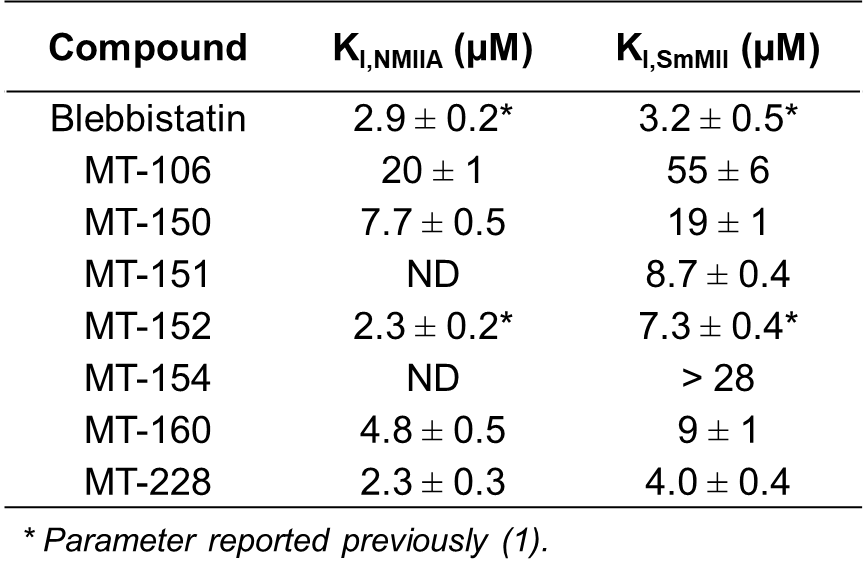

**Table S2. NMII Inhibitor ADMET Properties (CYP, hERG, P-gp, PPB)**

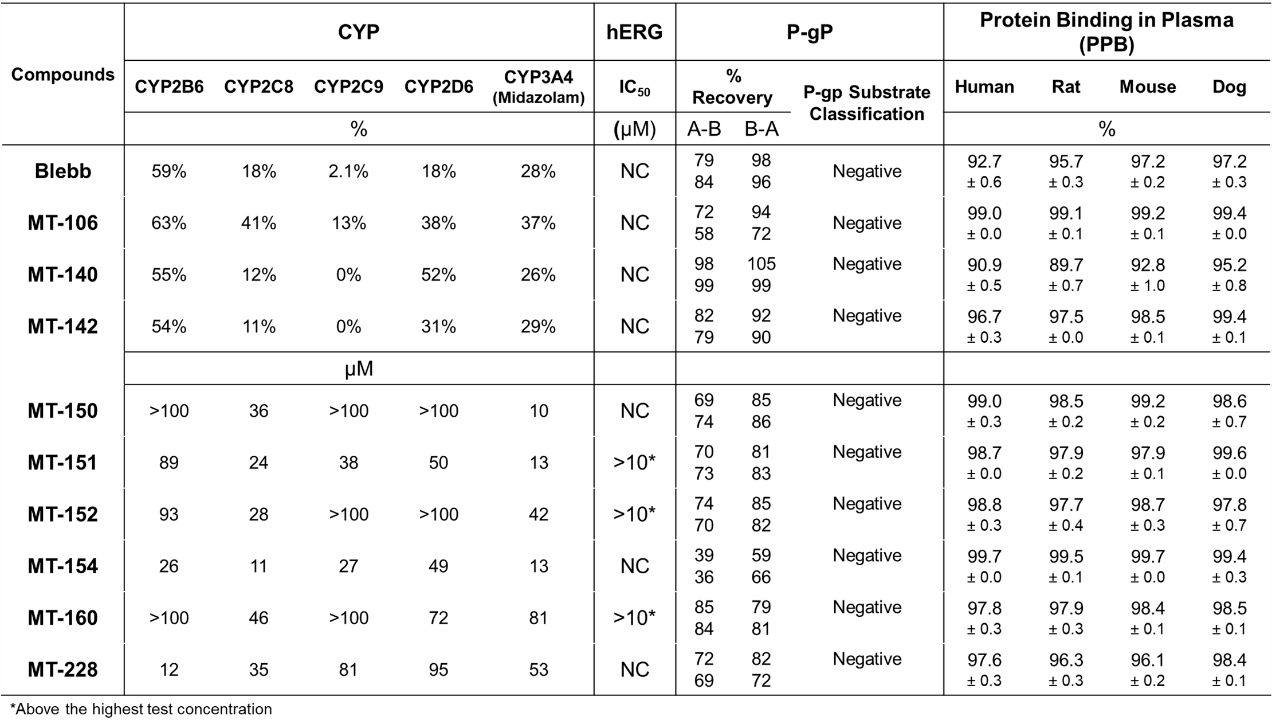

**Table S3. NMII Inhibitor Metabolic Stability**

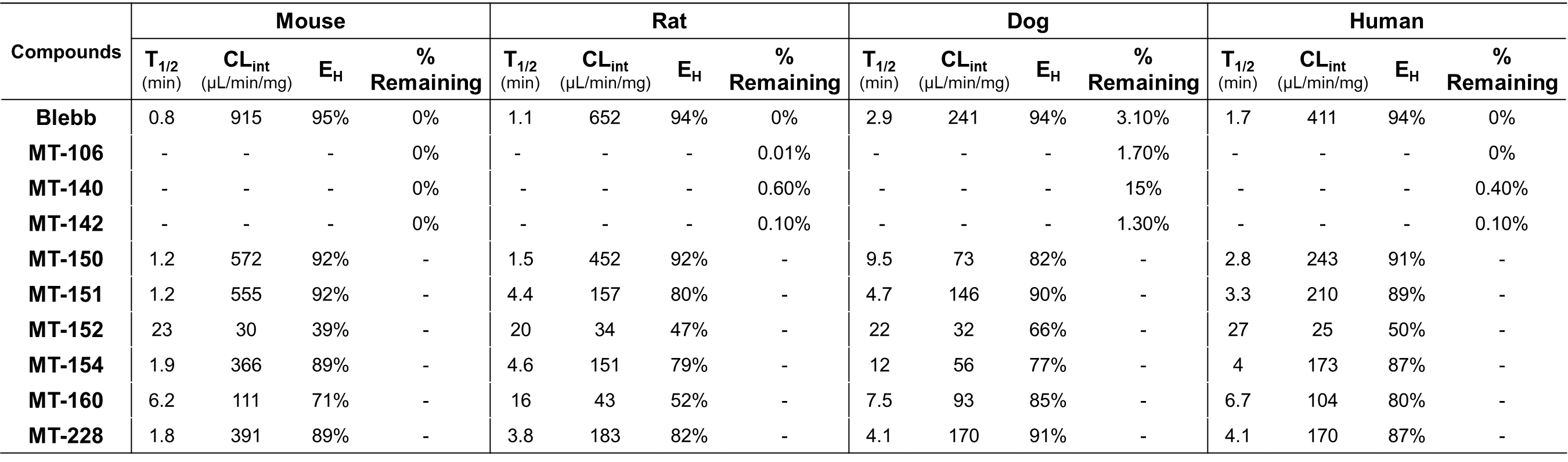

**Table S4. Comparison between the MT-228 binding site in SmMII and the Blebb binding site in Dicty Myo2.**

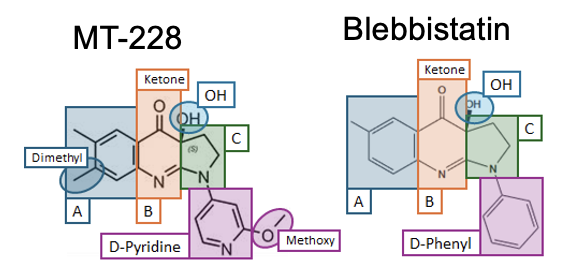

| **MT-228** | **SMM2** | **Closest Interaction** | **Blebbistatin** | **Dd Myo2 (1YV3)** | **Closest Interaction** |
| --- | --- | --- | --- | --- | --- |
| **(A)** | _Hw-h_L664 | 3.6 Å | (A) | _Hw-h_L641 | 3.6 Å |
| **(A)*** | _Hw-h_Q660 | 4.5 Å | (A) | _Hw-h_Q637 | 3.6 Å |
| **(A)** | _Hp-h_T479 | 3.7 Å (different anchor) | (A) | _Hp-h_T474 | 3.7 Å |
| **(A)** | _Hw-h_**Y657** | 3.3 Å (pi-stacking) | (A) | _Hw-h_**Y634** | 3.6 Å (pi-stacking) |
| **(A)** | _ɸLL-linker_Y264 | 3.6 Å (pi-stacking) | (A) | _ɸLL-linker_Y261 | 3.5 Å (pi-stacking) |
| **(A)** | _Hw-h_L661 | 4.5 Å | (A) | _Hw-h_L638 | 4.2 Å |
| **(A)** | _Sw2_I460 | 4.0 Å | (A) | _Sw2_I455 | 4.5 Å |
| **A-Dimethyl** | _Hw-h_Q660 | 4.0 Å | **N/A** | _Hw-h_Q637 |  |
| **A-Dimethyl** | _ɸLL-linker_Y264 | 4.4 Å | **N/A** | _ɸLL-linker_Y261 |  |
| **A-Dimethyl** | _Hw-h_**Y657** | 3.7 Å | **N/A** | _Hw-h_**Y634** |  |
| **A-Dimethyl** | _Hw-h_L661 | 3.7 Å | **N/A** | _Hw-h_L638 |  |
| **(B)** | _Hw-h_**Y657** | 4.3 Å | (B) | _Hw-h_**Y634** | 3.8 Å |
| **(B)** | _ɸLL-linker_Y264 | 4.8 Å | (B) | _ɸLL-linker_Y261 | 4.4 Å |
| **(B)** | _ɸLL-linker_L265 | 4.0 Å | (B) | _ɸLL-linker_L262 | 3.7 Å |
| **B-Ketone** | _Sw2_I460 | 3.8 Å | B-Ketone | _Sw2_I455 | 4.0 Å |
| **B-Ketone** | _Transd_G243 mc | 3.7 Å (electrostatic) | B-Ketone | _Transd_G240 mc | 3.4 Å (electrostatic) |
| **B-Ketone** | _Sw2_A461 mc | 2.9 Å (electrostatic) | B-Ketone | _Sw2_S456 mc | 3.2 Å (electrostatic) |
| **OH group** | _ɸLL-linker_L265 mc O | 2.6 Å (H-bond) | OH group | _ɸLL-linker_L262 mc O | 2.5 Å (H-bond) |
| **OH group** | _ɸLL-linker_G243 mc N | 2.9 Å (H-bond) | OH group | _ɸLL-linker_G240 mc N | 2.8 Å (H-bond) |
| **OH group** | _ɸLL-linker_Y264 | 3.9 Å | OH group | _ɸLL-linker_Y261 | 3.4 Å |
| **(C)** | _Sw2_A461 | 3.6 Å | (C) | _Sw2_S456 | 3.1 Å (O) 3.6 Å (C) |
| **(C)** | _Hp-h_I476 | 3.7 Å | (C) | _Hp-h_I471 | 3.8 Å |
| **(C)** | _ɸLL-linker_L265 | 4.2 Å | (C) | _ɸLL-linker_L262 | 4.0 Å |
| **(C)** | _Transd_F242 mc | 3.7 Å | (C) | _Transd_F239 mc | 3.9 Å |
| **(C)** | _Sw1_R241 mc | 3.7 Å | (C) | _Sw1_R238 mc | 3.7 Å |
| **D-Pyridine** | _ɸLL-linker_L265 | 3.8 Å | **D-Phenyl** | _ɸLL-linker_L262 | 3.6 Å |
| **D-Pyridine** | _Hp-h_F471 |  | **D-Phenyl** | _Hp-h_F466 | 4.2 Å |
| **D-Pyridine** | _ɸLL-linker_L265-L266 mc | 3.4 Å | **D-Phenyl** | _ɸLL-linker_L262-L263 mc | 3.7 Å |
| **D-Pyridine** | _ɸLL-linker_E267 mc | 3.6 Å | **D-Phenyl** | _ɸLL-linker_E264 mc | 4.0 Å |
| **D-Pyridine** | _Hp-h_E472 | 3.4 Å | **D-Phenyl** | _Hp-h_E467 | 3.5 Å |
| **D-Pyridine** | _Hp-h_C475 | 4.0 Å | **D-Phenyl** | _Hp-h_C470 | 3.6 Å |
| **D-Pyridine** | _Hw-h_V653 | 4.9 Å | **D-Phenyl** | _Hw-h_V630 | 4.1 Å |
| **D-Pyridine** | _Hw-h_**Y657** | 5.2 Å | **D-Phenyl** | _Hw-h_**Y634** | 4.3 Å |
| **D-Methoxy** | _Hw-h_**Y657** | 4.8 Å | **N/A** | _Hw-h_**Y634** |  |
| **D-Methoxy** | _Hw-h_V653 | 3.7 Å | **N/A** | _Hw-h_V630 |  |
| **D-Methoxy** | _Hp-h_F471 | 3.5 Å | **N/A** | _Hp-h_F466 |  |
| **D-Methoxy** | _Hp-h_C475 | 3.4 Å | **N/A** | _Hp-h_C470 |  |
| **D-Methoxy** | _ɸLL-linker_L265 | 3.5 Å | **N/A** | _ɸLL-linker_L262 |  |

* Grey cells indicate interactions are noticeably different (> 0.5 Å difference in minimal distance found, or interaction only exists for one structure); orange cells indicate no interaction.

**Table S5. Data collection and refinement statistics**

|  | **PDB 9FU2 (SmMII・MT-228)** |
| --- | --- |
| Wavelength (Å) | 0.9801 |
| Resolution range (Å) | 71.97  - 2.581 (2.66  - 2.58*) |
| Space group | P 3_1_ |
| Unit cell (*a,b,c; α,ß,γ*) | 83.1 Å 83.1 Å 132.3 Å 90° 90° 120° |
| Total reflections | 48765 (4136) |
| Unique reflections | 30543 (2620) |
| Multiplicity | 1.6 (1.6) |
| Completeness (%) | 99.96 (100.00) |
| Mean I/sigma(I) | 5.31 (1.29) |
| Wilson B-factor (Å^2^) | 62.8 |
| R-merge | 0.05975 (0.7954) |
| R-meas | 0.08252 (1.109) |
| R-pim | 0.05661 (0.7704) |
| CC1/2 | 0.997 (0.369) |
| CC* | 0.999 (0.734) |
| Reflections used in refinement | 32138 (2682) |
| Reflections used for R-free | 1621 (131) |
| R-work | 0.2292 |
| R-free | 0.2497 |
| Number of non-hydrogen atoms | 5855 |
| macromolecules | 5690 |
| ligands | 95 |
| solvent | 70 |
| Protein residues | 713 |
| RMS(bonds) | 0.091 |
| RMS(angles) | 1.12 |
| Ramachandran favored (%) | 96.29 |
| Ramachandran allowed (%) | 3.71 |
| Ramachandran outliers (%) | 0.00 |
| Rotamer outliers (%) | 1.80 |
| Clashscore | 1.56 |
| Average B-factor (Å^2^) | 79.59 |
| macromolecules | 79.87 |
| ligands | 74.76 |
| solvent | 63.42 |

* Statistics for the highest-resolution shell are shown in parentheses.

Supporting Figures

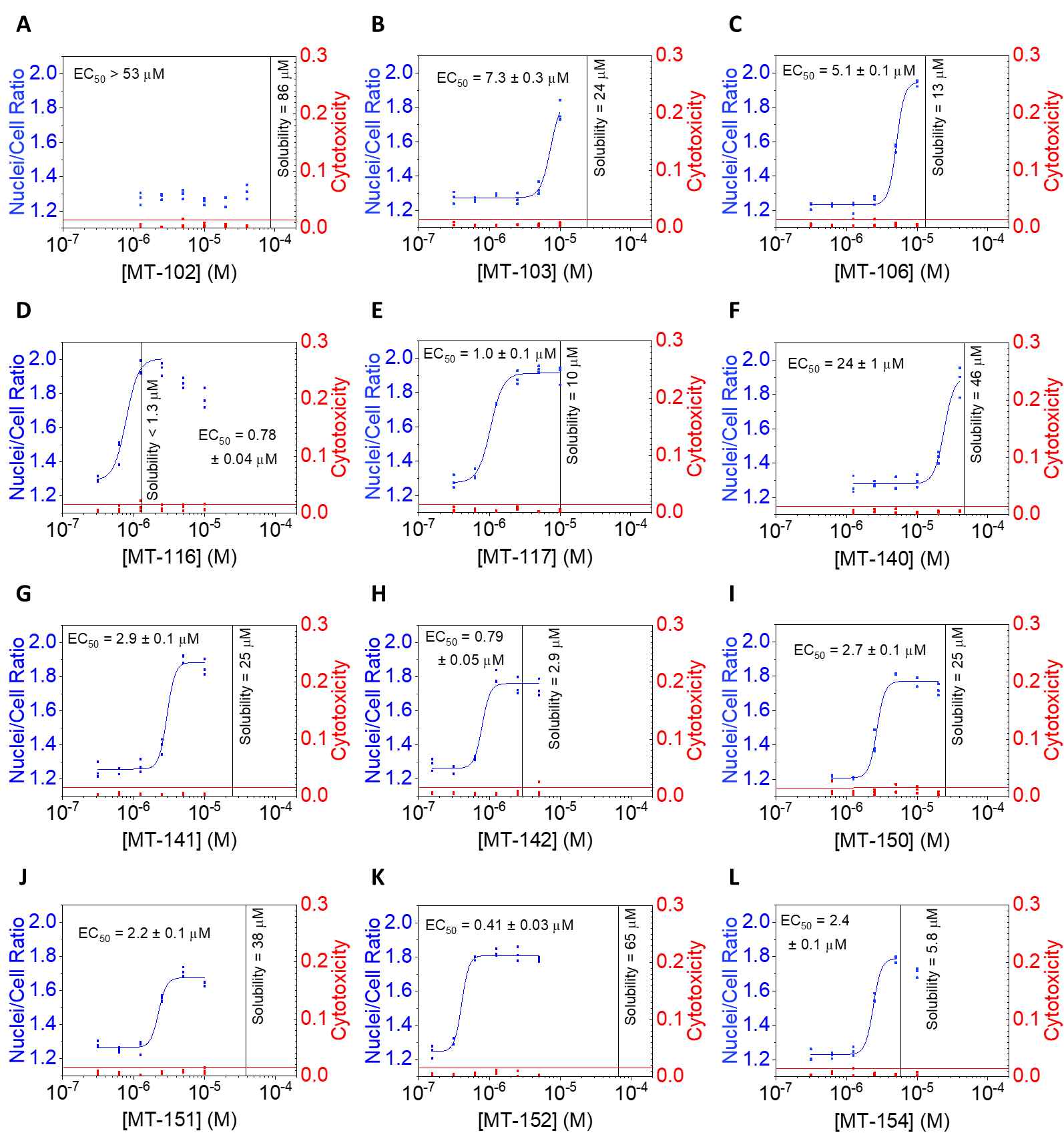

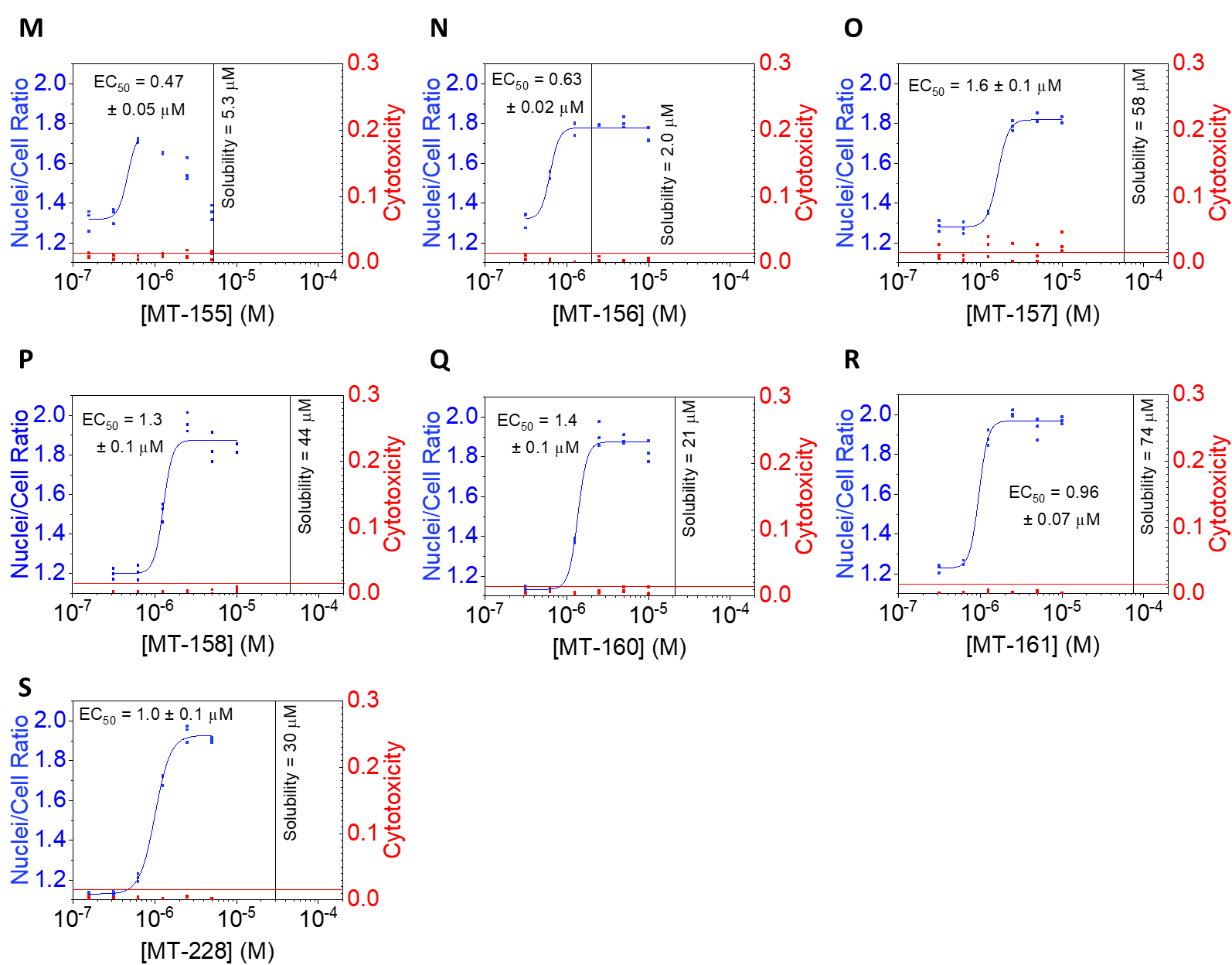

**Figure S1. NMIIB inhibition in the cytokinesis assay.** The inhibitory effect of Blebb derivatives on NMIIB was evaluated in a cytokinesis assay as described previously (*26*). The nuclei-to-cell ratio (left axes, blue) and cytotoxicity (right axes, red) were determined for each well in the assay plate. Cytotoxicity was calculated as the dead nuclei to total nuclei ratio. Cytotoxicity is considered significant if above the empirical threshold of 0.015 (red horizontal lines) (*26*). Vertical black lines represent the kinetic aqueous solubility determined for each compound. It must be noted that solubility in culture medium may be different due to the presence of proteins, salts, and other components. Since compound precipitates may influence dose-response curves (*26*), data analysis was generally avoided in potentially affected regions (see D, L, and M). EC_50_ values were determined by fitting the dose-response data to the Hill equation (*26*). EC_50_ values are shown with the standard error of fitting.

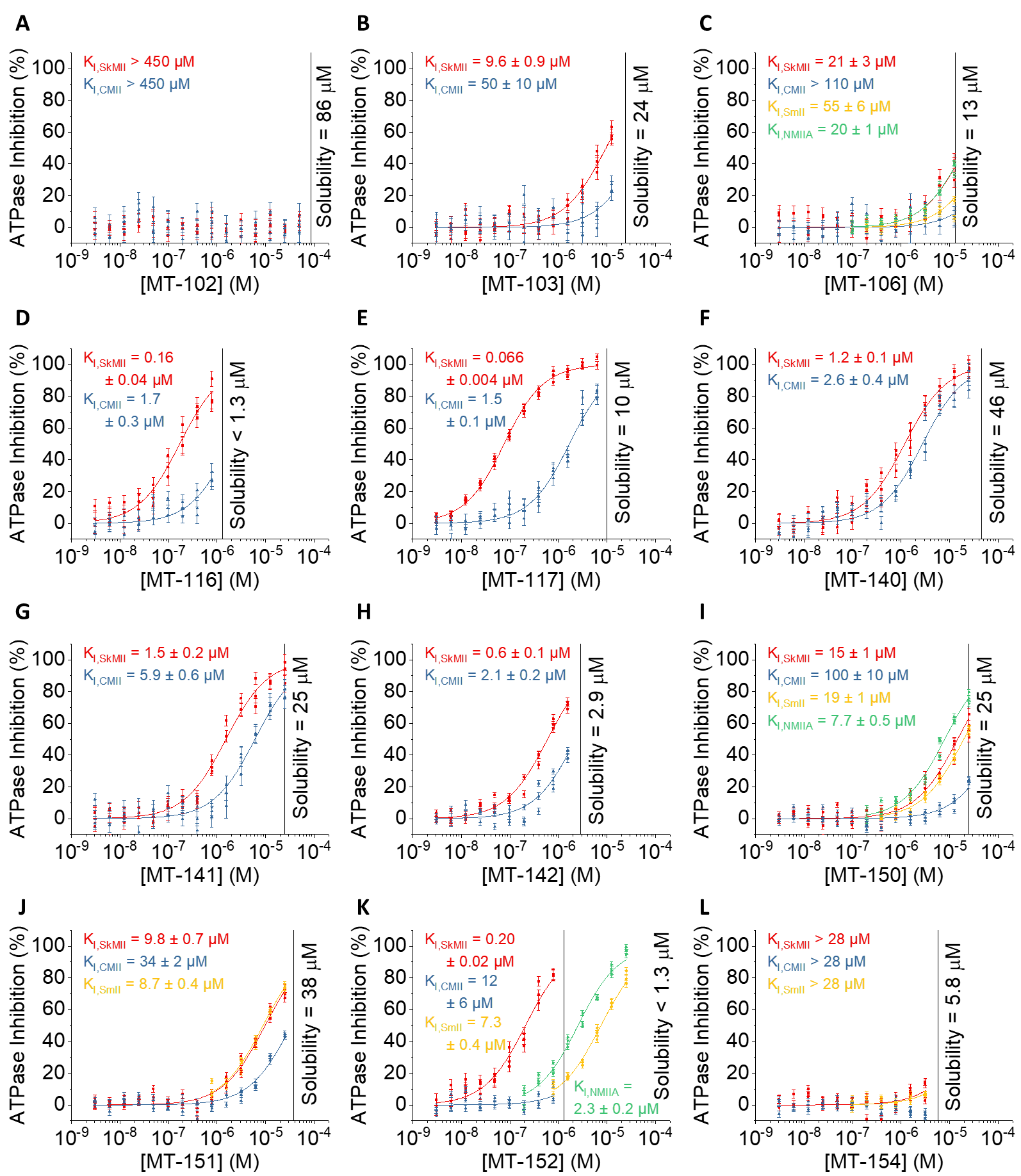

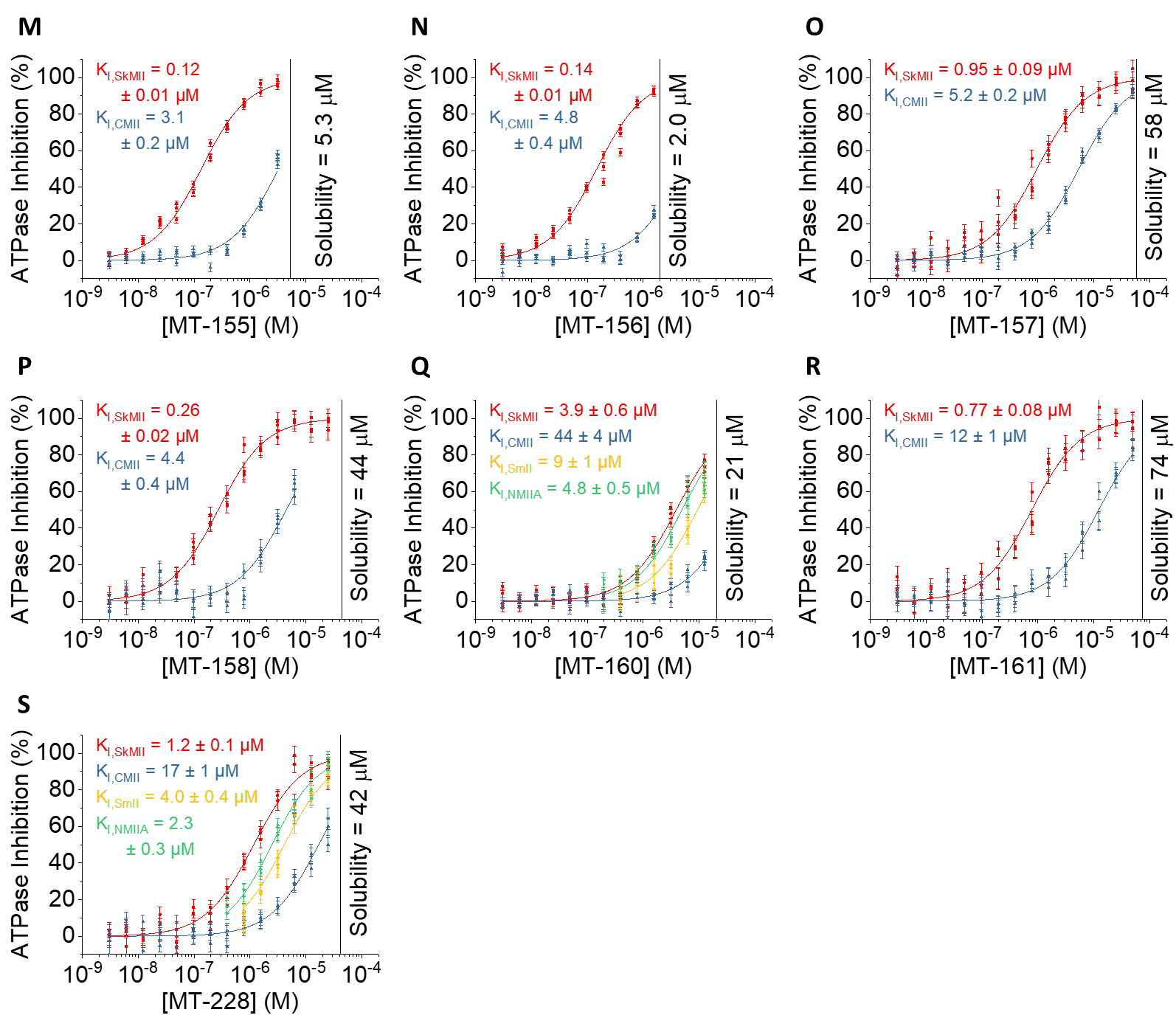

**Figure S2. Inhibitory effects of Blebb derivatives on the actin-activated steady-state ATPase activity of SkMII (red), CMII (blue), SmMII (yellow), and NMIIA (green).** ATPase assays were performed as described previously(*2*). Inhibitory constants (K_I_) were determined by fitting the dose response data to a quadratic equation corresponding to a simple 1:1 equilibrium binding model(*2*). No data analysis was performed above the kinetic aqueous solubilities of the compounds (vertical black lines). The only exception was MT-152 where the more accurate determination of K_I,SmII_ and K_I,NMIIA_ required some data above the solubility. Although no sign of compound precipitation was observed in these cases, it is important to note that these inhibitory constants could be affected by compound precipitation-related artifacts. Errors represent the standard error of fitting.

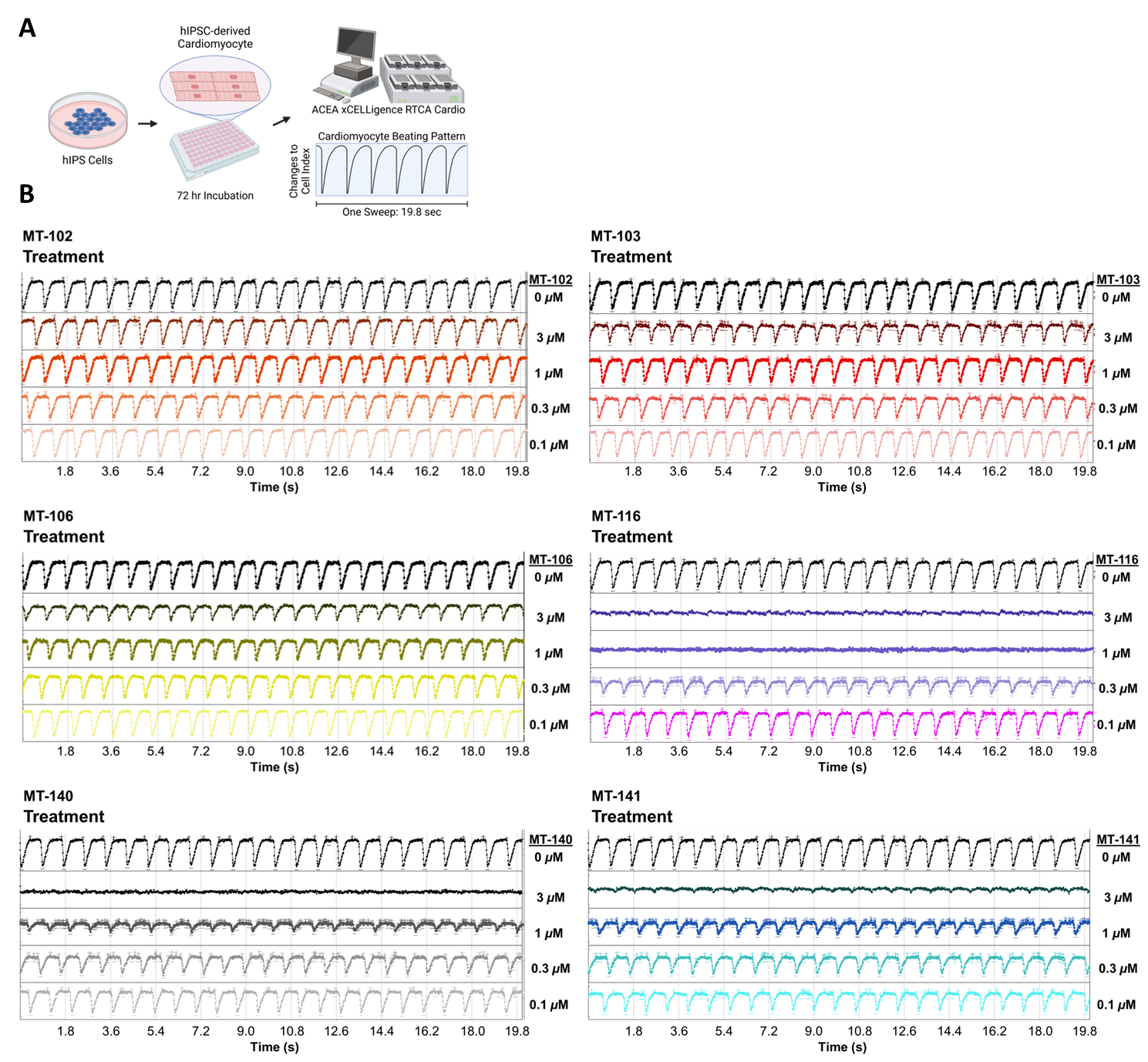

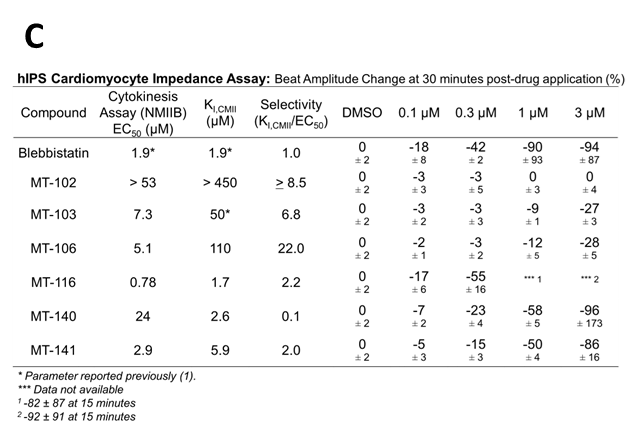

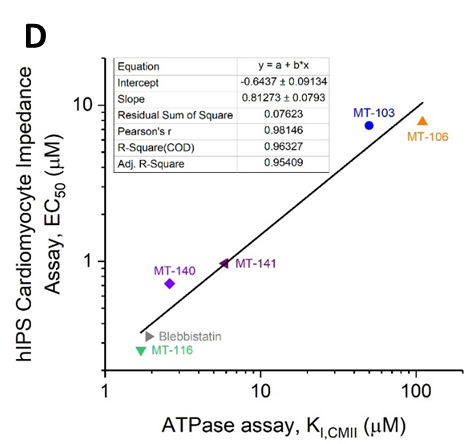

**Figure S3. CMII potency of derivatives corresponds to contractility effects in hIPS cardiomyocytes.** A) The experimental timeline for incubation of hIPS cardiomyocytes with different compounds and the measurement of contractility. B) Traces of contractility before (“Baseline”) and after 30 min of incubation with a derivative (“Treatment”). C) Displays the selectivity and percentage change in contractility for each derivative at each dose. D) Inhibitory constants determined in the ATPase assay predict the potency (EC_50_) of blebb derivatives in the hIPS Cardiomyocyte Impedance Assay (Pearson correlation coefficient = 0.98).

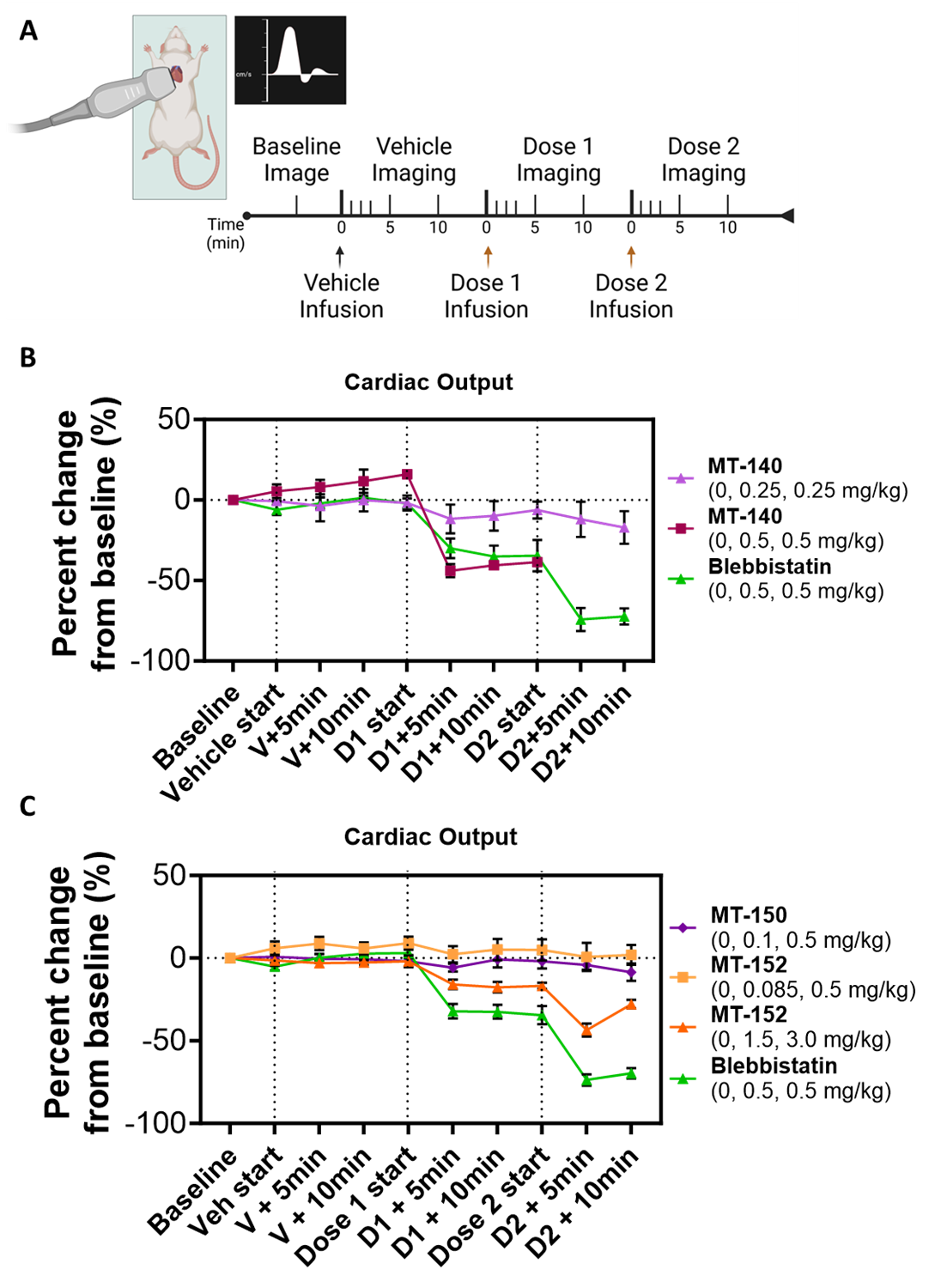

**Figure S4. Derivatives with reduced CMII potency show improved cardiac safety.** A) Echocardiogram (echo) experimental timeline indicating timing of treatments and imaging. B) **MT-140** showed improved cardiac safety compared to Blebb at lower doses, but still produced significant effects at higher doses. C) Compared to the same dose as Blebb, **MT-150** and **MT-152** had no significant cardiac effects. However, increasing the dose of **MT-152** to 1.5 and 3.0 mg/kg resulted in cardiac effects. Error bars represent SEM, * *P* < 0.05, ** *P* < 0.01.

```
SMII  |P35749|MYH11_HUMAN MAQKGQLSDD EKFLFVDKN- FINSP----- -VAQADWAAK RLVWVPSEKQ GFEAASIKEE KGDEVVVELV ENGKKVTVGK DDIQKMNPPK FSKVEDMAEL    93
NMIIA |P35579|MYH9_HUMAN  ----MAQQAA DKYLYVDKN- FINNP----- -LAQADWAAK KLVWVPSDKS GFEPASLKEE VGEEAIVELV ENGKKVKVNK DDIQKMNPPK FSKVEDMAEL    89
NMIIB |P35580|MYH10_HUMAN MAQRTGLEDP ERYLFVDRAV IYNP------ -ATQADWTAK KLVWIPSERH GFEAASIKEE RGDEVMVELA ENGKKAMVNK DDIQKMNPPK FSKVEDMAEL    93
CMII  |P12883|MYH7_HUMAN  ------MGDS EMAVFGAAAP YLRKSEKERL EAQTRPFDLK KDVFVPDDKQ EFVKAKIVSR EGGKVTAET- EYGKTVTVKE DQVMQQNPPK FDKIEDMAML    93
```

SkMII |Q9UKX3|MYH13_HUMAN -----MSSDA EMAIFGEAAP YLRKPEKERI EAQNRPFDSK KACFVADNKE MYVKGMIQTR ENDKVIVKTL DD-RMLTLNN DQVFPMNPPK FDKIEDMAMM	94

Transd

Transd

```
SMII  |P35749|MYH11_HUMAN TCLNEASVLH NLRERYFSGL IYTYSGLFCV VVNPYKHLPI YSEKIVDMYK GKKRHEMPPH IYAIADTAYR SMLQDREDQS ILCTGESGAG KTENTKKVIQ   193
NMIIA |P35579|MYH9_HUMAN  TCLNEASVLH NLKERYYSGL IYTYSGLFCV VINPYKNLPI YSEEIVEMYK GKKRHEMPPH IYAITDTAYR SMMQDREDQS ILCTGESGAG KTENTKKVIQ   189
NMIIB |P35580|MYH10_HUMAN TCLNEASVLH NLKDRYYSGL IYTYSGLFCV VINPYKNLPI YSENIIEMYR GKKRHEMPPH IYAISESAYR CMLQDREDQS ILCTGESGAG KTENTKKVIQ   193
CMII  |P12883|MYH7_HUMAN  TFLHEPAVLY NLKDRYGSWM IYTYSGLFCV TVNPYKWLPV YTPEVVAAYR GKKRSEAPPH IFSISDNAYQ YMLTDRENQS ILITGESGAG KTVNTKRVIQ   193
```

SkMII |Q9UKX3|MYH13_HUMAN THLHEPAVLY NLKERYAAWM IYTYSGLFCV TVNPYKWLPV YKPEVVAAYR GKKRQEAPPH IFSISDNAYQ FMLTDRDNQS ILITGESGAG KTVNTKRVIQ	194

Sw1

Transd

ΦLL-linker

```
SMII  |P35749|MYH11_HUMAN YLAVVA-SSH KGKKDTS--- ITGELEKQLL QANPILEAFG NAKTVKNDNS SRFGKFIRIN FDVTGYIVGA NIETYLLEKS RAIRQARDER TFHIFYYMIA   289
NMIIA |P35579|MYH9_HUMAN  YLAYVA-SSH KSKKDQ---- --GELERQLL QANPILEAFG NAKTVKNDNS SRFGKFIRIN FDVNGYIVGA NIETYLLEKS RAIRQAKEER TFHIFYYLLS   282
NMIIB |P35580|MYH10_HUMAN YLAHVA-SSH KGRKDHN--- IPGELERQLL QANPILESFG NAKTVKNDNS SRFGKFIRIN FDVTGYIVGA NIETYLLEKS RAVRQAKDER TFHIFYQLLS   289
CMII  |P12883|MYH7_HUMAN  YFAVIAAIGD RSKKDQSPGK --GTLEDQII QANPALEAFG NAKTVRNDNS SRFGKFIRIH FGATGKLASA DIETYLLEKS RVIFQLKAER DYHIFYQILS   291
SkMII |Q9UKX3|MYH13_HUMAN YFATIAVTGD K-KKETQPGK MQGTLEDQII QANPLLEAFG NAKTVRNDNS SRFGKFIRIH FGATGKLASA DIETYLLEKS RVTFQLSSER SYHIFYQIMS   293

SMII  |P35749|MYH11_HUMAN GAKEKMRSDL LLEGFN--NY TFLSNGFVPI PAAQDDEMFQ ETVEAMAIMG FSEEEQLSIL KVVSSVLQLG NIVFKKERNT DQASMPDNT- AAQKVCHLMG   386
NMIIA |P35579|MYH9_HUMAN  GAGEHLKTDL LLEPYN--KY RFLSNGHVTI PGQQDKDMFQ ETMEAMRIMG IPEEEQMGLL RVISGVLQLG NIVFKKERNT DQASMPDNT- AAQKVSHLLG   379
NMIIB |P35580|MYH10_HUMAN GAGEHLKSDL LLEGFN--NY RFLSNGYIPI PGQQDKDNFQ ETMEAMHIMG FSHEEILSML KVVSSVLQFG NISFKKERNT DQASMPENT- VAQKLCHLLG   386
CMII  |P12883|MYH7_HUMAN  NKKPEL-LDM LLITNNPYDY AFISQGETTV ASIDDAEELM ATDNAFDVLG FTSEEKNSMY KLTGAIMHFG NMKFKLKQRE EQAE-PDGTE EADKSAYLMG   389
```

SkMII |Q9UKX3|MYH13_HUMAN NKKPEL-IDL LLISTNPFDF PFVSQGEVTV ASIDDSEELL ATDNAIDILG FSSEEKVGIY KLTGAVMHYG NMKFKQKQRE EQAE-PDGTE VADKAGYLMG	391

Transd

Sw2

Hp-h

```
SMII  |P35749|MYH11_HUMAN INVTDFTRSI LTPRIKVGRD VVQKAQTKEQ ADFAVEALAK ATYERLFRWI LTRVNKALDK THRQGASFLG ILDIAGFEIF EVNSFEQLCI NYTNEKLQQL   486
NMIIA |P35579|MYH9_HUMAN  INVTDFTRGI LTPRIKVGRD YVQKAQTKEQ ADFAIEALAK ATYERMFRWL VLRINKALDK TKRQGASFIG ILDIAGFEIF DLNSFEQLCI NYTNEKLQQL   479
NMIIB |P35580|MYH10_HUMAN MNVMEFTRAI LTPRIKVGRD YVQKAQTKEQ ADFAVEALAK ATYERLFRWL VHRINKALDR TKRQGASFIG ILDIAGFEIF ELNSFEQLCI NYTNEKLQQL   486
CMII  |P12883|MYH7_HUMAN  LNSADLLKGL CHPRVKVGNE YVTKGQNVQQ VIYATGALAK AVYERMFNWM VTRINATLE- TKQPRQYFIG VLDIAGFEIF DFNSFEQLCI NFTNEKLQQF   488
```

SkMII |Q9UKX3|MYH13_HUMAN LNSAEMLKGL CCPRVKVGNE YVTKGQNVQQ VTNSVGALAK AVYEKMFLWM VTRINQQLD- TKQPRQYFIG VLDIAGFEIF DFNSLEQLCI NFTNEKLQQF	490

Hp-h

```
SMII  |P35749|MYH11_HUMAN FNHTMFILEQ EEYQREGIEW NFIDFGLDLQ PCIELIERPN NPPGVLALLD EECWFPKATD KSFVEKLCTE Q-GSHPKFQK PKQLKDKTE- -FSIIHYAGK   583
NMIIA |P35579|MYH9_HUMAN  FNHTMFILEQ EEYQREGIEW NFIDFGLDLQ PCIDLIEKPA GPPGILALLD EECWFPKATD KSFVEKVMQE Q-GTHPKFQK PKQLKDKAD- -FCIIHYAGK   576
NMIIB |P35580|MYH10_HUMAN FNHTMFILEQ EEYQREGIEW NFIDFGLDLQ PCIDLIERPA NPPGVLALLD EECWFPKATD KTFVEKLVQE Q-GSHSKFQK PRQLKDKAD- -FCIIHYAGK   583
CMII  |P12883|MYH7_HUMAN  FNHHMFVLEQ EEYKKEGIEW TFIDFGMDLQ ACIDLIEKPM ---GIMSILE EECMFPKATD MTFKAKLFDN HLGKSANFQK PRNIKGKPEA HFSLIHYAGI   585
```

SkMII |Q9UKX3|MYH13_HUMAN FNHHMFVLEQ EEYKKEGIEW EFIDFGMDLA ACIELIEKPM ---GIFSILE EECMFPKATD TSFKNKLYDQ HLGKSNNFQK PKPAKGKAEA HFSLVHYAGT	587

Transd

Hw-h

```
SMII  |P35749|MYH11_HUMAN VDYNASAWLT KNMDPLNDNV TSLLNASSDK FVADLWKDVD RIVGLDQMAK MTESSLPSAS KTKKG-MFRT VGQLYKEQLG KLMTTLRNTT PNFVRCIIPN   682
NMIIA |P35579|MYH9_HUMAN  VDYKADEWLM KNMDPLNDNI ATLLHQSSDK FVSELWKDVD RIIGLDQVAG MSETALPGAF KTRKG-MFRT VGQLYKEQLA KLMATLRNTN PNFVRCIIPN   675
NMIIB |P35580|MYH10_HUMAN VDYKADEWLM KNMDPLNDNV ATLLHQSSDR FVAELWKDVD RIVGLDQVTG MTETAFGSAY KTKKG-MFRT VGQLYKESLT KLMATLRNTN PNFVRCIIPN   682
CMII  |P12883|MYH7_HUMAN  VDYNIIGWLQ KNKDPLNETV VGLYQKSSLK LLSTLFAN-- -YAGADAPIE ------KGKG KAKKGSSFQT VSALHRENLN KLMTNLRSTH PHFVRCIIPN   676
SkMII |Q9UKX3|MYH13_HUMAN VDYNIAGWLD KNKDPLNETV VGLYQKSSLK LLSFLFSN-- -YAGAET--- -GDSGGSKKG GKKKGSSFQT VSAVFRENLN KLMTNLRSTH PHFVRCLIPN   680

SMII  |P35749|MYH11_HUMAN HEKRSGKLDA FLVLEQLRCN GVLEGIRICR QGFPNRIVFQ EFRQRYEILA ANAIPKG-FM DGKQACILMI KALELDPNLY RIGQSKIFFR TGVLAHLEEE   781
NMIIA |P35579|MYH9_HUMAN  HEKKAGKLDP HLVLDQLRCN GVLEGIRICR QGFPNRVVFQ EFRQRYEILT PNSIPKG-FM DGKQACVLMI KALELDSNLY RIGQSKVFFR AGVLAHLEEE   774
NMIIB |P35580|MYH10_HUMAN HEKRAGKLDP HLVLDQLRCN GVLEGIRICR QGFPNRIVFQ EFRQRYEILT PNAIPKG-FM DGKQACERMI RALELDPNLY RIGQSKIFFR AGVLAHLEEE   781
CMII  |P12883|MYH7_HUMAN  ETKSPGVMDN PLVMHQLRCN GVLEGIRICR KGFPNRILYG DFRQRYRILN PAAIPEGQFI DSRKGAEKLL SSLDIDHNQY KFGHTKVFFK AGLLGLLEEM   776
SkMII |Q9UKX3|MYH13_HUMAN ETKTPGVMDH YLVMHQLRCN GVLEGIRICR KGFPSRILYA DFKQRYRILN ASAIPEGQFI DSKNASEKLL NSIDVDREQF RFGNTKVFFK AGLLGLLEEM   780

SMII  |P35749|MYH11_HUMAN RDLKITDVIM AFQAMCRGYL ARKAFAKRQQ QLTAMKVIQR NCAAYLKLRN WQWWRLFTKV KPLLQVTRQE EEMQAKEDEL QKTKERQQKA ENELKELEQK   881
NMIIA |P35579|MYH9_HUMAN  RDLKITDVII GFQACCRGYL ARKAFAKRQQ QLTAMKVLQR NCAAYLKLRN WQWWRLFTKV KPLLQVSRQE EEMMAKEEEL VKVREKQLAA ENRLTEMETL   874
NMIIB |P35580|MYH10_HUMAN RDLKITDIII FFQAVCRGYL ARKAFAKKQQ QLSALKVLQR NCAAYLKLRH WQWWRVFTKV KPLLQVTRQE EELQAKDEEL LKVKEKQTKV EGELEEMERK   881
CMII  |P12883|MYH7_HUMAN  RDERLSRIIT RIQAQSRGVL ARMEYKKLLE RRDSLLVIQW NIRAFMGVKN WPWMKLYFKI KPLLKSAERE KEMASMKEEF TRLKEALEKS EARRKELEEK   876
SkMII |Q9UKX3|MYH13_HUMAN RDEKLVTLMT STQAVCRGYL MRVEFKKMME RRDSIFCIQY NIRSFMNVKH WPWMNLFFKI KPLLKSAEAE KEMATMKEDF ERTKEELARS EARRKELEEK   880

SMII  |P35749|MYH11_HUMAN HSQLTEEKNL LQEQLQAETE LYAEAEEMRV RLAAKKQELE EILHEMEARL EEEEDRGQQL QAERKKMAQQ MLDLEEQLEE EEAARQKLQL EKVTAEAKIK   981
NMIIA |P35579|MYH9_HUMAN  QSQLMAEKLQ LQEQLQAETE LCAEAEELRA RLTAKKQELE EICHDLEARV EEEEERCQHL QAEKKKMQQN IQELEEQLEE EESARQKLQL EKVTTEAKLK   974
NMIIB |P35580|MYH10_HUMAN HQQLLEEKNI LAEQLQAETE LFAEAEEMRA RLAAKKQELE EILHDLESRV EEEEERNQIL QNEKKKMQAH IQDLEEQLDE EEGARQKLQL EKVTAEAKIK   981
CMII  |P12883|MYH7_HUMAN  MVSLLQEKND LQLQVQAEQD NLADAEERCD QLIKNKIQLE AKVKEMNERL EDEEEMNAEL TAKKRKLEDE CSELKRDIDD LELTLAKVEK EKHATENKVK   976
SkMII |Q9UKX3|MYH13_HUMAN MVSLLQEKND LQLQVQSETE NLMDAEERCE GLIKSKILLE AKVKELTERL EEEEEMNSEL VAKKRNLEDK CSSLKRDIDD LELTLTKVEK EKHATENKVK   980

SMII  |P35749|MYH11_HUMAN KLEDEILVMD DQNNKLSKER KLLEERISDL TTNLAEEEEK AKNLTKLKNK HESMISELEV RLKKEEKSRQ ELEKLKRKLE GDASDFHEQI ADLQAQIAEL  1081
NMIIA |P35579|MYH9_HUMAN  KLEEEQIILE DQNCKLAKEK KLLEDRIAEF TTNLTEEEEK SKSLAKLKNK HEAMITDLEE RLRREEKQRQ ELEKTRRKLE GDSTDLSDQI AELQAQIAEL  1074
NMIIB |P35580|MYH10_HUMAN KMEEEILLLE DQNSKFIKEK KLMEDRIAEC SSQLAEEEEK AKNLAKIRNK QEVMISDLEE RLKKEEKTRQ ELEKAKRKLD GETTDLQDQI AELQAQIDEL  1081
CMII  |P12883|MYH7_HUMAN  NLTEEMAGLD EIIAKLTKEK KALQEAHQQA LDDLQAEEDK VNTLTKAKVK LEQQVDDLEG SLEQEKKVRM DLERAKRKLE GDLKLTQESI MDLENDKQQL  1076
SkMII |Q9UKX3|MYH13_HUMAN NLSEEMTALE ENISKLTKEK KSLQEAHQQT LDDLQVEEDK VNGLIKINAK LEQQTDDLEG SLEQEKKLRA DLERAKRKLE GDLKMSQESI MDLENDKQQI  1080

SMII  |P35749|MYH11_HUMAN KMQLAKKEEE LQAALARLDD EIAQKNNALK KIRELEGHIS DLQEDLDSER AARNKAEKQK RDLGEELEAL KTELEDTLDS TATQQELRAK REQEVTVLKK  1181
NMIIA |P35579|MYH9_HUMAN  KMQLAKKEEE LQAALARVEE EAAQKNMALK KIRELESQIS ELQEDLESER ASRNKAEKQK RDLGEELEAL KTELEDTLDS TAAQQELRSK REQEVNILKK  1174
NMIIB |P35580|MYH10_HUMAN KLQLAKKEEE LQGALARGDD ETLHKNNALK VVRELQAQIA ELQEDFESEK ASRNKAEKQK RDLSEELEAL KTELEDTLDT TAAQQELRTK REQEVAELKK  1181
CMII  |P12883|MYH7_HUMAN  DERLKKKDFE LNALNARIED EQALGSQLQK KLKELQARIE ELEEELEAER TARAKVEKLR SDLSRELEEI SERLEEAGGA TSVQIEMNKK REAEFQKMRR  1176
SkMII |Q9UKX3|MYH13_HUMAN EEKLKKKEFE LSQLQAKIDD EQVHSLQFQK KIKELQARIE ELEEEIEAEH TLRAKIEKQR SDLARELEEI SERLEEASGA TSAQIEMNKK REAEFQKMRR  1180

SMII  |P35749|MYH11_HUMAN ALDEETRSHE AQVQEMRQKH AQAVEELTEQ LEQFKRAKAN LDKNKQTLEK ENADLAGELR VLGQAKQEVE HKKKKLEAQV QELQSKCSDG ERARAELNDK  1281
NMIIA |P35579|MYH9_HUMAN  TLEEEAKTHE AQIQEMRQKH SQAVEELAEQ LEQTKRVKAN LEKAKQTLEN ERGELANEVK VLLQGKGDSE HKRKKVEAQL QELQVKFNEG ERVRTELADK  1274
NMIIB |P35580|MYH10_HUMAN ALEEETKNHE AQIQDMRQRH ATALEELSEQ LEQAKRFKAN LEKNKQGLET DNKELACEVK VLQQVKAESE HKRKKLDAQV QELHAKVSEG DRLRVELAEK  1281
CMII  |P12883|MYH7_HUMAN  DLEEATLQHE ATAAALRKKH ADSVAELGEQ IDNLQRVKQK LEKEKSEFKL ELDDVTSNME QIIKAKANLE KMCRTLEDQM NEHRSKAEET QRSVNDLTSQ  1276
SkMII |Q9UKX3|MYH13_HUMAN DLEEATLQHE ATAATLRKKQ ADSVAELGEQ IDNLQRVKQK LEKEKSELKM EIDDMASNIE ALSKSKSNIE RTCRTVEDQF SEIKAKDEQQ TQLIHDLNMQ  1280

SMII  |P35749|MYH11_HUMAN VHKLQNEVES VTGMLNEAEG KAIKLAKDVA SLSSQLQDTQ ELLQEETRQK LNVSTKLRQL EEERNSLQDQ LDEEMEAKQN LERHISTLNI QLSDSKKKLQ  1381
NMIIA |P35579|MYH9_HUMAN  VTKLQVELDN VTGLLSQSDS KSSKLTKDFS ALESQLQDTQ ELLQEENRQK LSLSTKLKQV EDEKNSFREQ LEEEEEAKHN LEKQIATLHA QVADMKKKME  1374
NMIIB |P35580|MYH10_HUMAN ASKLQNELDN VSTLLEEAEK KGIKFAKDAA SLESQLQDTQ ELLQEETRQK LNLSSRIRQL EEEKNSLQEQ QEEEEEARKN LEKQVLALQS QLADTKKKVD  1381
CMII  |P12883|MYH7_HUMAN  RAKLQTENGE LSRQLDEKEA LISQLTRGKL TYTQQLEDLK RQLEEEVKAK NALAHALQSA RHDCDLLREQ YEEETEAKAE LQRVLSKANS EVAQWRTKYE  1376
SkMII |Q9UKX3|MYH13_HUMAN KARLQTQNGE LSHRVEEKES LISQLTKSKQ ALTQQLEELK RQMEEETKAK NAMAHALQSS RHDCDLLREQ YEEEQEAKAE LQRALSKANS EVAQWRTKYE  1380

SMII  |P35749|MYH11_HUMAN -DFASTVEAL EEGKKRFQKE IENLTQQYEE KAAAYDKLEK TKNRLQQELD DLVVDLDNQR QLVSNLEKKQ RKFDQLLAEE KNISSKYADE RDRAEAEARE  1480
NMIIA |P35579|MYH9_HUMAN  -DSVGCLETA EEVKRKLQKD LEGLSQRHEE KVAAYDKLEK TKTRLQQELD DLLVDLDHQR QSACNLEKKQ KKFDQLLAEE KTISAKYAEE RDRAEAEARE  1473
NMIIB |P35580|MYH10_HUMAN DD-LGTIESL EEAKKKLLKD AEALSQRLEE KALAYDKLEK TKNRLQQELD DLTVDLDHQR QVASNLEKKQ KKFDQLLAEE KSISARYAEE RDRAEAEARE  1480
CMII  |P12883|MYH7_HUMAN  TDAIQRTEEL EEAKKKLAQR LQEAEEAVEA VNAKCSSLEK TKHRLQNEIE DLMVDVERSN AAAAALDKKQ RNFDKILAEW KQKYEESQSE LESSQKEARS  1476
SkMII |Q9UKX3|MYH13_HUMAN TDAIQRTEEL EEAKKKLAQR LQEAEENTET ANSKCASLEK TKQRLQGEVE DLMRDLERSH TACATLDKKQ RNFDKVLAEW KQKLDESQAE LEAAQKESRS  1480

SMII  |P35749|MYH11_HUMAN KETKALSLAR ALEEALEAKE ELERTNKMLK AEMEDLVSSK DDVGKNVHEL EKSKRALETQ MEEMKTQLEE LEDELQATED AKLRLEVNMQ ALKGQFERDL  1580
NMIIA |P35579|MYH9_HUMAN  KETKALSLAR ALEEAMEQKA ELERLNKQFR TEMEDLMSSK DDVGKSVHEL EKSKRALEQQ VEEMKTQLEE LEDELQATED AKLRLEVNLQ AMKAQFERDL  1573
NMIIB |P35580|MYH10_HUMAN KETKALSLAR ALEEALEAKE EFERQNKQLR ADMEDLMSSK DDVGKNVHEL EKSKRALEQQ VEEMRTQLEE LEDELQATED AKLRLEVNMQ AMKAQFERDL  1580
CMII  |P12883|MYH7_HUMAN  LSTELFKLKN AYEESLEHLE TFKRENKNLQ EEISDLTEQL GSSGKTIHEL EKVRKQLEAE KMELQSALEE AEASLEHEEG KILRAQLEFN QIKAEIERKL  1576
SkMII |Q9UKX3|MYH13_HUMAN LSTELFKMRN AYEEVVDQLE TLRRENKNLQ EEISDLTEQI AETGKNLQEA EKTKKLVEQE KSDLQVALEE VEGSLEHEES KILRVQLELS QVKSELDRKV  1580

SMII  |P35749|MYH11_HUMAN QARDEQNEEK RRQLQRQLHE YETELEDERK QRALAAAAKK KLEGDLKDLE LQADSAIKGR EEAIKQLRKL QAQMKDFQRE LEDARASRDE IFATAKENEK  1680
NMIIA |P35579|MYH9_HUMAN  QGRDEQSEEK KKQLVRQVRE MEAELEDERK QRSMAVAARK KLEMDLKDLE AHIDSANKNR DEAIKQLRKL QAQMKDCMRE LDDTRASREE ILAQAKENEK  1673
NMIIB |P35580|MYH10_HUMAN QTRDEQNEEK KRLLIKQVRE LEAELEDERK QRALAVASKK KMEIDLKDLE AQIEAANKAR DEVIKQLRKL QAQMKDYQRE LEEARASRDE IFAQSKESEK  1680
CMII  |P12883|MYH7_HUMAN  AEKDEEMEQA KRNHLRVVDS LQTSLDAETR SRNEALRVKK KMEGDLNEME IQLSHANRMA AEAQKQVKSL QSLLKDTQIQ LDDAVRANDD LKENIAIVER  1676
SkMII |Q9UKX3|MYH13_HUMAN IEKDEEIEQL KRNSQRAAEA LQSVLDAEIR SRNDALRLKK KMEGDLNEME IQLGHSNRQM AETQKHLRTV QGQLKDSQLH LDDALRSNED LKEQLAIVER  1680

SMII  |P35749|MYH11_HUMAN KAKSLEADLM QLQEDLAAAE RARKQADLEK EELAEELASS LSGRNALQ-D EKRRLEARIA QLEEELEEEQ GNMEAMSDRV RKATQQAEQL SNELATERST  1779
NMIIA |P35579|MYH9_HUMAN  KLKSMEAEMI QLQEELAAAE RAKRQAQQER DELADEIANS -SGKGALALE EKRRLEARIA QLEEELEEEQ GNTELINDRL KKANLQIDQI NTDLNLERSH  1772
NMIIB |P35580|MYH10_HUMAN KLKSLEAEIL QLQEELASSE RARRHAEQER DELADEITNS ASGKSAL-LD EKRRLEARIA QLEEELEEEQ SNMELLNDRF RKTTLQVDTL NAELAAERSA  1779
CMII  |P12883|MYH7_HUMAN  RNNLLQAELE ELRAVVEQTE RSRKLAEQEL IETSERVQLL HSQNTSL-IN QKKKMDADLS QLQTEVEEAV QECRNAEEKA KKAITDAAMM AEELKKEQDT  1775
SkMII |Q9UKX3|MYH13_HUMAN RNGLLLEELE EMKVALEQTE RTRRLSEQEL LDASDRVQLL HSQNTSL-IN TKKKLEADIA QCQAEVENSI QESRNAEEKA KKAITDAAMM AEELKKEQDT  1779

SMII  |P35749|MYH11_HUMAN AQKNESARQQ LERQNKELRS KLHEMEGAVK SKFKSTIAAL EAKIAQLEEQ VEQEAREKQA ATKSLKQKDK KLKEILLQVE DERKMAEQYK EQAEKGNARV  1879
NMIIA |P35579|MYH9_HUMAN  AQKNENARQQ LERQNKELKV KLQEMEGTVK SKYKASITAL EAKIAQLEEQ LDNETKERQA ACKQVRRTEK KLKDVLLQVD DERRNAEQYK DQADKASTRL  1872
NMIIB |P35580|MYH10_HUMAN AQKSDNARQQ LERQNKELKA KLQELEGAVK SKFKATISAL EAKIGQLEEQ LEQEAKERAA ANKLVRRTEK KLKEIFMQVE DERRHADQYK EQMEKANARM  1879
CMII  |P12883|MYH7_HUMAN  SAHLERMKKN MEQTIKDLQH RLDEAEQIAL KGGKKQLQKL EARVRELENE LEAEQKRNAE SVKGMRKSER RIKELTYQTE EDRKNLLRLQ DLVDKLQLKV  1875
SkMII |Q9UKX3|MYH13_HUMAN SAHLERMKKN LEQTVKDLQH RLDEAEQLAL KGGKKQIQKL ENRVRELENE LDVEQKRGAE ALKGAHKYER KVKEMTYQAE EDHKNILRLQ DLVDKLQAKV  1879
```

**Figure S5. Sequence alignment of human SmII, NMIIA, NMIIB, CMII and SkMII.** Residues that differ from consensus are identified with a dark background. Residues involved in binding to MT-228 are outlined in green. A red box outlines the Leu476 and its homolog Phe in other class II myosins.

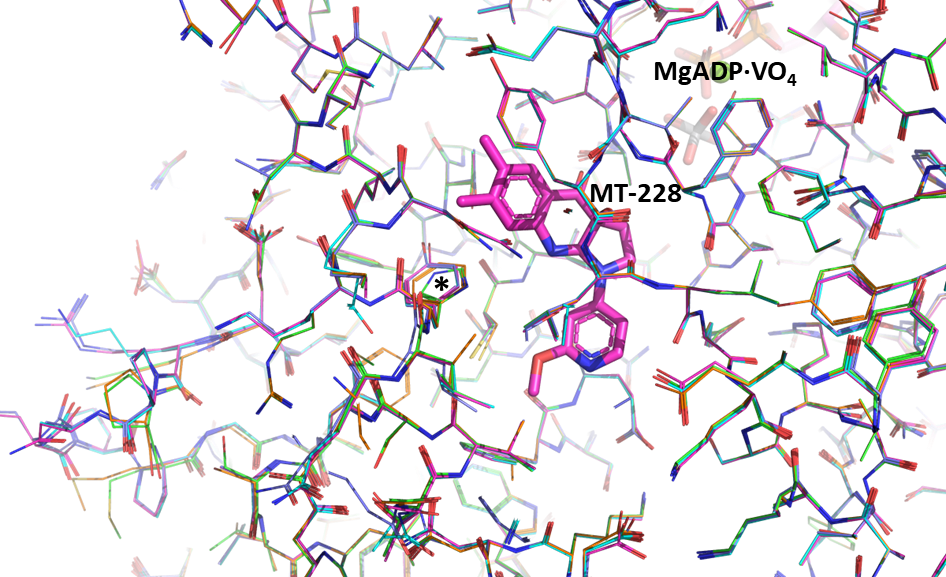

**Figure S6. Homology models of other class II myosins superimposed to the SmMII·MT-228 structure.** By color of the carbon atoms: pink: SmMII·MT-228 crystal structure; green: CMII; orange: SkMII; light blue: NMIIA; dark blue: NMIIB. The star outlines the main sequence difference in the interface with MT-228. This figure was prepared with PyMol 2.5.0.

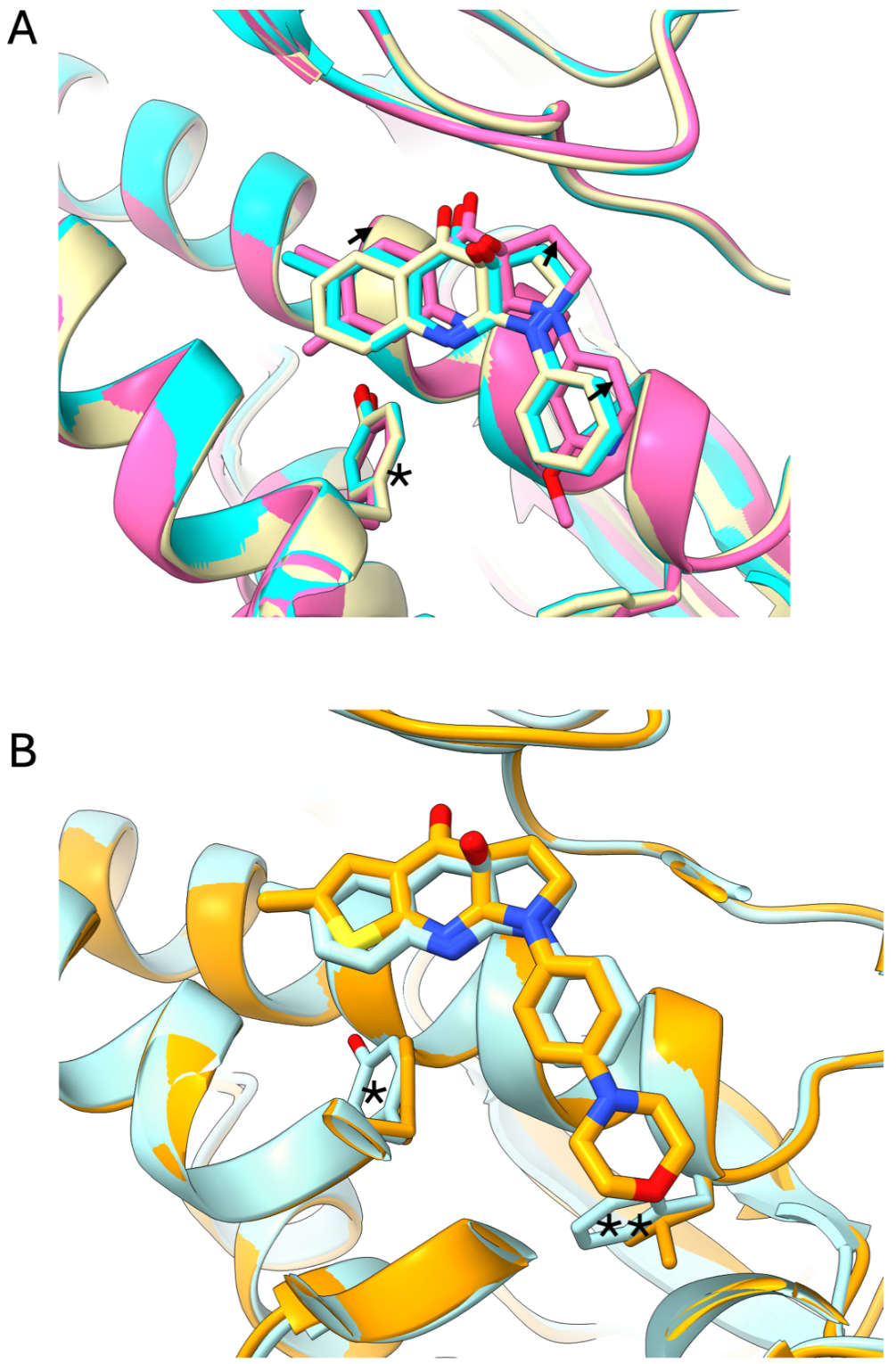

**Figure S7. MT-140 and MT-228 show different structural mechanisms of specificity.** Superimposition of DictyM2·MT-140 (beige carbons, PDB 3BZ9), DictyM2·Blebb (cyan carbons, PDB 1YV3), and SmMII·MT-228 (pink carbons). The shift in compound positioning is outlined by arrows. Note that the two derivatives are displaced in the opposite direction relative to Blebb.

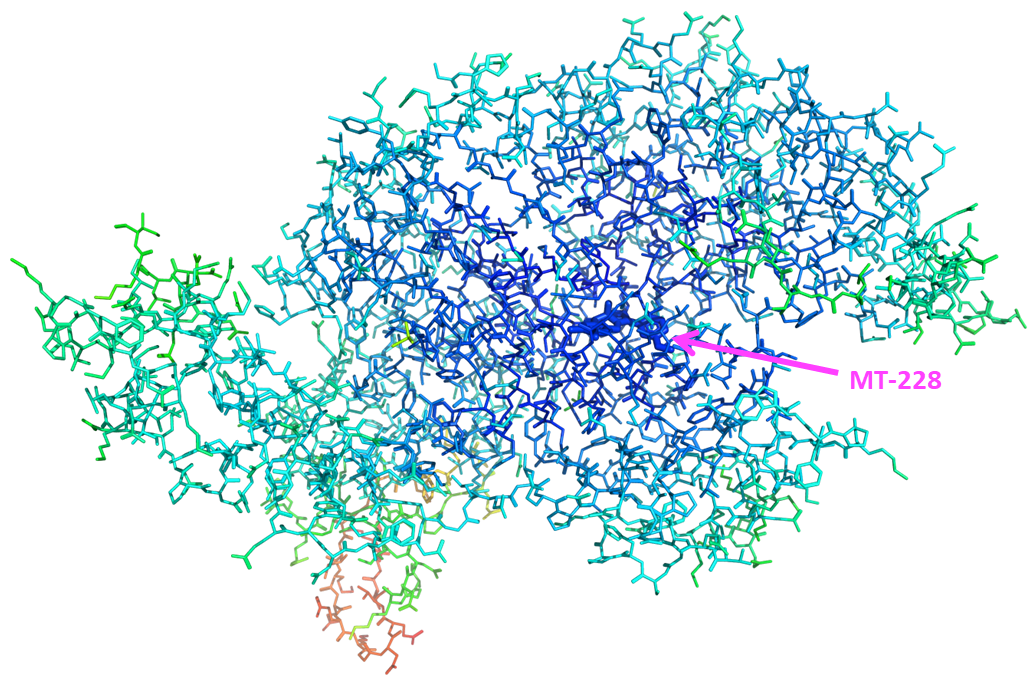

**Figure S8. structure of the SmII·MT-228 complex colored by B-factors**. MT-228 bonds are represented by thicker lines. B-factors calculated by Buster are pictured as a rainbow from dark blue (30 Å^2^) to red (212 Å^2^). This figure was prepared with PyMol 2.5.0.

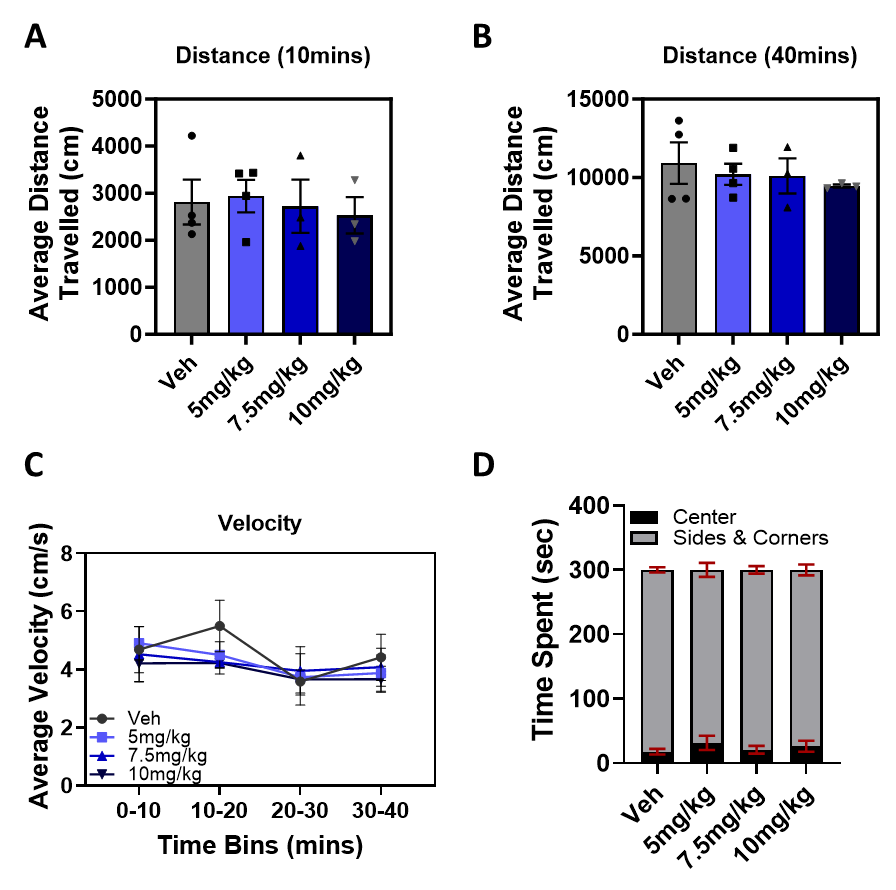

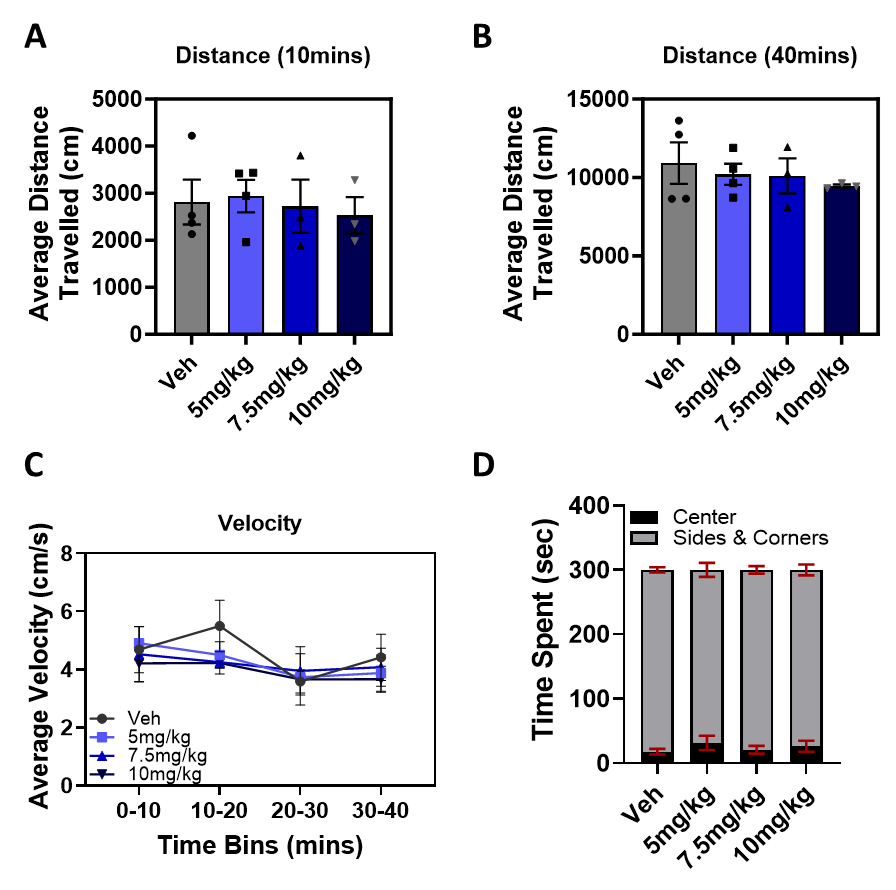

**Figure S9. MT-228 has no effect on general tolerability or locomotion.** Mice were dosed with Veh, 5, 7.5, or 10 mg/kg and monitored in an Open Field for 40 min. There was no significant difference in the distance moved within the A) first 10 min or B) over the entire 40 min test period. C) Similarly, there were no significant differences in animals’ velocity across the 40 min test period. D) MT-228 did not alter time spent in the center versus sides and corners at any dose tested, indicating that the compound did not alter this measure of anxiety-like behavior. Error bars represent SEM, * *P* < 0.05.

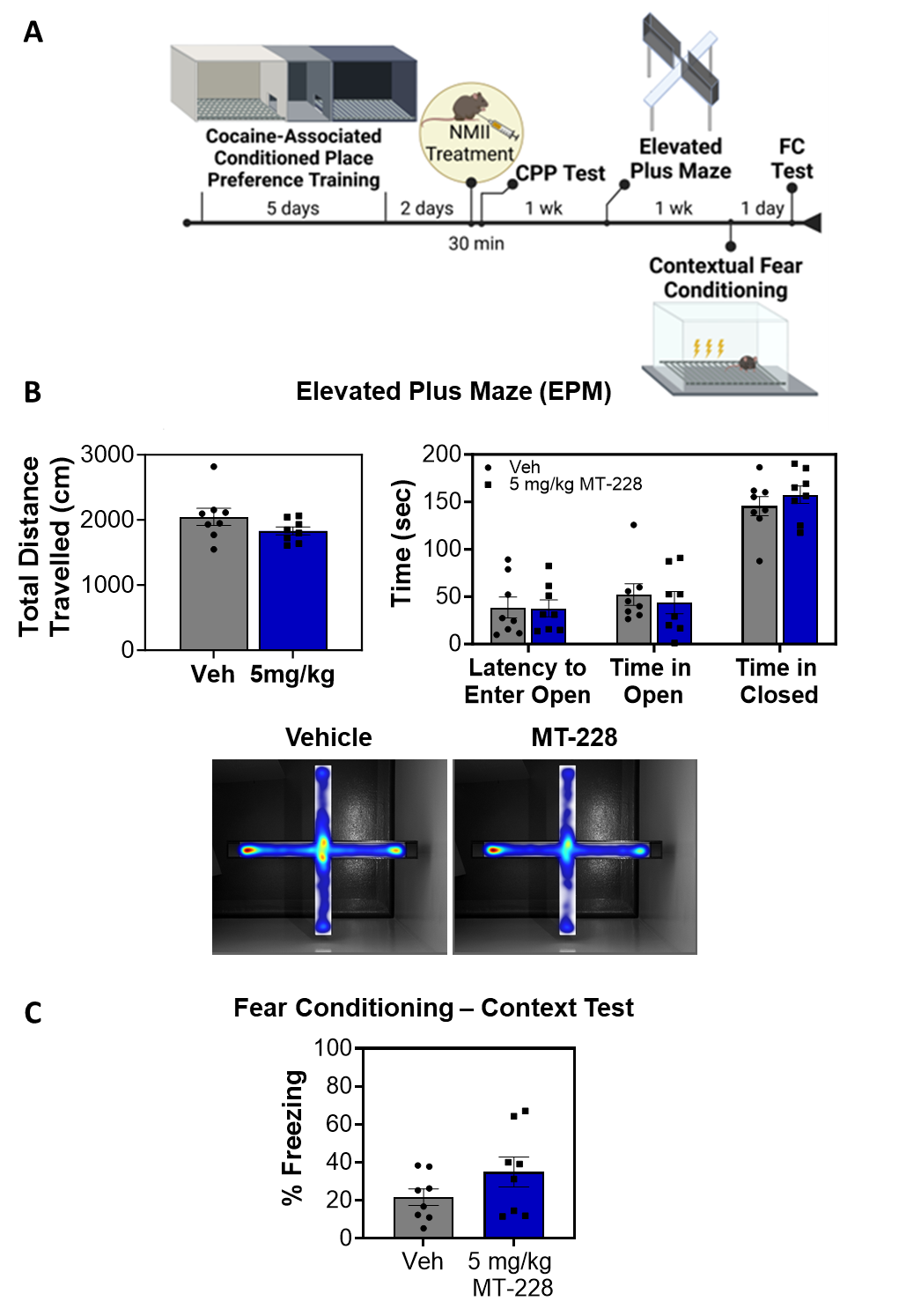

**Figure S10. Previous treatment with MT-228 had no effect on subsequent anxiety-like behavior or fear learning.** A) Experimental timeline of behavior battery. B) One week after COC CPP testing, mice were tested for anxiety-like behavior in an elevated plus maze. Previous treatment had no effect on distance travelled, latency to enter open arm or time spent in either the open or closed arm. C) One week later, animals underwent contextual fear conditioning. Similarly, previous treatment with MT-228 had no effect on animals’ ability to form contextual fear memories. Error bars represent SEM, * *P* < 0.05.

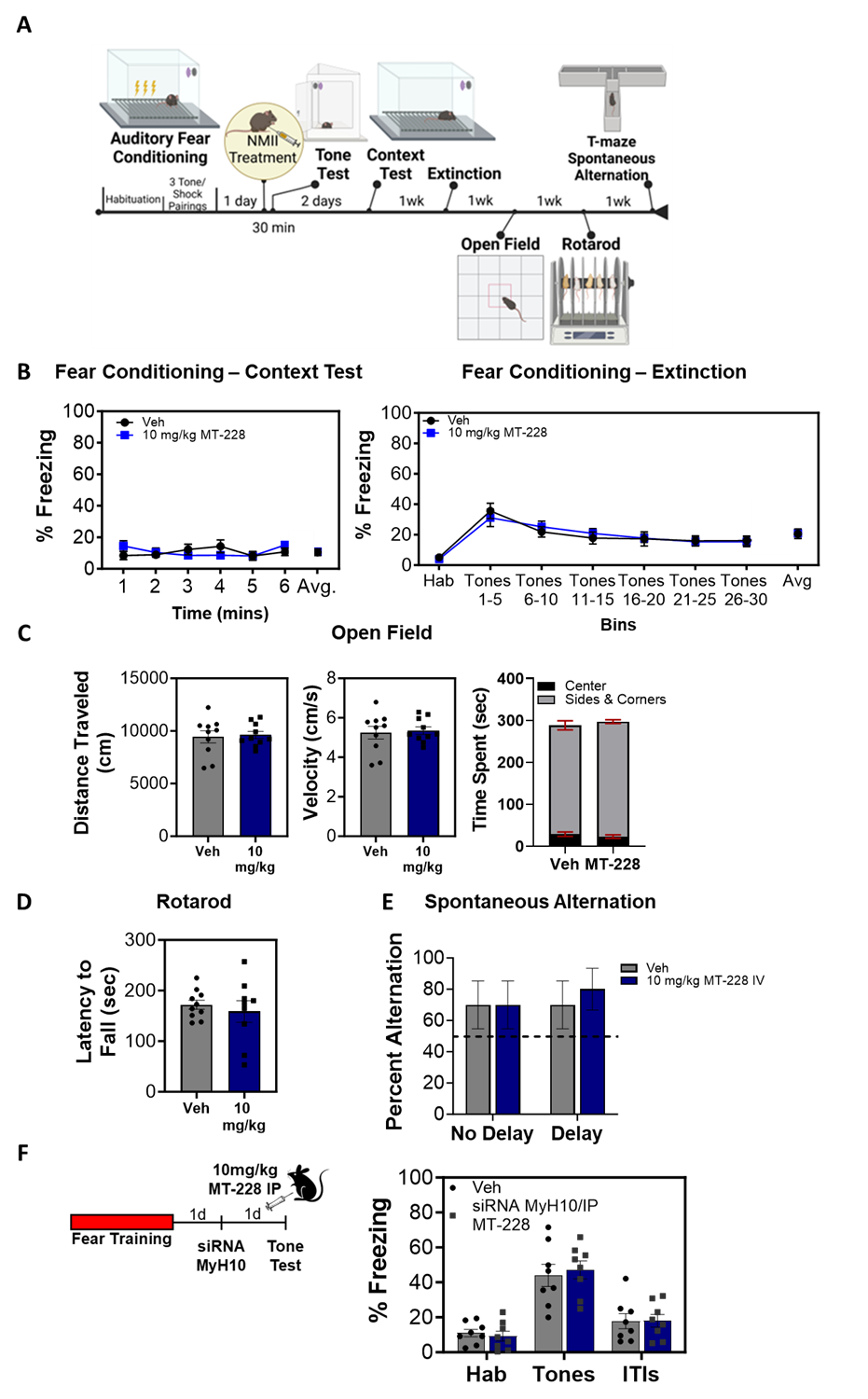

**Figure S11. Treatment with MT-228 does not influence fear memory or extinction, locomotion, or short-term memory.** A) Experimental design of the second behavior battery. B) Following the auditory tone test, mice underwent a contextual fear memory test, then extinction training. Prior treatment with MT-228 had no effect on either behavior. Similarly, there were no long-term changes to locomotion measured in the C) open field or D) on the rotarod. E) Spontaneous alternation in a T-maze was also unaffected by prior exposure to MT-228. F) In a separate group of mice, 10 mg/kg MT-228 (IP) combined with targeted delivery of an siRNA against *Myh10* to the BLA after training, but prior to testing to produce a complete blockade of NMII function had no effect on expression of an auditory fear memory. Error bars represent SEM, * *P* < 0.05.

### Accession codes

Numbering of alignments:

hSmII [P35749](https://www.uniprot.org/uniprotkb/P35749/entry#sequences)

NMIIA [P35579](https://www.uniprot.org/uniprotkb/P35579/entry#sequences)

NMIIB [P35580](https://www.uniprot.org/uniprotkb/P35580/entry#sequences)

CMII [P12883](https://www.uniprot.org/uniprotkb/P12883/entry#sequences)

SkMII [Q9UKX3](https://www.uniprot.org/uniprotkb/Q9UKX3/entry#sequences)

### Software versions

Autoproc 12.6.7

Buster 2.10.4

Coot 0.9.8.93

CCP4 8.0.013

XDS 20220820

PyMOL 2.5.0 Open-Source

ChimeraX 1.7.1

Cresset Flare 8.0.0
